# Engineering ionizable lipids for rapid biodegradation balances mRNA vaccine efficacy and tolerability

**DOI:** 10.1101/2024.08.02.606386

**Authors:** Julien Couture-Senécal, Grayson Tilstra, Omar F. Khan

## Abstract

The optimization of lipid nanoparticles (LNPs) has played a key role in enhancing the efficacy of mRNA vaccines, yet challenges with LNP tolerability remain. The ionizable lipid component within LNPs is critical to the efficient delivery of mRNA. Ionizable lipids can also trigger innate immune activation, which is beneficial for vaccine efficacy but may contribute to adverse inflammatory reactions. Engineering ionizable lipids for rapid biodegradation is a promising, yet underexplored, strategy to dampen inflammation. Here, we report the rational design and optimization of biodegradable ionizable lipids for intramuscular mRNA vaccines in mice. We show that in vivo biodegradability is enhanced by controlling lipid hydrolysis kinetics and that protein output is maximized by tuning the LNP apparent pK_a_. In an influenza vaccine model, the lead lipid (δO3) generates equivalent neutralizing antibodies and stronger antigen-specific T cell responses compared to a benchmark lipid (SM-102) used in approved mRNA vaccines. Furthermore, by comparing ionizable lipid analogs with similar potency but opposing biodegradation kinetics, we find that faster lipid clearance from tissues coincides with a lower inflammatory response while preserving strong vaccine immunogenicity. These findings demonstrate that fast-degrading ionizable lipids can balance the efficacy and tolerability of mRNA vaccines, with implications for addressing side effects and patient acceptance of new vaccine applications.

## Introduction

Messenger ribonucleic acid (mRNA) vaccines offer a promising alternative to conventional vaccines and are projected to become a dominant vaccine platform.^1,2^ Vaccine efficacy depends on the efficient delivery of mRNA to cells. The most advanced mRNA vaccines use lipid nanoparticles (LNPs) as a delivery system for mRNA. LNPs encapsulate mRNA to protect it from enzymatic degradation, are readily internalized by innate immune cells via endocytosis, and facilitate mRNA delivery to the cytosol where translation occurs.^3^ LNPs used in approved mRNA vaccines contain four lipid components: an ionizable lipid, cholesterol, a phospholipid, and a polyethylene glycol (PEG) lipid.^4–6^ The chemical structure of the ionizable lipid is critical to the delivery efficiency of mRNA, which directly impacts protein output and vaccine efficacy.^7–10^ Ionizable lipids are carefully designed such that LNPs have a neutral surface charge at physiological pH and then become positively charged in acidic conditions, facilitating both mRNA encapsulation and endosomal escape. While LNPs are currently the most potent and well-tolerated delivery system for mRNA, new ionizable lipid chemistry may further improve the efficacy and tolerability of mRNA vaccines.^11–13^

Unlike conventional vaccines such as inactivated viruses or protein subunits, mRNA vaccines do not require the co-administration of adjuvants. Both mRNA and the ionizable lipid possess intrinsic immunostimulatory properties, though their relative contributions to the adjuvant activity remain poorly understood.^14^ The innate immune sensing of mRNA by pattern recognition receptors is well-studied and can be alleviated via nucleoside modifications and removal of double-stranded RNA (dsRNA) impurities.^15,16^ In contrast, the mechanisms through which ionizable lipids are sensed by the innate immune system have yet to be elucidated. Recent reports have revealed that ionizable lipids are highly inflammatory compounds that may trigger innate immune activation through endosomal damage.^17–20^ Innate immune activation is beneficial to vaccine efficacy but can also be detrimental to the safety profile, requiring careful balance. Excessive innate immune activation induced by ionizable lipids in mRNA vaccines may be responsible for inflammation-related side effects observed in patients such as pain, swelling, fever, and rare cases of myocarditis.^21–23^ The higher reactogenicity profile of current mRNA vaccines compared to conventional vaccines has led to vaccine hesitancy and negatively impacted public health strategies, highlighting the need for safer and more effective formulations.^24^

Designing ionizable lipids that undergo rapid biodegradation is a promising yet underexplored strategy to dampen excessive innate immune activation. To enhance metabolism and elimination, biodegradable linkers, typically ester groups, have been successfully incorporated into ionizable lipids without compromising potency.^25,26^ However, the biodegradation kinetics of ester groups can vary drastically based on their susceptibility to enzymatic hydrolysis. The ionizable lipids used in approved mRNA vaccines contain sterically hindered esters which are slowly metabolized over the course of several days, presenting an opportunity to design ionizable lipids that clear more rapidly from the body.^27^

To directly examine the effect of ionizable lipid biodegradability on mRNA vaccine responses, we designed ionizable lipid analogs with similar potency but opposing biodegradation kinetics. This was achieved through the rational design and optimization of a novel series of biodegradable ionizable lipids for intramuscular mRNA vaccines in mice. We found that the faster clearance of ionizable lipids lowered the inflammatory response while preserving strong vaccine immunogenicity. We also established clear structure-activity relationships regarding the pK_a_ and hydrolysis kinetics of ionizable lipids to guide the design of more potent and better-tolerated LNPs for intramuscular mRNA vaccines. Ultimately, improving ionizable lipid chemistry may alleviate the reactogenicity associated with mRNA vaccines and improve their efficacy against infectious diseases beyond COVID-19.

## Results and discussion

### Rational design of biodegradable ionizable lipids with tunable pK_a_

We began this study by designing a novel series of biodegradable ionizable lipids (Figure 1A,B). Our group previously optimized an ionizable lipid, herein referred to as βN2, for intramuscular mRNA delivery (Figures S1– S3).^8^ βN2 contains three molecular regions: an amine core, amide linkers, and hydrocarbon tails (Figure 1A). Hydrocarbon tails control the lipid self-assembly with the mRNA payload, the number of carbons in each linker fine-tunes the lipid pK_a_, and the ethanolamine core enhances delivery efficiency. The linkers within βN2 contain amides, known for their chemical stability. Since esters are more susceptible to enzymatic hydrolysis than amides, we reasoned that replacing the amides of βN2 with esters, and positioning them to avoid steric hindrance, would result in faster biodegradation and clearance from the body.^28–30^ However, the amide-to-ester substitution presents additional design challenges to maintain LNP quality attributes and high delivery efficiency. Esters are much stronger electron-withdrawing groups than amides and lack the ability to donate hydrogen bonds. We predicted using cheminformatics that the direct ester analog of βN2, called βO2, would have a lower pK_a_ and a higher lipophilicity (logP), both of which contribute to reducing the LNP apparent pK_a_ (Figure 1C and Figure S4).^31^ The apparent pK_a_ roughly indicates the pH at which LNPs acquire a positive surface charge and must be precisely adjusted between 6 and 7. LNPs with an apparent pK_a_ outside of this range are typically toxic or exhibit low delivery efficiency.^7,8,32^ It is generally believed that the pK_a_ directly influences the efficiency of endosomal escape, but direct evidence to support this hypothesis is lacking. To precisely adjust apparent pK_a_, we synthesized six additional ester analogs with various carbon linkers (Figure 1B and Supplementary Methods). Each ionizable lipid contains the same amine core and hydrocarbon tails as βO2, but the length of the carbon linkers between the ester group and tertiary amines was increased systematically to tune the pK_a_ (Figures S5– S25).

**Figure 1.**
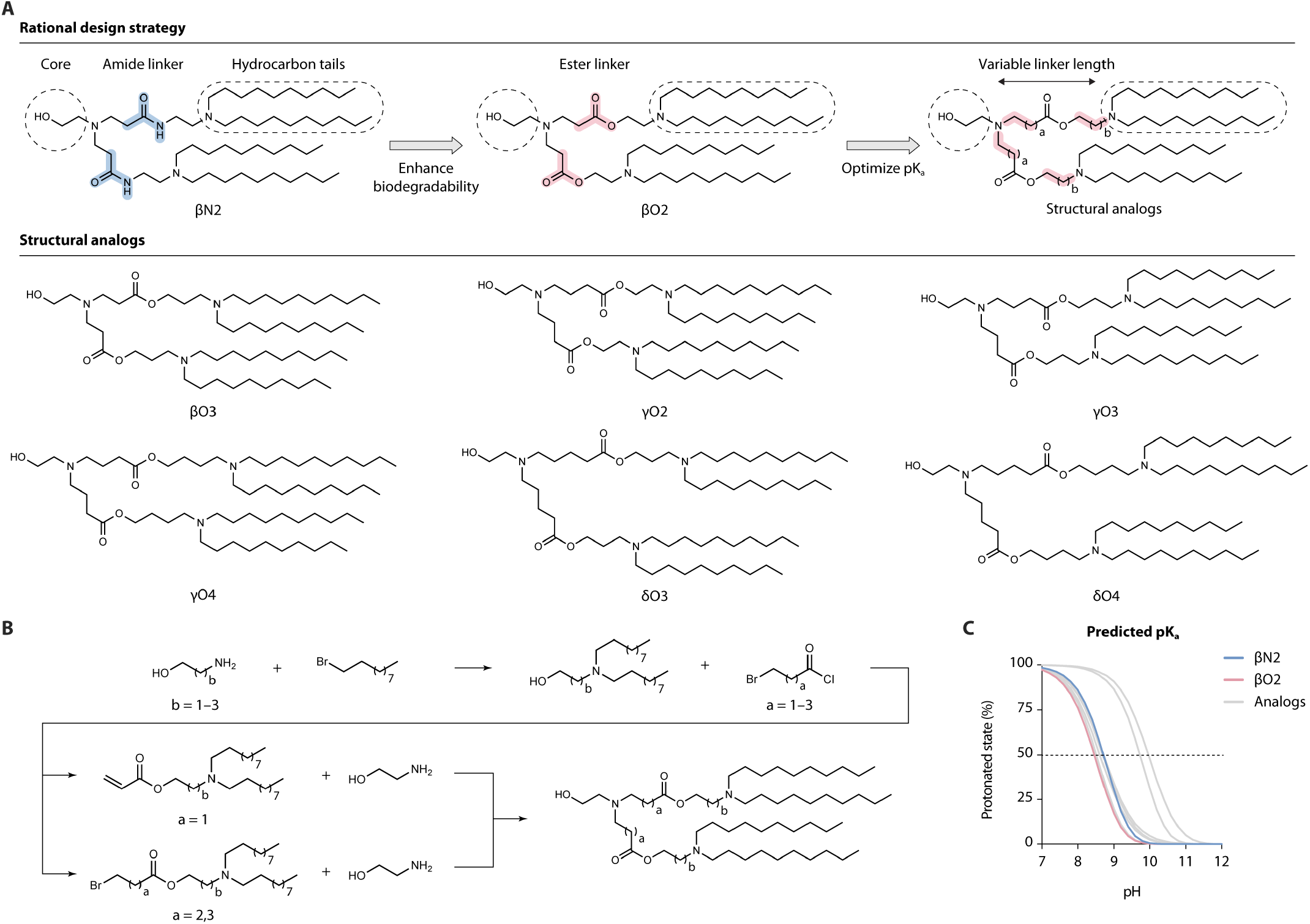
Rational design of biodegradable ionizable lipids with tunable pK_a_. **A**, Rational design strategy and chemical structures of ionizable lipids synthesized in this study. The nomenclature indicates the number of carbons on the carbonyl side (Greek letter) and the number of carbons on the alcohol side (numeral) of the linker. **B**, Synthesis pathways of a novel series of biodegradable ionizable lipids. **C**, Predicted pH-dependent ionization behaviour of ionizable lipids. The pK_a_ corresponds to the pH at 50% protonated state.

### Tuning LNP apparent pK_a_ maximizes intramuscular mRNA delivery

To identify a lead ester analog, we characterized each ionizable lipid’s ability to form high quality particles and generate protein output after intramuscular injection.^33^

The lipid composition of LNPs used in this study was modeled after mRNA-1273, the Moderna COVID-19 mRNA vaccine (Figure 2A).^5^ Each mRNA-LNP formulation was identical except for the ionizable lipid component. DLin-MC3-DMA (MC3), ALC-0315, and SM-102 were used as benchmark ionizable lipids as they are gold standards for RNA delivery.^4–6,34^ MC3 is a component of Patisiran, ALC-0315 is a component of BNT162b2, and SM-102 is a component of mRNA-1273 and mRNA-1345 (Figures S26–S28). Firefly luciferase mRNA was used as the payload to provide reliable measurements of protein expression through in vivo luminescence imaging (Table S1). Each ionizable lipid was evaluated based on its properties once formulated into LNPs (size distribution, zeta potential, mRNA encapsulation efficiency, and apparent pK_a_) (Figure 2B). To screen ionizable lipids for intramuscular expression, BALB/c mice were injected intramuscularly with 0.5 µg of firefly luciferase (FLuc) mRNA-LNPs in both quadriceps, and the total luminescent flux was measured at 6, 24, and 48 hours (Figure 2C).

**Figure 2.**
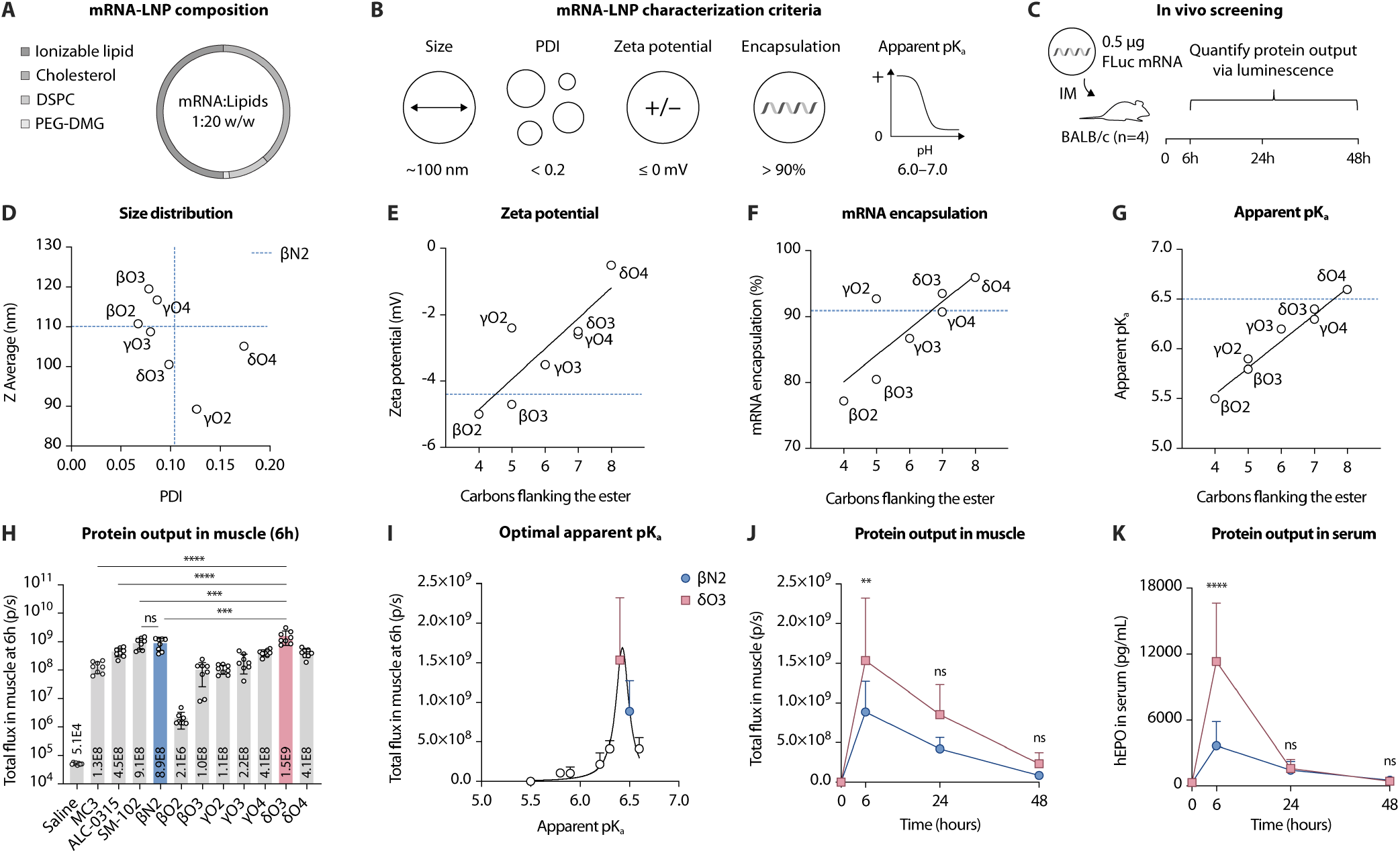
Optimization of biodegradable ionizable lipids for intramuscular mRNA delivery. **A**, mRNA-LNP composition. The lipid molar ratios were fixed at ionizable lipid:cholesterol:DSPC:DMG-PEG 2000 = 50:38.5:10:1.5 mol%. The mass ratio of mRNA:lipids was fixed at 1:20 w/w. **B**, mRNA-LNP characterization criteria. Z average size should be about 100 nm, polydispersity index (PDI) should be lower than 0.2, zeta potential should have a slight negative charge, mRNA encapsulation efficiency should be greater than 90%, and apparent pK_a_ should be between 6 and 7. **C**, Graphical summary of the in vivo screening method. BALB/c mice (n = 4) were injected intramuscularly with 0.5 µg of firefly luciferase (FLuc) mRNA-LNPs in both quadriceps. The total luminescent flux in injected hindlimbs was detected at 6, 24, and 48 hours as a correlate of protein output. **D–G**, Physicochemical characterization of FLuc mRNA-LNPs. Each data point represents an LNP formulation containing a different ester analog (n = 3 technical replicates). Blue dotted lines indicate the results for βN2. **D**, Size distribution of LNPs displayed as the Z average size versus PDI. **E–G**. LNP characteristics as a function of linker length (number of carbons flanking the ester). Data were fitted to a linear regression. **E**, Average zeta potential (R^2^ = 0.71). **F**, mRNA encapsulation efficiency (R^2^ = 0.66). **G**, Apparent pK_a_ (R^2^ = 0.97). **H**, Total luminescent flux in injected hindlimbs of BALB/c mice (n = 4 mice, 8 hindlimbs) at 6 hours after intramuscular injection with FLuc mRNA-LNPs containing different ionizable lipids. **I**, Relationship between the total luminescent flux at 6 hours (Figure H) and LNP apparent pK_a_ (Figure 2G) fitted to a Lorentzian distribution (R^2^ = 0.71). **J**, Total luminescent flux in injected muscle of BALB/c mice (n = 4 mice, 8 hindlimbs) after intramuscular injection with FLuc mRNA-LNPs containing βN2 or δO3. **K**, Serum concentration of human erythropoietin (hEPO) in BALB/c mice (n = 4 mice, terminal timepoints) after intramuscular injection with 0.5 µg hEPO mRNA-LNPs containing βN2 or δO3. Data for βN2 are indicated by blue bars or circles, and data for δO3 are indicated by red bars or squares. Data and error bars indicate the mean and standard deviation. P-values were determined by (**H**) one-way or (**J**,**K**) two-way ANOVA with Bonferroni’s multiple comparisons test.

We found that the direct ester analog of βN2 (βO2) formed LNPs with similar size and charge as βN2, with an identical Z-average size (110 nm), a low polydispersity index (PDI < 0.1), and a small negative average zeta potential (between -5 and -4 mV) (Figure 2D,E and Table S2). However, βO2 was worse at encapsulating mRNA than βN2, with an efficiency of 77% compared to 91% (Figure 2F). As predicted, the apparent pK_a_ was lower for βO2 than for βN2, with a value of 5.5 compared to 6.5 for βN2 (Figure 2G). All other ester analogs formed LNPs with a Z-average size between 89 and 120 nm, with a monomodal size distribution and a PDI below 0.2 (Figure 2D). There was no clear relationship between the ionizable lipid structure and size distribution. Most ester analogs formed stable LNPs after one freeze-thaw cycle, except for δO4 which had the highest PDI before freezing (Figure S29 and Table S3). We established positive correlations between the carbon linker length and zeta potential (Figure 2E), encapsulation efficiency (Figure 2F), and apparent pK_a_ (Figure 2G). Increasing the linker length of βO2 increased zeta potential from -5 mV to -0.5 mV (R^2^ = 0.71), mRNA encapsulation efficiency from 77% to 96% (R^2^ = 0.66), and apparent pK_a_ from 5.5 to 6.6 (R^2^ = 0.97). These results suggest that ionizable lipids with higher pK_a_ are better at complexing mRNA during the formulation process, resulting in fewer mRNA molecules close to the LNP surface and in a less negative surface charge (Figure S30).

The highest luminescent flux in hindlimbs was measured at 6 hours post-injection for all LNPs and rapidly declined over the span of 48 hours (Figures S31 and S32). As previously reported, βN2 generated expression levels equivalent to SM-102 and higher than ALC-0315 and MC3 (Figure 2H and Table S4). In contrast, the flux generated by βO2 was nearly three orders of magnitude lower than the flux generated by βN2 at the 6-hour timepoint. All other analogs achieved comparable or better levels of expression as the benchmarks. As anticipated, LNPs with higher apparent pK_a_ improved the protein output. We established a non-linear relationship between apparent pK_a_ and luminescent flux, with an optimal apparent pK_a_ of 6.4 (Figure 2I). δO3 was the most effective ester analog for intramuscular mRNA delivery, generating a higher flux than βN2 (2x, P = 0.0003), SM-102 (2x, P = 0.0005), ALC-0315 (3x, P < 0.0001), and MC3 (11x, P < 0.0001) at 6 hours. We also screened these LNPs for expression in THP-1 monocytes and found a weak correlation with in vivo expression (R^2^ = 0.599) (Figure S33). To determine if the high efficacy of δO3 for intramuscular delivery was independent of the mRNA payload, we compared the ability of βN2 and δO3 to deliver human erythropoietin (hEPO) mRNA (Table S1). hEPO is a secreted protein that can be detected in serum using an enzyme-linked immunosorbent assay. BALB/c mice were injected intramuscularly with 0.5 µg of hEPO mRNA-LNPs containing δO3 or βN2. At the 6-hour timepoint, δO3 generated higher serum expression than βN2 (3x, P < 0.0001) (Figure 2J,K). Regardless of the mRNA payload, δO3 and βN2 efficiently encapsulated mRNA in LNPs with similar physicochemical properties (Figure S34), and δO3 was more effective at generating protein expression than βN2 after intramuscular administration of mRNA-LNPs.

### Faster lipid hydrolysis kinetics accelerate tissue clearance

δO3 was designed to degrade faster than βN2 since esters are typically more reactive to hydrolysis than amides. To quantify differences in the hydrolysis kinetics of βN2 and δO3, we monitored their reaction with potassium hydroxide in methanol using time-series ^1^H-NMR spectroscopy. The hydrolysis half-life of βN2 was about 300 times higher than that of δO3 (Figure 3A and Figure S35). Importantly, δO3 did not hydrolyze after 12-month storage in ethanol (Figure S36). To assess in vivo clearance kinetics, CD-1 mice were injected intramuscularly with 1 µg hEPO mRNA-LNPs containing βN2 or δO3, and the concentration of intact ionizable lipid in plasma, injected muscle, liver, and spleen were measured by liquid chromatography with tandem mass spectrometry (LC-MS/MS) at various timepoints up to 7 days (Figure 3B). Pooled urine and feces samples were also analyzed from each cage. δO3 lowered the overall exposure in all analyzed tissues compared to βN2, with significant differences in plasma and liver (Figure 3C). Although the estimated apparent half-life for βN2 in plasma (∼295 hours) could not be accurately determined, the half-life for δO3 was much shorter (15 hours) (Figure 3D). The total exposure for δO3 was 92% lower in plasma, 60% lower in muscle, 93% lower in liver, and 84% lower in the spleen than the exposure for βN2 (Figure C–G, Tables S5 and S6). Approximately 18% of βN2 was excreted in feces after 21 days, while only small and variable amounts of δO3 (between 0.1% and 3%) were detected in feces (Table S7). Neither compound was detected in urine. Together, these results demonstrate that βN2 was slowly eliminated through hepatobiliary pathways and suggest that δO3 was more rapidly metabolized and eliminated from organs and tissues.

**Figure 3.**
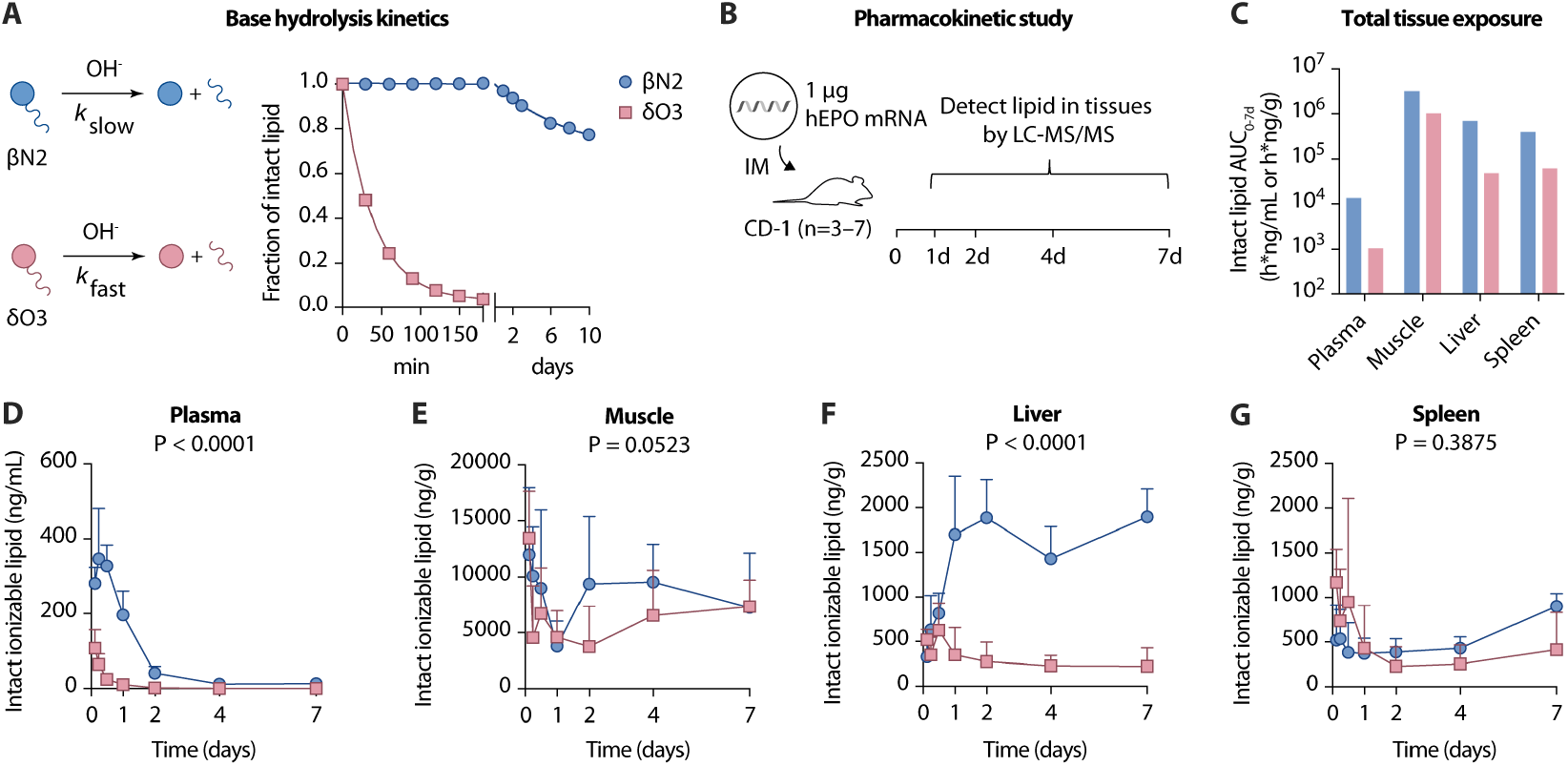
In vitro and in vivo biodegradability of βN2 and δO3. **A**, Base hydrolysis kinetics of βN2 and δO3. Ionizable lipids were reacted with potassium hydroxide in deuterated methanol. The fraction of intact lipid over time was monitored by time-series ^1^H-NMR and data were fitted to a one phase decay exponential function. **B**, Graphical summary of the pharmacokinetic study design. CD-1 mice were injected intramuscularly with 1 µg of hEPO mRNA-LNPs containing βN2 or δO3. Intact ionizable lipid was detected in tissues by LC/MS-MS at various timepoints up to 7 days. **C**, Total exposure (AUC_0–7d_) of βN2 and δO3 in plasma, injected muscle, liver, and spleen. **D–G**, Clearance kinetics of βN2 and δO3 in (**D**) plasma, (**E**) injected muscle, (**F**) liver, and (**G**) spleen. Data for βN2 are indicated by blue bars or circles, and data for δO3 are indicated by red bars or squares. Data and error bars indicate the mean and standard deviation across samples from independent mice (n = 3-7). P-values were determined by two-way ANOVA.

### Ionizable lipids drive systemic inflammation

To investigate the effect of ionizable lipid biodegradability on systemic inflammation, BALB/c mice were injected intramuscularly with 0.5 µg of hEPO mRNA-LNPs containing βN2 or δO3 (Figure 4A). The concentrations of several inflammatory cytokines in serum were measured by Luminex assay at 6, 24, and 48 hours after injection. Both LNPs elicited an acute inflammatory response characterized by an increase in IL-6, CSF3, CXCL1, CCL2, IL-5, and CXCL10 that returned to baseline by 48 hours (Figure 4B–H and Figure S37). IL-6 was the most upregulated cytokine, with mean concentrations 33x above baseline for βN2 (P < 0.0001) and 25x above baseline for δO3 (P < 0.0001). Compared to βN2, δO3 lowered the total expression of IL-6 by 32%, CSF3 by 60%, CXCL1 by 30%, CCL2 by 20%, and CXCL10 by 37% over 48 hours (Tables S8).

**Figure 4.**
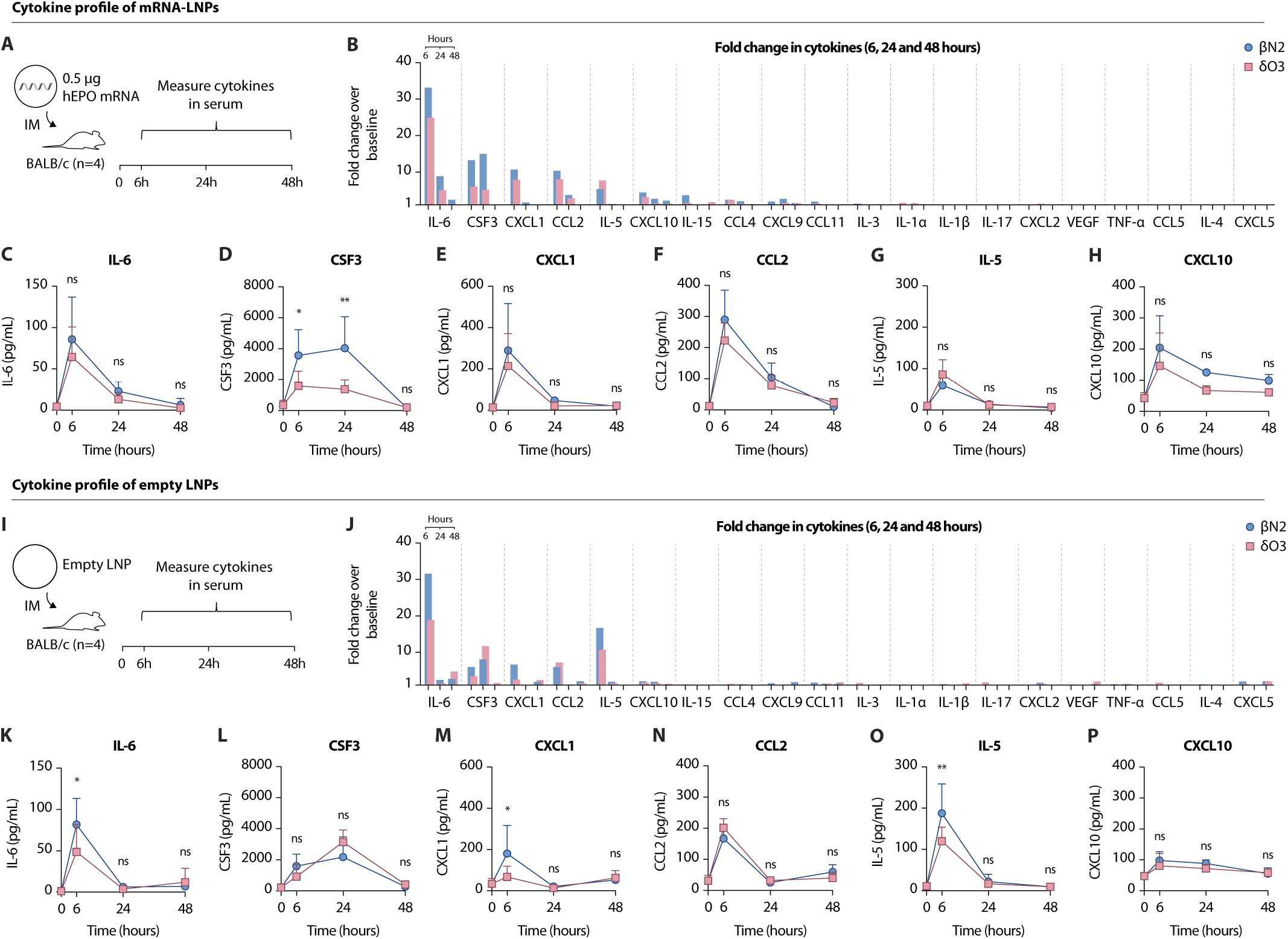
Cytokine profile of mRNA-LNPs and empty-LNPs. **A–H**, Cytokine profile of mRNA-LNPs. **A**, Graphical summary of the study design. BALB/c mice were injected intramuscularly with 0.5 µg of hEPO mRNA-LNPs containing βN2 or δO3. Cytokines were measured in serum at 6, 24, and 48 hours by Luminex assay. **B**, Fold change in serum cytokine concentrations of mice treated with mRNA-LNPs over untreated mice at 6, 24, and 48 hours. Data for βN2 and δO3 at each timepoint are superimposed. **C–H**, Kinetic profile of (**C**) IL-6, (**D**) CSF3, (**E**) CXCL1, (**F**) CCL2, (**G**) IL-5, and (**H**) CXCL10. **I–P**, Cytokine profile of empty-LNPs. **I**, Graphical summary of the study design. BALB/c mice were injected intramuscularly with a comparable dose of empty-LNPs containing βN2 or δO3. Cytokines were measured in serum at 6, 24, and 48 hours by Luminex assay. **J**, Fold change in serum cytokine concentrations of mice treated with empty-LNPs over untreated mice at 6, 24, and 48 hours. Data for βN2 and δO3 at each timepoint are superimposed. **K–P**, Kinetic profile of (**K**) IL-6, (**L**) CSF3, (**M**) CXCL1, (**N**) CCL2, (**O**) IL-5, and (**P**) CXCL10. Data for βN2 are indicated by blue bars or circles, and data for δO3 are indicated by red bars or squares. Data and error bars indicate the mean and standard deviation across samples from independent mice (n = 4, terminal timepoints). P-values were determined by two-way ANOVA with Bonferroni’s multiple comparisons test

Although the mRNA payload was chemically modified with N^1^-methylpseudouridine and purified to reduce dsRNA impurities (Figure S38), it may still be sensed by pattern recognition receptors and induce a cytokine response characterized by type 1 interferon signaling.^35,36^ To decouple the cytokines induced by ionizable lipids from those induced by mRNA, we injected BALB/c mice with empty LNPs containing βN2 or δO3 (Figure 4I). We observed that empty LNPs were smaller in size than mRNA-LNPs (Table S2). Like mRNA-LNPs, empty-LNPs induced a cytokine profile characterized by an increase in IL-6, CSF3, CXCL1, CCL2, IL-5, and CXCL10 at 6 hours that waned by 48 hours (Figure 4J– P and Figure S39). IL-6 was again the most upregulated cytokine, with mean concentrations 32x above baseline for βN2 (P < 0.0001) and 19x above baseline for δO3 (P < 0.0001). Compared to βN2, δO3 lowered the total expression of IL-6 by 32%, CXCL1 by 41%, and IL-5 by 32% over 48 hours, with significant differences at the 6-hour timepoint (Table S9). While differences in IL-6, CXCL1, and IL-5 between LNPs were more pronounced in the absence of an mRNA payload, this was not the case for CSF3, CCL2, and CXCL10, suggesting that the innate immune sensing of mRNA components might have contributed to the cytokine profile of mRNA-LNPs. Overall, the striking similarities between the cytokine profile of mRNA-LNPs and empty-LNPs demonstrate that ionizable lipids are the main driver of systemic inflammation. LNPs containing δO3 are also generally less inflammatory than LNPs containing βN2, which may result from differences in lipid biodegradability.

### Ionizable lipids influence cellular immunity

The inflammatory cytokine profile induced by LNPs contributes to reactogenicity but also directly influences the type and strength of vaccine responses. IL-6 has been shown to be a hallmark of the adjuvant activity of ionizable lipids in mice.^17^ In the context of mRNA vaccination, IL-6 polarizes T follicular helper (Tfh) cells, and CXCL10 polarizes T helper 1 (Th1) cells.^35–37^ CSF3, CXCL1, CCL2, and IL-5 may also contribute to the adjuvant activity of LNPs by regulating the recruitment and activation of various immune cells. To determine whether differences in the systemic cytokine profile induced by βN2 and δO3 would affect mRNA vaccine responses, we used an established influenza vaccine model in mice.^38–40^ LNPs containing βN2, δO3, or SM-102 were formulated with mRNA encoding the hemagglutinin (HA) protein from the H1N1 Influenza A/Puerto Rico/8/1934 virus (Tables S1 and S2). BALB/c mice were injected intramuscularly with 0.5 μg of HA mRNA-LNP on day 0 (prime) and day 30 (boost) (Figure 5A).

**Figure 5.**
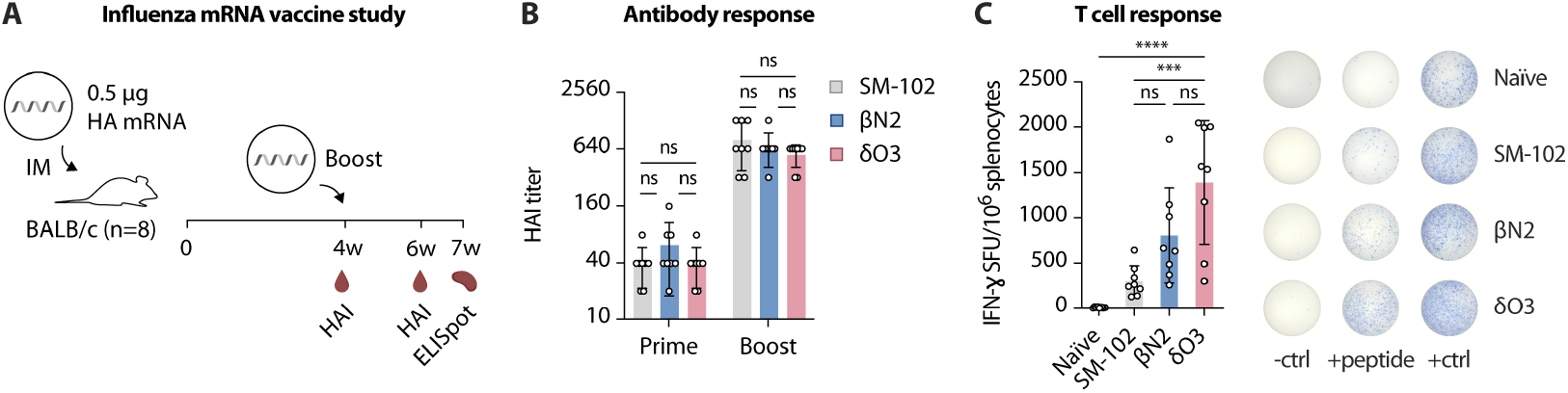
Immunogenicity of mRNA vaccines against influenza. **A**, Graphical summary of the influenza mRNA vaccine study. BALB/c mice were injected intramuscularly 30 days apart (prime and boost) with 0.5 µg HA mRNA-LNPs containing SM-102, βN2, or δO3. Serum was collected 4 weeks post-prime and 2 weeks post-boost to measure HAI titers. Splenocytes were harvested 3 weeks post-boost to measure HA-specific T cell responses by IFN-γ ELISpot. **B**, HAI titers after the prime and boost. **C**, IFN-γ ELISpot spot-forming units (SFU) per 10^6^ splenocytes after ex vivo stimulation with a peptide pool spanning the HA protein. Representative images (median SFU) of IFN-γ ELISpot assay wells for negative control (-ctrl), peptide stimulation (+peptide), and concanavalin A stimulation (+ctrl) groups. Data for SM-102 are indicated by gray bars, data for βN2 are indicated by blue bars, and data for δO3 are indicated by red bars. Data and error bars indicate the mean and standard deviation across samples from independent mice (n = 8). P-values were determined by one-way ANOVA with Bonferroni’s multiple comparisons test.

Hemagglutinin inhibition (HAI) titers in serum were measured 4 weeks after the prime (pre-boost) and 2 weeks after the boost. HAI titer is the major clinical correlate of protection against influenza as it is a simple readout of both the quality and quantity of neutralizing antibodies.^41^ An HAI titer of 40 was achieved after the prime dose for all LNPs (Figure 5B and Figure S40). After the boost dose, HAI titers increased by more than one order of magnitude, with titers as high as 1280 for some animals. All LNPs induced a similarly strong neutralizing antibody response. 3 weeks after the boost dose, HA-specific T cell responses were measured by IFN-γ T cell ELISpot assay by stimulating isolated splenocytes with an overlapping peptide pool spanning the HA protein. All LNPs induced HA-specific T cells, as determined by an increase in the average number of IFN-γ spot forming units (SFU) over naïve mice (Figure 5C). Surprisingly, both δO3 and βN2 induced higher HA-specific T cell responses than SM-102. δO3 resulted in slightly more SFU than βN2 (1.7x, P = 0.0762) and significantly more than SM-102 (4.6x, P = 0.0002) (Figure S41). Although δO3 induced a lower inflammatory cytokine profile than βN2, it generated comparable or better vaccine immunogenicity in an influenza model. These results show that the chemistry of ionizable lipids has a greater impact on cellular responses than on humoral responses in mice.

## Conclusion

In this study, we aimed to optimize a novel series of biodegradable ionizable lipids for intramuscular mRNA vaccines in mice. These lipids were designed by substituting the amide linker in βN2 with an ester linker to enhance biodegradability.^8^ By fine-tuning the linker length, we developed an ionizable lipid (δO3) which produced greater intramuscular expression than βN2 and industry benchmarks, making it one of the most potent ionizable lipids in the literature. This optimization process highlighted an direct relationship between the LNP apparent pK_a_ and intramuscular expression, building on previous studies.^7,8^

Furthermore, we examined whether enhancing the biodegradation kinetics of ionizable lipids could dampen inflammation. We demonstrated that δO3 was cleared more rapidly than βN2 from mice organs and tissues. While δO3 generated more protein output than βN2, it reduced the release of inflammatory cytokines such as IL-6. By combining these findings with those from the literature, we suggest that the prolonged retention of ionizable lipids within cells may contribute to innate immune activation (Figure 6).^7,14,20^ Therefore, the ideal ionizable lipid should be degraded immediately after delivering mRNA to the cytosol to minimize excessive inflammation linked to vaccine reactogenicity.

**Figure 6.**
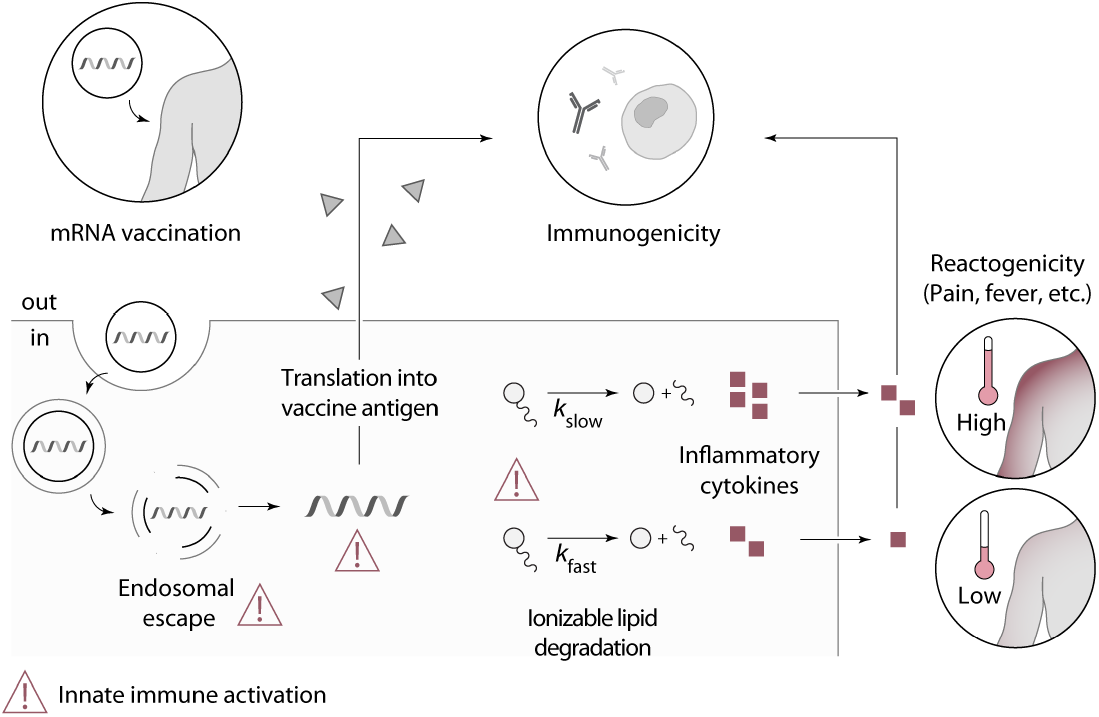
Proposed innate immune mechanism of mRNA vaccines. mRNA-LNPs injected intramuscularly are internalized by innate immune cells in muscle and draining lymph nodes.^42^ Following uptake, ionizable lipids become protonated in acidic endosomes, leading to the disruption of endosomal membranes, and allowing for the transport of mRNA into the cytosol where translation occurs. Vaccine antigens are then secreted by cells and processed for presentation on MHC molecules. Endosomal damage by ionizable lipids triggers the first activation signal. mRNA components elicit a second signal upon sensing by pattern recognition receptors. Intact ionizable lipids may trigger a third signal by disrupting intracellular membranes and activating innate immune sensors, which can be minimized by their rapid metabolism and elimination. These three innate immune activation signals lead to the production and release of inflammatory cytokines by innate immune cells, contributing to vaccine immunogenicity and reactogenicity.

Lastly, we evaluated the performance of δO3 and βN2 in an mRNA vaccine model against influenza. Both δO3 and βN2 elicited comparable vaccine immunogenicity based on HAI titers and HA-specific T cell responses. Interestingly, δO3 generated significantly higher T cell responses than SM-102. Although factors influencing cellular immunity are poorly understood, this result revealed that the development of new ionizable lipids is a promising strategy to improve the efficacy of T cell-inducing mRNA vaccines, which has implications for combatting rapidly mutating viruses like influenza.^43^

In conclusion, we developed a potent biodegradable ionizable lipid for intramuscular vaccine applications. We found that protein output can be maximized by tuning the LNP apparent pK_a_, that the inflammatory response to LNPs can be dampened by enhancing lipid biodegradation kinetics, and that T cell responses to mRNA vaccines is highly dependent on ionizable lipid chemistry. Together, these findings contribute to the development of new strategies to balance the efficacy and tolerability of mRNA vaccines, thereby harnessing the full potential of the vaccine platform to combat diseases.

## Methods

### Ionizable lipids

Detailed synthetic procedures and characterization data are presented in the Supplementary Information. Final ionizable lipids were characterized by proton (^1^H) and proton-decoupled carbon-13 (^13^C{^1^H}) nuclear magnetic resonance spectrometry (NMR), and electrospray ionization (ESI) mass spectrometry. The purity of δO3 and βN2 was >95% by mol as determined by ^1^H NMR.

Stock solutions of ionizable lipids were prepared by filtering a solution of the ionizable lipids in anhydrous, 200-proof ethanol through 0.2 μm polyethersulfone syringe filters, concentrating to a neat oil under high vacuum to determine exact mass, and then re-diluting in anhydrous, 200-proof ethanol to a final concentration between 50 to 100 mg/mL. Ionizable lipid stocks were stored in tightly sealed glass vials at -20°C, protected from light, and used for up to 6 months post-preparation. SM-102 (Cayman Chemicals, 33474), DLin-MC3-DMA (Cayman Chemicals, 34364), and ALC-0315 (Cayman Chemicals, 34337) were purchased from Cayman Chemicals (Ann Arbor, Michigan, USA). Marvin was used for predicting logP and pK_a_ (Marvin 23.11.0, Chemaxon). Electrospray ionization mass spectra in positive mode (ESI+) were acquired at the AIMS Mass Spectrometry Laboratory in the Department of Chemistry at the University of Toronto. NMR spectra were acquired at the CSICOMP NMR Facility in the Department of Chemistry at the University of Toronto (Toronto, Canada).

### mRNA synthesis

Custom linearized DNA templates (Genscript) containing a 101-nucleotide poly(A) tail were linearized with SapI or XbaI and transcribed using the HiScribe T7 mRNA Kit with CleanCap Reagent AG (New England Biolabs, E2080). In vitro transcription was performed with complete substitution of uridine with N^1^-methylpseudouridine (Trilink Biotechnologies, N-1081). All transcripts were purified by silica-column chromatography followed by cellulose purification to remove dsRNA impurities.^44^ The presence of dsRNA impurities was determined by dot blot (500 ng/dot) using J2 dsRNA-specific antibody (Novus Biologicals, NBP3-11395). The length and integrity of mRNA were validated on an Agilent TapeStation system (Agilent Technologies) at The Centre for Applied Genomics at The Hospital for Sick Children (Toronto, Canada). All mRNA sequences are provided in the Supplementary Information (Table S1).

### LNP formulation

Lipid components were combined at specific molar ratios (50 mol% ionizable lipid, 38.5 mol% cholesterol, 10 mol% DSPC, and 1.5 mol% DMG-PEG 2000) in ethanol at a final lipid concentration of 16 mg/mL.^8,45^ Ultrapure cholesterol (VWR, 97061-660) and DSPC (VWR, TCD3926) were purchased from VWR International (Radnor, Pennsylvania, USA), and DMG-PEG 2000 (Avanti Polar Lipids, 880151) was purchased from Avanti Polar Lipids (Alabaster, Alabama, USA). mRNA was diluted in 25 mM sodium acetate (pH 4.5) at 0.27 mg/mL. For empty LNPs, the mRNA was replaced with water. A typical formulation batch used 0.1 mg of mRNA and 2 mg of lipid components (mRNA:lipids = 1:20 w/w). The mRNA and lipid phases were rapidly injected into a PDMS herringbone micromixer at flow rates of 4.5 mL/min and 1.5 mL/min using a dual syringe pump. The mixture was directly collected (without handling) into a dialysis device with a molecular weight cutoff of 10 kDa (Thermo Scientific, 66383), and dialyzed twice against 1000-fold volume of 20 mM Tris-HCl pH 7.5 at 4°C. The first dialysis was performed for 4 hours, followed by a second dialysis overnight. The resulting LNP suspension was removed from the dialysis device and sterile filtered through a 0.2 μm polyethersulfone syringe filter (Cytiva, 99161302). The LNP solution was diluted with concentrated sterile sucrose to a final buffer composition of 20 mM Tris-HCl pH 7.5, 8% sucrose (Tris-sucrose buffer). LNP formulations in Tris-sucrose buffer were stored at -80°C in 1.5 mL polypropylene tubes for up to 1 month until use. LNPs were thawed overnight at 4°C the day before injections and diluted in Tris-sucrose buffer for injections. LNPs were characterized after sterile filtration.

### Nanoparticle size, zeta potential and concentration

A Zetasizer Ultra Red (Malvern Panalytical, Malvern, UK) was used to determine the nanoparticle size (Z average of the hydrodynamic diameter), the polydispersity index (PDI), zeta potential, and nanoparticle concentration. Measurements were done in triplicates for size and concentration. One measurement was done for zeta potential. LNPs were diluted at 2 ng/µL of mRNA or an equivalent volume of empty-LNPs in 1 mL of 20 mM Tris-HCl pH 7.5. The material refractive index was 1.45 and absorption was 0.001.

### mRNA concentration and encapsulation efficiency

The concentration and encapsulation efficiency of mRNA formulated in LNPs were determined using a modified Quant-iT RiboGreen assay (Thermo Fisher, R11491). The assay was performed in triplicates in a 96 well plate. 12 μL of the LNP sample or TE buffer (composed of 10 mM Tris-HCl containing 1 mM EDTA) (blank sample) was diluted in 300 μL of 1X TE buffer pH 7.5. 50 μL of the diluted samples were transferred to 6 separate wells and either 50 μL of TE buffer or 50 μL of a 4% Triton X-100 (Thermo Fisher, 85112) solution in TE buffer was added to the wells. Two 6-point standard curves (5 mg/mL to 0.156 g/mL) were prepared in the same plate by diluting mRNA to a final volume of 100 μl in 2% Triton X-100 or TE buffer. The plate was then incubated for 10 minutes at 40° C with gentle shaking (500 rpm) on a thermal mixer. The Ribogreen reagent was diluted 1:100 in TE buffer, and 100 μL of this solution was added to each well. 180 μL of this mixture was immediately transferred to an opaque, black 96-well plate. The plate was incubated for 5 minutes with gentle shaking away from light. Fluorescence intensity was measured with a plate reader at an excitation wavelength of 485 nm and an emission wavelength of 528 nm. The raw fluorescence values were subtracted by the background fluorescence (blank sample). The concentration of free mRNA and total mRNA were determined by linear interpolation of the appropriate standard curve. Encapsulation efficiency was determined by dividing the calculated concentration of encapsulated mRNA by the total mRNA concentration in the LNP sample. The mean of triplicates is reported.

### Apparent pK_a_

LNP apparent pK_a_ was determined using a 6-(p-toluidino)-2-naphthalenesulfonic acid (TNS) assay. A series of buffers was generated by titrating a solution containing 10 mM citrate, 10 mM phosphate, 10 mM borate, and 150 mM NaCl with 1.0 M HCl. 12 total buffers were prepared, ranging from pH 4.0 to 9.5 in increments of 0.5 pH. LNPs (diluted to 100 µM of ionizable lipid in distilled water) and TNS (Sigma Aldrich,T9792) (diluted to 60 µM in distilled water) were diluted into these buffers to a final concentration of 9 and 5.45 µM, respectively. Each LNP formulation was tested in triplicate in the same plate. The plate was covered and equilibrated at room temperature for 30 min. Fluorescence intensity was determined using a plate reader (325 nm excitation/435 nm emission). The raw fluorescence values were fitted to the following equation:

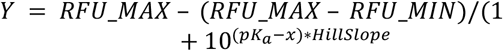

The apparent pK_a_ value corresponds to the pH value at 50% maximum fluorescence. The mean of replicates is reported.

### Animals

All procedures conducted at the University of Toronto were approved by Institutional Animal Care and Use Committee under protocol number 20012621 and comply with all applicable local, provincial, and federal regulations. Female BALB/c mice (7-8 weeks old, 17-20g) were purchased from Charles River Laboratories (Wilmington, Massachusetts, USA) and acclimated for 1 to 2 weeks. Mice were injected intramuscularly with 50 µL of LNP solution in the quadricep.

### Intramuscular detection of firefly luciferase luminescent flux in mice

BALB/c mice (n = 4) were injected in both quadriceps with 0.5 μg of firefly luciferase mRNA-LNPs. At various timepoints (6, 24, 48, and 96 hours post-dose), mice were injected intraperitoneally with 3 mg D-Luciferin (Thermo Fisher, 88291) in 200 μl 1X PBS. 5 minutes later, mice were anesthetized and imaged on an in vivo imaging system (PerkinElmer, IVIS Spectrum) to measure total bioluminescent flux (photons/second) in each hindlimb.

### In vitro transfection of THP-1 monocytes

THP-1 monocytes were cultured in RPMI-1640 (ATCC modification) containing 10% heat-inactivated fetal bovine serum and 1% Penicillin-Streptomycin at 37°C, 5% CO_2_. Cells were plated at a density of 100 000 cells per well in a 96-well plate and transfected in quadruplicates with 500 ng/well of firefly luciferase mRNA-LNPs. Cell viability and firefly luciferase expression were determined after 24 hours with a plate reader assay, following the manufacturer’s protocol (Promega, E7110).

### Base hydrolysis

Ionizable lipids (2 μmol) were hydrolyzed with 2.5 eq. of KOH (200 μl, 0.025 M in deuterated methanol) in a 3 mm NMR tube (Norell, Z693189). ^1^H-NMR was acquired with a 700 MHz Agilent DD2 Spectrometer every 5 minutes for a total of 180 minutes. Additionally, spectra were acquired at 24, 48, 72, 144, 192, and 240 hours for βN2. Data acquisition started ∼10 minutes after the start of the reaction due to sample preparation and tune/lock/shim. Ester peaks were integrated using MestReNova 14.3.1 (Mestrelab Research, Santiago de Compostela, Spain) and normalized to value at t = 0 minute. The data series were fitted to a one phase decay exponential function (y = e^-kt^), where k is the rate of hydrolysis and ln(2)/k is the hydrolysis half-life.

### Pharmacokinetics of ionizable lipids

This study was conducted according to Transpharmation Canada Ltd. (Fergus, Ontario, Canada) standard operating procedures. Female CD-1 mice (n = 3-7) were injected intramuscularly in the right quadricep with 1 µg hEPO mRNA-LNPs, corresponding to 12.312 µg of βN2 or 12.764 µg of δO3. Animals were group housed in metabolic cages and euthanized at 3, 6, 12, 24, 48, 96, and 168 hours for the collection of blood, liver, spleen, and muscle. A single pooled urine and feces sample was also collected from each cage. Blood samples were divided for bioanalysis and diagnostic testing. Livers and feces were homogenized in DPBS using Qiagen TissueRuptor. Spleens and muscles were homogenized in DPBS using Precellys 24 and tubes containing ceramic beads. Tandem liquid chromatography-mass spectrometry (LC-MS/MS) methods were developed for the quantification of βN2 and δO3 in mouse plasma, muscle, liver, spleen, urine, and feces. The qualified methods were used to determine the concentration of the intact lipid in each matrix, using the other lipid as an internal standard. Samples were injected and separated on ZORBAX Eclipse XDB-C18 or ACE 3 C4-300 Å columns, with solvent A containing acetonitrile/H_2_O/1M Ammonium formate (600/390/10 with 0.1% formic acid) and solvent B containing isopropanol/acetonitrile/1M Ammonium formate (900/90/10 with 0.1%FA). A triple-quadrupole MS/MS system (Applied Biosystems, QTRAP 6500 or QTRAP 4000) operated in positive ion mode was used for signal detection. The pharmacokinetics parameters were estimated using Phoenix® WinNonlin 8.3 (Certara, Mountainview, California, USA).

### Human EPO expression in serum and cytokine analysis

BALB/c mice (n = 4 per timepoint) were injected intramuscularly in the left quadricep with 0.5 µg of hEPO mRNA-LNPs or 2×10^11^ empty LNPs. Mice were euthanized at 6, 24, and 48 hours post-injection. Blood was collected by cardiac puncture in serum tubes, incubated for 30 minutes at room temperature, and centrifuged at 10,000 x g for 5 minutes to isolate serum. Serum samples were diluted two-fold in 1X PBS and analyzed by Eve Technologies (Calgary, Alberta, Canada) using a 32-plex cytokine/chemokine array and human myokine custom assay to detect hEPO.

### mRNA vaccination

BALB/c mice (n *=* 8) were injected intramuscularly in the left quadricep with 0.5 μg of mRNA encoding the hemagglutinin protein from the tissue culture-adapted H1N1 Influenza A/Puerto Rico/8/1934 virus (ATCC, VR-1469) formulated in LNPs. Prime and boost doses were administered 30 days apart.

### HAI titer

4 weeks post-prime (pre-boost) and 2 weeks post-boost, blood was collected from the saphenous vein on the contralateral hindlimb using capillary blood collection tubes. Hemagglutinin inhibition (HAI) titers of serum samples were determined using a protocol adapted from the World Health Organization manual.^41^ First, sera were treated with receptor-destroying enzyme (Hardy Diagnostics, 370013) at 37°C for 18 hours to inactivate nonspecific inhibitors, followed by incubation at 56°C for 60 minutes to inactivate receptor-destroying enzyme per the manufacturer’s instructions. 25 µL of sera were serially diluted (1:2) with DPBS in a 96-well V bottom plate, starting with a 1:20 dilution and up to a 1:2560 dilution. 4 hemagglutination units of H1N1 Influenza A/Puerto Rico/8/1934 virus (ATCC, VR-1469) in 25 µL were added to diluted sera and incubated for 30 minutes at room temperature. Finally, 50 µL of a 0.5% v/v turkey erythrocyte solution (Rockland Immunochemicals, RLR408) was added to each well and incubated for 45 minutes at room temperature, after which the plate was turned on the side for one minute. Inhibition of agglutination was observed as the blood forming a “tear drop” shape. HAI titers were expressed as the inverse of the highest dilution with completely non-agglutinated erythrocytes. Each plate included a back-titration to confirm the virus dose (4 HAU/25 μl) and a negative control (serum from an untreated mouse).

### IFN-γ ELISpot

3 weeks post-boost, mice spleens were collected in 5 mL of RPMI 1640 supplemented with 1% penicillin-streptomycin and 10% heat inactivated fetal bovine serum. Spleens were dissociated in gentleMACS C Tubes (Miltenyi Biotec, 130-093-237), filtered through a 70 µm mesh and treated with ammonium-chloride-potassium lysis buffer to eliminate red blood cells. Splenocytes were seeded at 250 000 cells per well on precoated PVDF plates in serum-free CLT-Test medium (Cellular Technology Limited, CTLT-005) supplemented with 2mM fresh L-glutamine and 1% Penicillin-Streptomycin. Samples were tested individually by ex-vivo stimulation with overlapping 15-mer peptides (JPT Peptide Technologies) covering the entire hemagglutinin protein of the influenza A/PR/8/34 virus (1 µg/mL). Stimulation with concanavalin A (Thermo Fisher, 00-4978-03) was used as a positive control and test medium was used as a negative control. Cells were incubated for 18 hours at 37°C, 5% CO_2_. T cell ELISpot was performed using a Mouse IFN-γ single color kit according to the manufacturer’s instructions (Cellular Technology Limited, mIFNg-1M). Spot-forming units (SFU) were counted using an automated ELISpot reader (Cellular Technology Limited, S6 Entry M2).

### Statistics

Statistical analyses were performed using GraphPad Prism 9. *P < 0.05, **P < 0.01,***P < 0.001, and ****P < 0.0001 were considered statistically significant. ns = not significant.

## Acknowledgments

We thank Darcy C. Burns at University of Toronto CSICOMP NMR Facility and Matthew Forbes at the University of Toronto AIMS Mass Spectrometry Laboratory. J.C-S. thanks: Wildcat Foundation; B. and F. Milligan; Canadian Graduate Scholarship (CGS); and Emerging and Pandemic Infections Consortium (EPIC).

G.T thanks Ontario Graduate Scholarship (OGS) and the Precision Medicine (PRiME) Fellowship. O.F.K. thanks the following funding sources: University of Toronto’s Medicine by Design initiative and Pivotal Experiment Fund Round 3 which receive funding from the Canada First Research Excellence Fund (CFREF-2020); New Frontiers in Research – Exploration (NFRFE–2022– 00691); Canada Research Chairs Program (CRC-2020-00097); Natural Sciences and Engineering Research Council of Canada (RGPIN-2021-03421 and DGECR-2021-00467); Canadian Institutes of Health Research (PTT-184009); the Canada Foundation for Innovation John R. Evans Leaders Fund and Ontario Research Fund (41323); and the McCharles Prize for Early Career Research Distinction.

## Author contributions

J.C.-S., G.T. and O.F.K. conceptualized the project. J.C.-S. designed and performed experiments. G.T. synthesized ionizable lipids. J.C.-S. wrote the initial manuscript draft. All authors contributed to reviewing and editing the manuscript.

## Supplementary information

Supplementary information is available for this paper.

## References

1. Alameh, M.-G., Weissman, D. & Pardi, N. Messenger RNA-Based Vaccines Against Infectious Diseases. in mRNA Vaccines (eds. Yu, D. & Petsch, B.) 111–145 (Springer International Publishing, Cham, 2022). doi:10.1007/82_2020_202.

2. Xie, W., Chen, B. & Wong, J. Evolution of the market for mRNA technology. Nat. Rev. Drug Discov. 20, 735–736 (2021).

3. Lau, Y. M. A., Pang, J., Tilstra, G., Couture-Senécal, J. & Khan, O. F. The engineering challenges and opportunities when designing potent ionizable materials for the delivery of ribonucleic acids. Expert Opin. Drug Deliv. 19, 1650–1663 (2022).

4. Pfizer. COMIRNATY [package insert]. U.S. Food and Drug Administration website.https://www.fda.gov/media/151707/download?attachment. Revised October 2023.

5. Moderna. SPIKEVAX [package insert]. U.S. Food and Drug Administration website.https://www.fda.gov/media/155675/download. Revised April 2024.

6. Moderna. MRESVIA [package insert]. U.S. Food and Drug Administration website.https://www.fda.gov/media/179005/download. Revised May 2024.

7. Hassett, K. J. et al. Optimization of Lipid Nanoparticles for Intramuscular Administration of mRNA Vaccines. Mol. Ther. Nucleic Acids 15, 1–11 (2019).

8. Tilstra, G. et al. Iterative Design of Ionizable Lipids for Intramuscular mRNA Delivery. J. Am. Chem. Soc. 145, 2294–2304 (2023).

9. Lam, K. et al. Unsaturated, Trialkyl Ionizable Lipids are Versatile Lipid-Nanoparticle Components for Therapeutic and Vaccine Applications. Adv. Mater. 35, 2209624 (2023).

10. Goldman, R. L. et al. Understanding structure activity relationships of good HEPES lipids for lipid nanoparticle mRNA vaccine applications. Biomaterials 122243 (2023) doi:10.1016/j.biomaterials.2023.122243.

11. Parhiz, H., Atochina-Vasserman, E. N. & Weissman, D. mRNA-based therapeutics: looking beyond COVID-19 vaccines. The Lancet 403, 1192–1204 (2024).

12. Rohner, E., Yang, R., Foo, K. S., Goedel, A. & Chien, K. R. Unlocking the promise of mRNA therapeutics. Nat. Biotechnol. 40, 1586–1600 (2022).

13. Jackson, N. A. C., Kester, K. E., Casimiro, D., Gurunathan, S. & DeRosa, F. The promise of mRNA vaccines: a biotech and industrial perspective. Npj Vaccines 5, 1–6 (2020).

14. Verbeke, R., Hogan, M. J., Loré, K. & Pardi, N. Innate immune mechanisms of mRNA vaccines. Immunity 55, 1993–2005 (2022).

15. Karikó, K., Buckstein, M., Ni, H. & Weissman, D. Suppression of RNA Recognition by Toll-like Receptors: The Impact of Nucleoside Modification Couture-Senécal, Tilstra, and Khan. bioRxiv. 2024. 12 and the Evolutionary Origin of RNA. Immunity 23, 165–175 (2005).

16. Nelson, J. et al. Impact of mRNA chemistry and manufacturing process on innate immune activation. Sci. Adv. 6, eaaz6893 (2020).

17. Alameh, M.-G. et al. Lipid nanoparticles enhance the efficacy of mRNA and protein subunit vaccines by inducing robust T follicular helper cell and humoral responses. Immunity 54, 2877-2892.e7 (2021).

18. Tahtinen, S. et al. IL-1 and IL-1ra are key regulators of the inflammatory response to RNA vaccines. Nat. Immunol. 23, 532–542 (2022).

19. Ndeupen, S. et al. The mRNA-LNP platform’s lipid nanoparticle component used in preclinical vaccine studies is highly inflammatory. iScience 24, 103479 (2021).

20. Omo-Lamai, S. et al. Lipid Nanoparticle-Associated Inflammation is Triggered by Sensing of Endosomal Damage: Engineering Endosomal Escape Without Side Effects. 2024.04.16.589801 Preprint at 10.1101/2024.04.16.589801 (2024).

21. Polack, F. P. et al. Safety and Efficacy of the BNT162b2 mRNA Covid-19 Vaccine. N. Engl. J. Med. 383, 2603–2615 (2020).

22. Jackson, L. A. et al. An mRNA Vaccine against SARS-CoV-2 — Preliminary Report. N. Engl. J. Med. 383, 1920–1931 (2020).

23. Wilson, E. et al. Efficacy and Safety of an mRNA-Based RSV PreF Vaccine in Older Adults. N. Engl. J. Med. 389, 2233–2244 (2023).

24. Lee, I. T. et al. Safety and immunogenicity of a phase 1/2 randomized clinical trial of a quadrivalent, mRNA-based seasonal influenza vaccine (mRNA-1010) in healthy adults: interim analysis. Nat. Commun. 14, 3631 (2023).

25. Maier, M. A. et al. Biodegradable Lipids Enabling Rapidly Eliminated Lipid Nanoparticles for Systemic Delivery of RNAi Therapeutics. Mol. Ther. 21, 1570–1578 (2013).

26. Sabnis, S. et al. A Novel Amino Lipid Series for mRNA Delivery: Improved Endosomal Escape and Sustained Pharmacology and Safety in Non-human Primates. Mol. Ther. 26, 1509–1519 (2018).

27. Burdette, D. et al. Systemic Exposure, Metabolism, and Elimination of [14C]-Labeled Amino Lipid, Lipid 5,After a Single Administration of mRNA Encapsulating Lipid Nanoparticles to Sprague Dawley Rats. Drug Metab. Dispos. DMD-AR-2022-001194 (2023) doi:10.1124/dmd.122.001194.

28. Akinc, A. et al. A combinatorial library of lipid-like materials for delivery of RNAi therapeutics. Nat. Biotechnol. 26, 561–569 (2008).

29. Suzuki, Y. et al. Biodegradable lipid nanoparticles induce a prolonged RNA interference-mediated protein knockdown and show rapid hepatic clearance in mice and nonhuman primates. Int. J. Pharm. 519, 34–43 (2017).

30. Qiu, M. et al. Lung-selective mRNA delivery of synthetic lipid nanoparticles for the treatment of pulmonary lymphangioleiomyomatosis. Proc. Natl. Acad. Sci. 119, e2116271119 (2022).

31. Carrasco, M. J. et al. Ionization and structural properties of mRNA lipid nanoparticles influence expression in intramuscular and intravascular administration. Commun. Biol. 4, 1–15 (2021).

32. Jayaraman, M. et al. Maximizing the Potency of siRNA Lipid Nanoparticles for Hepatic Gene Silencing In Vivo**. Angew. Chem. 124, 8657–8661 (2012).

33. Khan, O. F. Best practices for characterizing RNA lipid nanoparticles. Nat. Rev. Methods Primer 3, 65 (2023).

34. Alnylam Pharmaceuticals. ONPATTRO [prescribing information]. U.S. Food and Drug Administration website.https://www.accessdata.fda.gov/drugsatfda_docs/label/2018/210922s000lbl.pdf. Revised August 2018.

35. Arunachalam, P. S. et al. Systems vaccinology of the BNT162b2 mRNA vaccine in humans. Nature 596, 410–416 (2021).

36. Li, C. et al. Mechanisms of innate and adaptive immunity to the Pfizer-BioNTech BNT162b2 vaccine. Nat. Immunol. 23, 543–555 (2022).

37. Bergamaschi, C. et al. Systemic IL-15, IFN-γ, and IP-10/CXCL10 signature associated with effective immune response to SARS-CoV-2 in BNT162b2 mRNA vaccine recipients. Cell Rep. 36, (2021).

38. Petsch, B. et al. Protective efficacy of in vitro synthesized, specific mRNA vaccines against influenza A virus infection. Nat. Biotechnol. 30, 1210–1216 (2012).

39. Pardi, N. et al. Nucleoside-modified mRNA vaccines induce potent T follicular helper and germinal center B cell responses. J. Exp. Med. 215, 1571–1588 (2018).

40. Arevalo, C. P. et al. A multivalent nucleoside-modified mRNA vaccine against all known influenza virus subtypes. Science 378, 899–904 (2022).

41. World Health Organization. Manual for the laboratory diagnosis and virological surveillance of influenza. WHO Glob. Influenza Surveill. Netw. Man. Lab. Diagn. Virol. Surveill. Influenza (2011).

42. Hassett, K. J. et al. mRNA Vaccine Trafficking and Resulting Protein Expression After Intramuscular Administration. Mol. Ther. - Nucleic Acids 0, (2023).

43. Mosmann, T. R., McMichael, A. J., LeVert, A., McCauley, J. W. & Almond, J. W. Opportunities and challenges for T cell-based influenza vaccines. Nat. Rev. Immunol. 1–17 (2024) doi:10.1038/s41577-024-01030-8.

44. Baiersdörfer, M. et al. A Facile Method for the Removal of dsRNA Contaminant from In Vitro-Transcribed mRNA. Mol. Ther. - Nucleic Acids 15, 26–35 (2019).

45. Manning, A. M. et al. Ionizable Lipid with Supramolecular Chemistry Features for RNA Delivery In Vivo. Small n/a, 2302917. Couture-Senécal, Tilstra, and Khan. bioRxiv. 2024. 13

