## Supplementary information for "Engineering ionizable lipids for rapid biodegradation balances mRNA vaccine efficacy and tolerability"

### Supplementary Methods

*Materials*

Reagents and solvents were purchased from Sigma Aldrich, TCI, Thermo Scientific Chemicals, Fisher Scientific, Alfa Aesar and used as received, unless otherwise noted. Dry solvents were prepared by adding molecular sieves (Sigma Aldrich, 3 Å, 8-12 mesh) directly to solvent bottles for at least 48 hours prior to use.

*General Synthesis Procedures*

All reactions were carried out in standard laboratory glassware under ambient atmospheric conditions. All intermediates and final compounds were covered with foil and stored away from light.

Reactions were monitored by thin-layer chromatography (Supelco TLC Silica Gel 60 $F_{254}$) and visualized using an iodine stain. TLC eluent was either 10% MeOH in dichloromethane, or 35% (22% MeOH, 3% $NH_4OH$ in dichloromethane) in dichloromethane.

Flash chromatography was carried out using an automated system (Buchi Pure C-815 Flash Chromatography System, ELSD detector) on pre-packed normal-phase silica cartridges (Buchi EcoFlex, 50uM irregular particles). Compounds were prepared for flash chromatography by adsorption to silica gel (Sigma Aldrich, technical grade, pore size 60 Å, 230-400 mesh, 40-63um particle size).

*Structural characterization*

Proton nuclear magnetic resonance ($^1H$ NMR) spectra were recorded on either an Agilent DD2-500 or Bruker Avance III-400 spectrometer, are reported in parts per million (ppm) downfield of tetramethylsilane, and are referenced to the residual proton signal of the NMR solvent ($CDCl_3$: 7.26ppm, [$CHCl_3$]). Proton-decoupled carbon-13 nuclear magnetic resonance ($^{13}C\{^1H\}$ NMR) spectra were recorded on either an Agilent DD2-500 (with cryogenic-cooled probe) or Bruker Avance III-400 spectrometer, are reported in parts per million (ppm) downfield of tetramethylsilane, and are referenced to the signal of the NMR solvent ($CDCl_3$: 77.16ppm). NMR peaks are reported as: br = broad, s = singlet, d = doublet, t = triplet, q = quartet, p = pentet, and m, multiplet; coupling constants ($J$) are reported in Hz.

Electrospray ionization mass spectrometry (ESI-MS) was performed by Matthew Forbes at the Advanced Instrumentation for Molecular Structure facility at the University of Toronto using an Agilent 6538 UHD spectrometer.

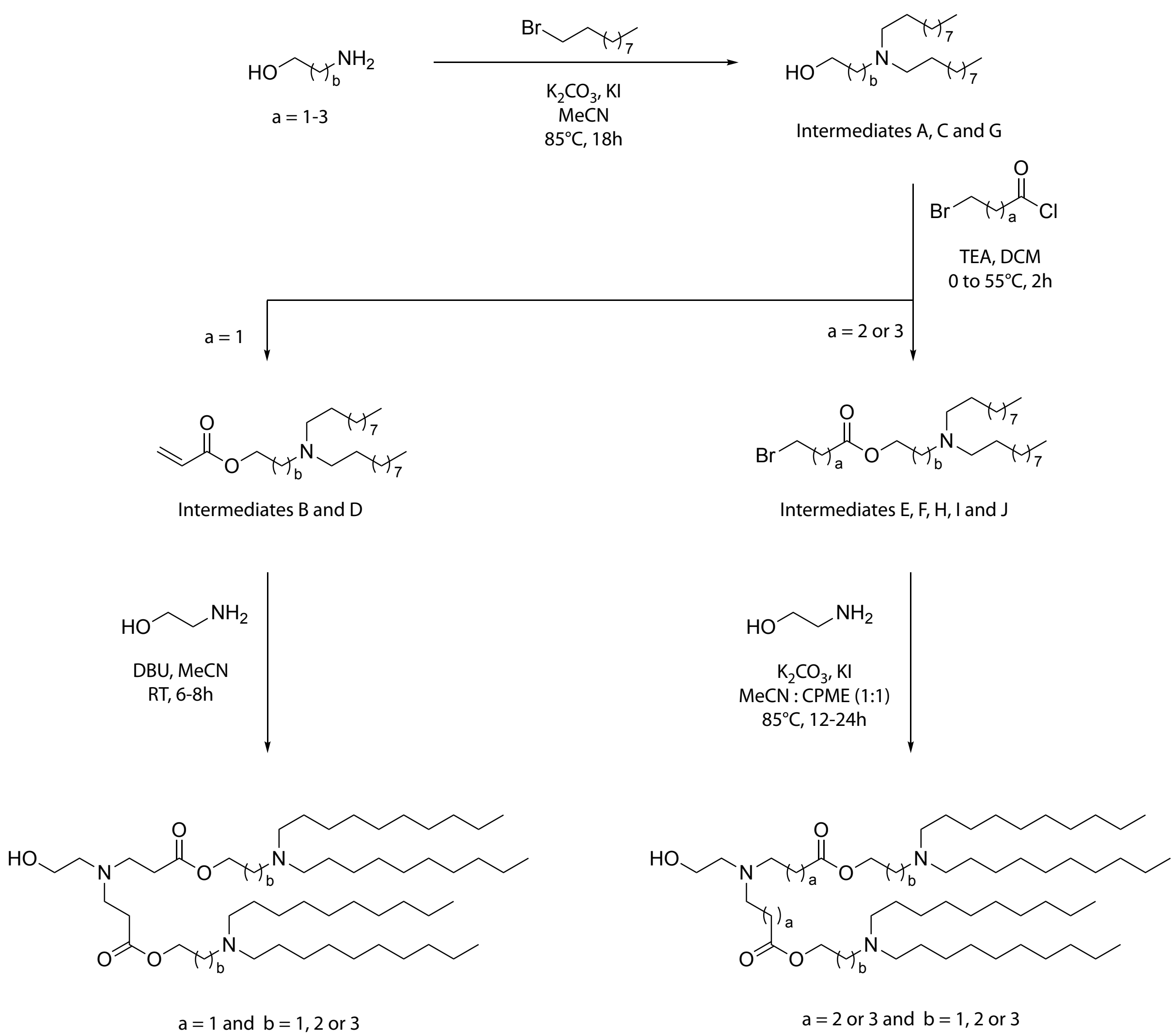


**Scheme 1. General synthesis of ionizable lipids containing ester linkers.**

*General Synthesis Method A – Alkylation of amino-alcohols*

Amino alcohol (1 equivalent) and 1-bromodecane (2.2 equivalents) were dissolved in acetonitrile (dried over molecular sieves) in a round bottom flask. $K_2CO_3$ (4.4 equivalents) and KI (0.2 equivalents) were added. The flask was then covered with aluminum foil, fitted with a condenser, and heated to 85°C for 18 hours. The reaction mixture was filtered through a celite plug and washed thoroughly with ethyl acetate. The filtrate was concentrated to a crude oil and then purified by column chromatography using a 10-40% (22% MeOH, 3% $NH_4OH$ in dichloromethane) in dichloromethane (DCM) gradient.

*General Synthesis Method B – Esterification of acid chlorides*

Alcohol (1.2 equivalents) and triethylamine (2 equivalents) were dissolved in dichloromethane (dried over molecular sieves) in a round bottoms flask and cooled to 0°C. Acid chloride (1 equivalent) was added slowly to the reaction while stirring. The ice bath was removed, and the flask was fitted with a condenser and heated to 55°C for 2 hours. The reaction mixture was concentrated to a crude oil and then purified by column chromatography using a 0-60% ethyl acetate in hexanes gradient.

*General Synthesis Method C – Alkylation of ethanolamine core*

Ethanolamine (1 equivalent) and halide substrate (1.8 equivalents) were dissolved in a 1:1 mixture of acetonitrile (dried over molecular sieves) : cyclopentyl methyl ether in a glass vial. $K_2CO_3$ (4 equivalents) and KI (0.4 equivalents) were added. The vial was then covered with aluminum foil and heated to 85°C for 12-24 hours. The reaction mixture was filtered through a celite plug and washed thoroughly with ethyl acetate. The filtrate was concentrated to a crude oil and then purified by column chromatography using a 10-40% (22% MeOH, 3% $NH_4OH$ in dichloromethane) in dichloromethane gradient.

*Synthesis of Intermediate A*

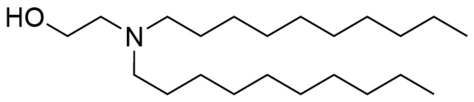

Intermediate A was synthesized according to General Synthesis Method A using ethanolamine and isolated as a pale yellow oil (80% yield).$^1$H NMR (500 MHz, $CDCl_3$) δ 3.56 (t, $J$ = 5.6 Hz, 0H), 3.18 (s, 0H), 2.61 (t, $J$ = 5.4 Hz, 0H), 2.52 – 2.43 (m, 1H), 1.45 (p, $J$ = 7.2 Hz, 1H), 1.32 – 1.21 (m, 5H), 0.88 (t, $J$ = 7.2 Hz, 1H). $^{13}$C NMR (126 MHz, $CDCl_3$) δ 58.29, 55.70, 54.00, 32.04, 29.77, 29.73, 29.69, 29.68, 29.46, 27.53, 27.03, 22.82, 14.25.

*Synthesis of Intermediate B*

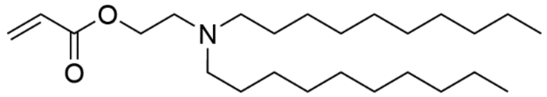

Intermediate B was synthesized according to General Synthesis Method B using Intermediate A and 3-bromopropionyl chloride and isolated as a light yellow oil that slowly transitioned to a crystalline solid (80% yield). $^1$H NMR (500 MHz, $CDCl_3$) δ 6.39 (dd, $J$ = 17.3, 1.5 Hz, 1H), 6.12 (dd, $J$ = 17.4, 10.5 Hz, 1H), 5.81 (dd, $J$ = 10.4, 1.5 Hz, 1H), 4.20 (t, $J$ = 6.2 Hz, 2H), 2.72 (t, $J$ = 6.3 Hz, 2H), 2.48 – 2.42 (m, 4H), 1.42 (p, $J$ = 6.6 Hz, 4H), 1.26 (d, $J$ = 5.2 Hz, 29H), 0.87 (t, $J$ = 7.0 Hz, 6H). $^{13}$C NMR (126 MHz, $CDCl_3$) δ 166.36, 130.69, 128.71, 63.09, 55.00, 52.30, 32.05, 29.82, 29.81, 29.76, 29.75, 29.48, 27.60, 27.42, 22.83, 14.25.

*Synthesis of Intermediate C*

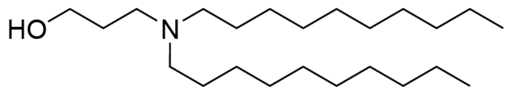

Intermediate C was synthesized according to General Synthesis Method A using 3-amino-1-propanol and isolated as a light yellow oil (80% yield). $^1$H NMR (500 MHz, $CDCl_3$) δ 3.78 (t, $J$ = 5.3 Hz, 2H), 2.69 (t, $J$ = 5.8 Hz, 2H), 2.50 – 2.42 (m, 4H), 1.74 – 1.66 (m, 2H), 1.53 – 1.44 (m, 4H), 1.33 – 1.17 (m, 30H), 0.86 (t, $J$ = 7.4 Hz, 6H).$^{13}$C NMR (126 MHz, $CDCl_3$) δ 64.29, 55.04, 54.11, 32.01, 29.71, 29.69, 29.65, 29.42, 27.68, 27.55, 26.47, 22.79, 14.22.

*Synthesis of Intermediate D*

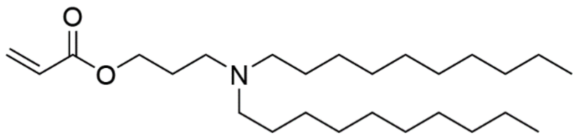

Intermediate D was synthesized according to General Synthesis Method B using Intermediate C and 3-bromopropionyl chloride and isolated as a light yellow oil that slowly transitioned to a crystalline solid (85% yield). $^1$H NMR (500 MHz, $CDCl_3$) δ 6.38 (dd, $J$ = 17.3, 1.5 Hz, 1H), 6.10 (dd, $J$ = 17.3, 10.4 Hz, 1H), 5.79 (dd, $J$ = 10.4, 1.5 Hz, 1H), 4.19 (t, $J$ = 6.5 Hz, 2H), 2.49 (t, $J$ = 7.2 Hz, 2H), 2.41 – 2.34 (m, 4H), 1.79 (p, $J$ = 7.1 Hz, 2H), 1.45 – 1.35 (m, 4H), 1.34 – 1.23 (m, 27H), 0.92 – 0.80 (m, 6H). $^{13}$C NMR (126 MHz, $CDCl_3$) δ 166.34, 130.52, 128.74, 63.20, 54.29, 50.51, 32.04, 29.79, 29.75, 29.74, 29.47, 27.68, 27.19, 26.52, 22.81, 14.23.

*Synthesis of Intermediate E*

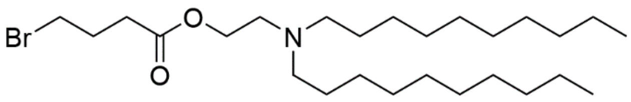


Intermediate E was synthesized according to General Synthesis Method B using Intermediate A and 4-bromobutyryl chloride and isolated as a light yellow oil (54% yield). $^{1}$H NMR (500 MHz, $CDCl_3$) δ 4.21 – 4.11 (m, 2H), 3.46 (t, *J* = 6.5 Hz, 1H), 2.70 (dt, *J* = 12.1, 6.1 Hz, 2H), 2.55 – 2.39 (m, 5H), 2.21 – 2.12 (m, 1H), 1.51 – 1.37 (m, 4H), 1.34 – 1.19 (m, 29H), 0.88 (t, *J* = 6.9 Hz, 6H). $^{13}$C NMR (126 MHz, $CDCl_3$) δ 174.89, 172.50, 62.81, 54.78, 54.76, 54.64, 52.20, 52.14, 32.68, 32.47, 31.90, 29.66, 29.65, 29.61, 29.60, 29.59, 29.59, 29.33, 27.76, 27.44, 27.44, 27.12, 22.67, 14.10, 12.88, 8.39.

*Synthesis of Intermediate F*

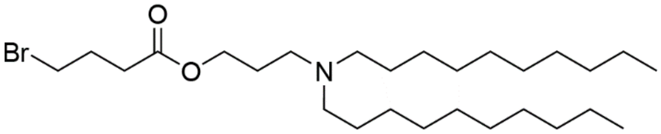


Intermediate F was synthesized according to General Synthesis Method B using Intermediate C and 4-bromobutyryl chloride and isolated as a light yellow oil (101% yield with remaining solvent). $^{1}$H NMR (500 MHz, $CDCl_3$) δ 4.13 (t, *J* = 6.5 Hz, 2H), 3.46 (t, *J* = 6.4 Hz, 2H), 2.50 (t, *J* = 7.1 Hz, 4H), 2.46 – 2.39 (m, 4H), 2.22 – 2.09 (m, 2H), 1.84 – 1.75 (m, 2H), 1.49 – 1.39 (m, 4H), 1.35 – 1.19 (m, 29H), 0.87 (t, *J* = 7.0 Hz, 6H). $^{13}$C NMR (126 MHz, $CDCl_3$) δ 172.66, 63.21, 54.19, 50.55, 32.86, 32.59, 32.05, 29.80, 29.78, 29.74, 29.73, 29.48, 27.89, 27.65, 22.83, 14.26.

*Synthesis of Intermediate G*

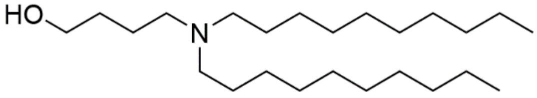


Intermediate G was synthesized according to General Synthesis Method A using 4-amino-1-butanol and isolated as a light yellow oil (58% yield). $^{1}$H NMR (500 MHz, $CDCl_3$) δ 3.57 (t, *J* = 5.4 Hz, 2H), 2.60 – 2.52 (m, 6H), 1.75 – 1.61 (m, 4H), 1.58 – 1.48 (m, 4H), 1.33 – 1.17 (m, 28H), 0.87 (t, *J* = 6.9 Hz, 5H). $^{13}$C NMR (126 MHz, $CDCl_3$) δ 77.27, 77.02, 76.76, 62.32, 54.25, 53.36, 31.99, 31.86, 29.55, 29.52, 29.40, 29.27, 27.45, 25.18, 22.65, 14.08.

*Synthesis of Intermediate H*

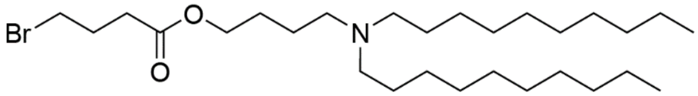


Intermediate H was synthesized according to General Synthesis Method B using Intermediate G and 4-bromobutyryl chloride and isolated as a light yellow oil (95% yield). $^{1}$H NMR (500 MHz, $CDCl_3$) δ 4.09 (t, *J* = 6.7 Hz, 2H), 3.46 (t, *J* = 6.5 Hz, 2H), 2.49 (t, *J* = 7.2 Hz, 2H), 2.43 – 2.36 (m, 3H), 2.40 – 2.33 (m, 5H), 2.17 (p, *J* = 6.4 Hz, 2H), 1.63 (p, *J* = 7.9 Hz, 2H), 1.52 – 1.43 (m, 2H), 1.44 – 1.35 (m, 4H), 1.36 – 1.18 (m, 29H), 0.87 (t, *J* = 7.1 Hz, 6H). $^{13}$C NMR (126 MHz, $CDCl_3$) δ 172.73, 64.84, 60.52, 54.34, 53.85, 32.84, 32.63, 32.06, 31.73, 29.83, 29.80, 29.76, 29.49, 27.92, 27.79, 27.22, 26.83, 23.81, 22.83, 14.34, 14.26.

*Synthesis of Intermediate I*

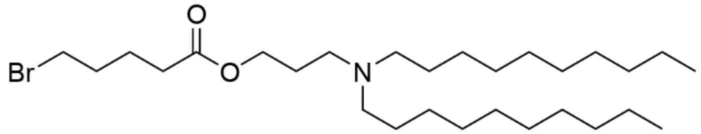


Intermediate I was synthesized according to General Synthesis Method B using Intermediate C and 5-bromovaleyryl chloride and isolated as a light yellow oil (97% yield). $^{1}$H NMR (500 MHz, $CDCl_3$) δ 4.10 (t, *J* = 6.5 Hz, 2H), 3.40 (t, *J* = 6.6 Hz, 2H), 2.48 – 2.42 (m, 2H), 2.35 (t, *J* = 7.4 Hz, 4H), 2.33 (t, *J* = 7.4 Hz, 2H), 1.93 – 1.85 (m, 2H), 1.82 – 1.69 (m, 4H), 1.39 (p, *J* = 7.1 Hz, 5H), 1.33 – 1.19 (m, 27H), 0.87 (t, *J* = 7.0 Hz, 6H). $^{13}$C NMR (126 MHz, $CDCl_3$) δ 173.25, 63.22, 54.33, 50.59, 33.45, 33.09, 32.15, 32.05, 31.72, 29.82, 29.79, 29.76, 29.48, 27.71, 27.30, 26.65, 23.67, 22.82, 22.78, 14.25.

*Synthesis of Intermediate J*

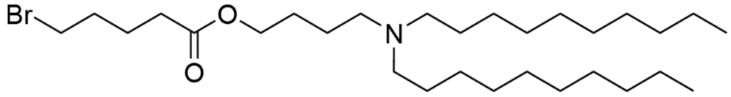


Intermediate J was synthesized according to General Synthesis Method B using Intermediate G and 5-bromovaleyryl chloride and isolated as a light yellow oil (95% yield). $^{1}$H NMR (500 MHz, $CDCl_3$) δ 4.08 (t, *J* = 6.7 Hz, 2H), 3.40 (t, *J* = 6.6 Hz, 2H), 2.40 (t, *J* = 7.5 Hz, 2H), 2.38 – 2.35 (m, 4H), 2.33 (t, *J* = 6.9 Hz, 2H), 1.93 – 1.86 (m, 2H), 1.82 – 1.73 (m, 2H), 1.63 (p, *J* = 7.9 Hz, 2H), 1.51 – 1.44 (m, 2H), 1.40 (p, *J* = 6.9 Hz, 4H), 1.32 – 1.19 (m, 29H), 0.87 (t, *J* = 7.1 Hz, 6H). $^{13}$C NMR (126 MHz, $CDCl_3$) δ 173.17, 64.53, 54.18, 53.71, 33.30, 32.96, 32.00, 31.90, 29.67, 29.64, 29.60, 29.58, 29.33, 27.63, 27.07, 26.70, 23.67, 23.52, 22.67, 14.10.

*Synthesis of Intermediate K*

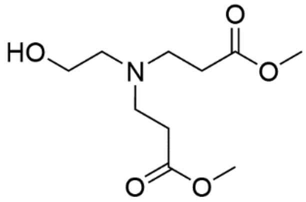


Ethanolamine (90.7µL, 1.5mmol, 1eq) and methyl acrylate (299µL, 3.3mmol, 2.2eq) were dissolved in methanol (5mL), covered with foil, and stirred at room temperature overnight. Excess solvent and methyl acrylate were removed under reduced pressure to produce a clear, colourless oil (349.9mg, 1.5mmol, quantitative yield). $^{1}$H NMR (500 MHz, $CDCl_3$) δ 3.66 (s, 6H), 3.56 (t, *J* = 5.1 Hz, 2H), 2.98 – 2.86 (m, 1H), 2.77 (t, *J* = 6.7 Hz, 4H), 2.56 (t, *J* = 5.2 Hz, 2H), 2.44 (t, *J* = 6.7 Hz, 4H). $^{13}$C NMR (126 MHz, $CDCl_3$) δ 173.14, 59.19, 56.05, 51.77, 49.27, 32.69.

*Synthesis of Intermediate L*

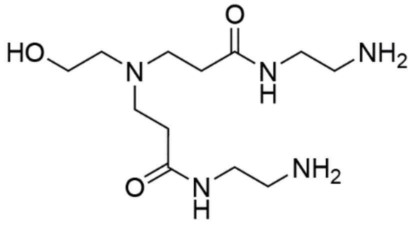


Ethylenediamine (1.001mL, 15mmol, 10eq) and Intermediate K (349.9mg, 1.5mmol, 1eq) were dissolved in methanol (10mL), covered with foil, and stirred for 3 days at room temperature. Excess ethylenediamine and solvent were removed under reduced pressure to produce a crude oil. The crude oil was washed 5x with 20mL of ice-cold diethyl ether to remove remaining ethylenediamine, to produce a light-yellow oil (185.3mg, 0.64mmol, 43%). $^{1}$H NMR (500 MHz, $CDCl_3$) δ 7.52 – 7.44 (m, 2H), 3.61 – 3.58 (m, 2H), 3.27 (q, *J* = 5.7 Hz, 4H), 2.82 (t, *J* = 5.8 Hz, 4H), 2.73 (t, *J* = 5.8 Hz, 4H), 2.54 (t, *J* = 4.9 Hz, 2H), 2.37 (t, *J* = 6.2 Hz, 4H), 2.22 (s, 6H). $^{13}$C NMR (126 MHz, $CDCl_3$) δ 173.00, 59.00, 56.79, 50.68, 50.58, 41.74, 41.40, 34.26.

*Synthesis of Compound βO2*

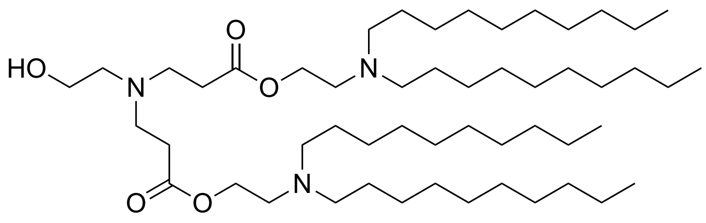

Ethanolamine (9.17uL, 0.15mmol, 0.4eq), Intermediate B (0.150g, 0.38mmol, 1eq), and 1,8-diazabicyclo(5.4.0)undec-7-ene (28.29uL, 0.19mmol, 0.5eq) were dissolved in acetonitrile (0.5mL) in a screw cap vial and stirred at ambient temperature for 6 hours. The reaction was concentrated and purified by column chromatography using a 10-40% (22% MeOH, 3% $NH_4OH$ in dichloromethane) in dichloromethane gradient. Product was isolated as a clear oil (13mg, 10%). $^1H$ NMR (400 MHz, $CDCl_3$) δ 4.13 (t, $J$ = 6.2 Hz, 4H), 3.60 – 3.52 (m, 2H), 2.78 (t, $J$ = 6.8 Hz, 5H), 2.73 – 2.64 (m, 6H), 2.57 (t, $J$ = 5.4 Hz, 2H), 2.51 – 2.40 (m, 13H), 1.41 (p, $J$ = 7.3 Hz, 9H), 1.34 – 1.17 (m, 70H), 0.86 (t, $J$ = 7.0 Hz, 14H). $^{13}C$ NMR (101 MHz, $CDCl_3$) δ 172.63, 59.27, 56.02, 54.79, 52.28, 49.21, 32.73, 32.02, 31.98, 29.79, 29.72, 29.64, 29.63, 29.45, 29.39, 27.57, 27.27, 27.08, 22.79, 22.77, 14.21, 14.20. MS (ESI+): m/z $[M+H]^+$ 852.81 for C52H105N3O5. Purity was >85% by mol as determined by $^1H$ NMR.

*Synthesis of Compound βO3*

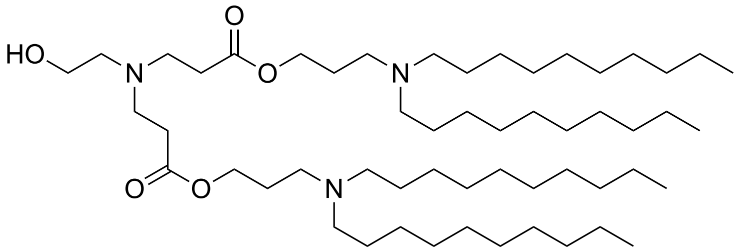

Ethanolamine (12.1uL, 0.20mmol, 1 eq) and Intermediate C (0.147g, 0.36mmol, 1.8eq) were combined in a screw cap vial and stirred at 70°C for 8 hours. Reaction was carried out without solvent. The reaction was concentrated and purified by column chromatography using a 10-40% (22% MeOH, 3% $NH_4OH$ in dichloromethane) in dichloromethane gradient. The product was isolated as a clear oil (61mg, 38% yield). $^1H$ NMR (400 MHz, $CDCl_3$) δ 4.10 (t, $J$ = 6.5 Hz, 4H), 3.56 (t, $J$ = 5.0 Hz, 2H), 2.78 (t, $J$ = 6.8 Hz, 4H), 2.51 (t, $J$ = 5.2 Hz, 17H), 1.89 – 1.74 (m, 4H), 1.51 – 1.38 (m, 8H), 1.33 – 1.18 (m, 57H), 0.87 (t, $J$ = 6.9 Hz, 12H). $^{13}C$ NMR (101 MHz, $CDCl_3$) δ 172.68, 68.68, 63.13, 59.17, 56.00, 53.99, 50.61, 49.21, 36.33, 32.74, 32.02, 29.77, 29.71, 29.68, 29.45, 27.59, 22.80, 14.23. MS (ESI+): m/z $[M+H]^+$ 880.84 for C54H109N3O5. Purity was >90% by mol as determined by $^1H$ NMR.

*Synthesis of Compound γO2*

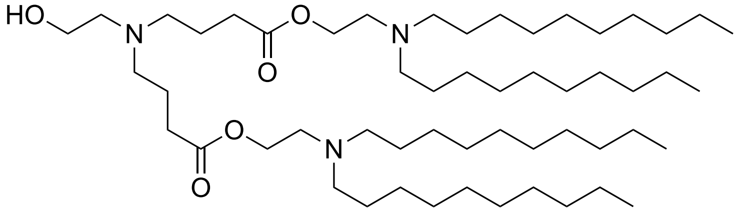

Compound *γO2* was synthesized according to General Synthesis Method C using Intermediate E. Product was isolated as a clear oil (24.9mg, 9% yield). $^1H$ NMR (500 MHz, $CDCl_3$) δ 4.11 (t, $J$ = 6.4 Hz, 4H), 3.53 (t, 2H), 2.67 (t, $J$ = 6.4 Hz, 4H), 2.58 (t, $J$ = 5.4 Hz, 2H), 2.48 (t, $J$ = 7.9 Hz, 4H), 2.43 (t, $J$ = 8.1 Hz, 8H), 2.31 (t, $J$ = 7.3 Hz, 3H), 1.76 (p, $J$ = 7.3 Hz, 4H), 1.46 – 1.35 (m, 8H), 1.32 – 1.20 (m, 56H), 0.87 (t, $J$ = 6.9 Hz, 12H). $^{13}C$ NMR (126 MHz, $CDCl_3$) δ 173.45, 62.72, 58.74, 55.71, 54.78, 52.80, 52.21, 31.91, 31.90, 29.67, 29.66, 29.64, 29.61, 29.60, 29.58, 29.33, 27.47, 27.16, 22.67, 22.21, 14.10. MS (ESI+): m/z $[M+H]^+$ 880.84 for C54H109N3O5. Purity was >80% by mol as determined by $^1H$ NMR.

*Synthesis of Compound γO3*

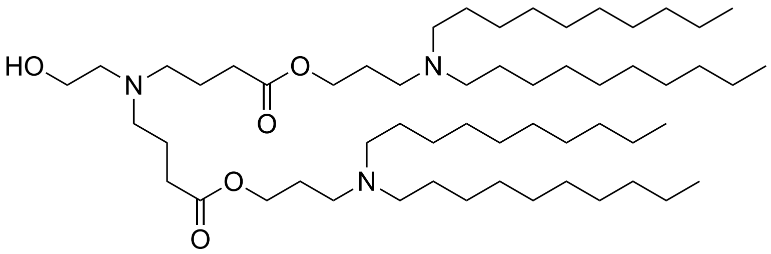


Compound *γO3* was synthesized according to General Synthesis Method C using Intermediate F. Product was isolated as a clear oil (22mg, 13%) $^{1}$H NMR (500 MHz, $CDCl_3$) δ 4.10 (t, *J* = 6.5 Hz, 4H), 3.79 (t, *J* = 5.4 Hz, 1H), 3.53 (t, *J* = 5.3 Hz, 2H), 2.70 (t, *J* = 5.7 Hz, 1H), 2.62 – 2.41 (m, 19H), 2.30 (t, *J* = 7.2 Hz, 4H), 1.87 – 1.67 (m, 9H), 1.45 (m, 9H), 1.34 – 1.19 (m, 72H), 0.87 (t, *J* = 6.9 Hz, 15H). $^{13}$C NMR (126 MHz, $CDCl_3$) δ 173.56, 62.97, 58.86, 55.88, 53.99, 52.97, 50.57, 32.03, 32.01, 29.77, 29.71, 29.69, 29.68, 29.64, 29.45, 29.43, 27.59, 27.54, 22.80, 22.38, 14.24. MS (ESI+): m/z $[M+H]^+$ 908.87 for C56H113N3O5. Purity was ~65% by mol by mol as determined by $^{1}$H NMR.

*Synthesis of Compound γO4*

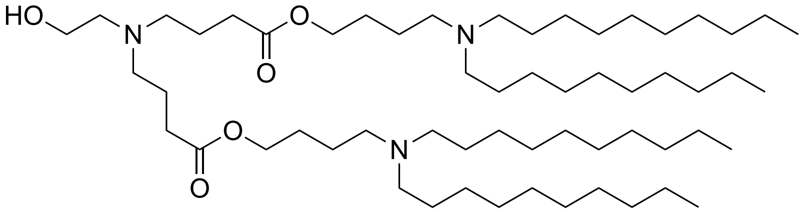


Compound *γO4* was synthesized according to General Synthesis Method C using Intermediate H. Product was isolated as a clear oil (78mg, 18% yield). $^{1}$H NMR (500 MHz, $CDCl_3$) δ 4.08 (t, *J* = 6.6 Hz, 4H), 3.53 (t, *J* = 5.4 Hz, 2H), 2.58 (t, *J* = 5.4 Hz, 2H), 2.49 (t, *J* = 7.1 Hz, 5H), 2.46 – 2.36 (m, 10H), 2.31 (t, *J* = 7.2 Hz, 4H), 1.76 (p, *J* = 7.3 Hz, 4H), 1.63 (p, *J* = 6.8 Hz, 4H), 1.51 (p, *J* = 8.2 Hz, 5H), 1.43 (p, *J* = 6.9 Hz, 8H), 1.26 (s, 56H), 0.87 (t, *J* = 6.8 Hz, 12H). $^{13}$C NMR (126 MHz, $CDCl_3$) δ 173.71, 64.55, 58.92, 55.90, 54.16, 53.77, 53.01, 32.10, 32.05, 29.81, 29.77, 29.75, 29.74, 29.48, 27.74, 26.82, 22.83, 22.43, 14.26. MS (ESI+): m/z $[M+H]^+$ 936.91 for C58H117N3O5. Purity was >95% by mol as determined by $^{1}$H NMR.

*Synthesis of Compound δO3*

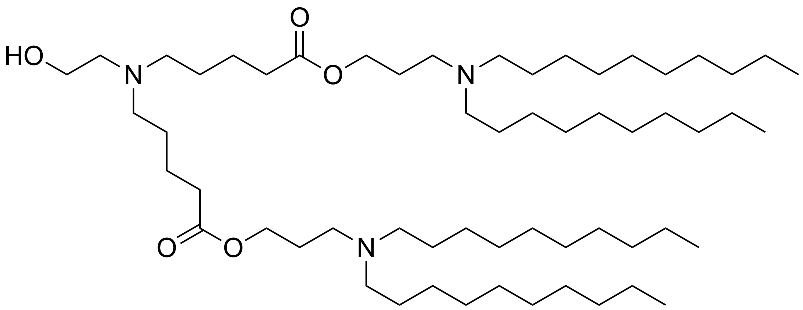


Compound δO3 was synthesized according to General Synthesis Method C using Intermediate I. Product was isolated as a clear oil (36mg, 8% yield). $^{1}$H NMR (500 MHz, $CDCl_3$) δ 4.10 (t, *J* = 6.5 Hz, 4H), 3.52 (t, *J* = 5.4 Hz, 2H), 2.57 (t, *J* = 5.4 Hz, 2H), 2.53 – 2.43 (m, 9H), 2.40 (t, *J* = 7.6 Hz, 10H), 2.30 (t, *J* = 7.4 Hz, 4H), 1.77 (p, *J* = 6.7 Hz, 4H), 1.61 (p, *J* = 7.4 Hz, 4H), 1.51 – 1.37 (m, 14H), 1.26 (d, *J* = 5.9 Hz, 61H), 0.87 (t, *J* = 7.0 Hz, 12H). $^{13}$C NMR (126 MHz, $CDCl_3$) δ 173.62, 63.01, 58.59, 55.69, 54.18, 53.65, 50.60, 34.20, 32.04, 29.80, 29.76, 29.75, 29.74, 29.47, 27.67, 26.97, 26.81, 26.38, 22.91, 22.82, 14.25. MS (ESI+): m/z $[M+H]^+$ 936.91 for C58H117N3O5. Purity was >95% by mol as determined by $^{1}$H NMR.

*Synthesis of Compound δO4*

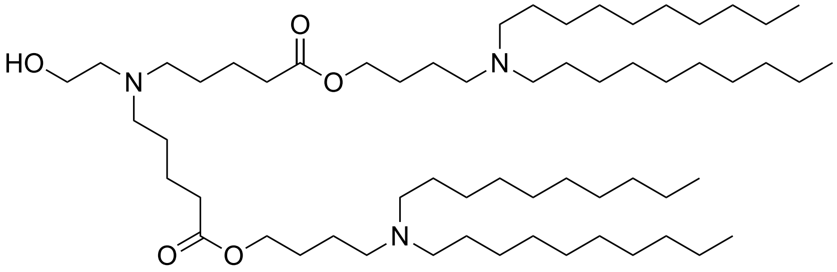


Compound δO4 was synthesized according to General Synthesis Method C using Intermediate J. Product was isolated as a clear oil (100mg, 24% yield). $^1$H NMR (500 MHz, $CDCl_3$) δ 4.05 (t, *J* = 6.7 Hz, 4H), 3.51 (t, *J* = 5.4 Hz, 2H), 2.54 (t, *J* = 5.4 Hz, 2H), 2.47 – 2.34 (m, 16H), 2.29 (t, *J* = 7.4 Hz, 4H), 1.66 – 1.55 (m, 8H), 1.52 – 1.35 (m, 16H), 1.24 (m, 56H), 0.86 (t, *J* = 6.9 Hz, 12H). $^{13}$C NMR (126 MHz, $CDCl_3$) δ 173.56, 64.33, 58.51, 55.56, 54.07, 53.65, 53.52, 50.54, 34.06, 31.88, 29.64, 29.60, 29.57, 29.56, 29.31, 27.60, 26.79, 26.69, 26.67, 23.48, 22.76, 22.65, 14.08. MS (ESI+): m/z $[M+H]^+$ 964.94 for C60H121N3O5. Purity was >90% by mol as determined by $^1$H NMR.

*Synthesis of Compound βN2*

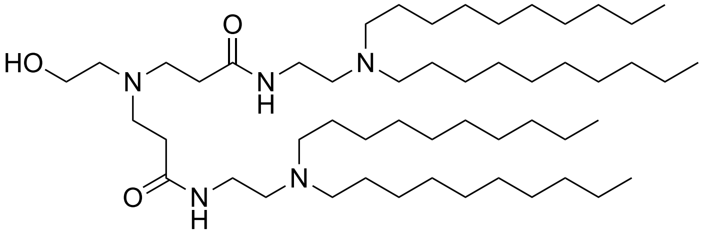


A mixture of Intermediate L (185.3mg, 0.64 mmol, 1 eq), 1-bromodecane (794µL, 3.84 mmol, 6 eq) and DIPEA (446µL, 2.56 mmol, 4 eq) were dissolved in ethanol (2mL) in a screw-cap vial. The mixture was covered with foil and stirred at 65°C for 3 days. The mixture was then concentrated under reduced pressure and re-suspended in ethyl acetate to precipitate DIPEA-HBr salts, then filtered and washed thoroughly with ethyl acetate. The filtrate was collected and concentrated to a crude oil, then purified by column chromatography using a 10-40% (22% MeOH, 3% $NH_4OH$ in dichloromethane) in dichloromethane gradient. $^1$H NMR (500 MHz, $CDCl_3$) δ 7.34 (br s, 2H), 3.58 (dd, *J* = 5.9, 3.8 Hz, 2H), 3.31 (q, *J* = 5.9 Hz, 4H), 2.75 (t, *J* = 6.4 Hz, 4H), 2.62 (br s, 2H), 2.55 (t, *J* = 4.8 Hz, 2H), 2.49 (t, *J* = 5.3 Hz, 6H), 2.34 (t, *J* = 6.4 Hz, 4H), 1.44 (p, *J* = 7.5 Hz, 8H), 1.31 – 1.18 (m, 58H), 0.86 (t, *J* = 7.1 Hz, 12H). $^{13}$C NMR (126 MHz, $CDCl_3$) δ 172.48, 59.17, 56.12, 53.77, 53.14, 50.08, 36.59, 33.93, 32.00, 29.77, 29.71, 29.68, 29.64, 29.43, 27.59, 25.92, 22.78, 14.21. MS (ESI+): m/z $[M+H]^+$ 850.84 for C52H107N5O3. Purity was >95% by mol as determined by $^1$H NMR.

### Supplementary Figures

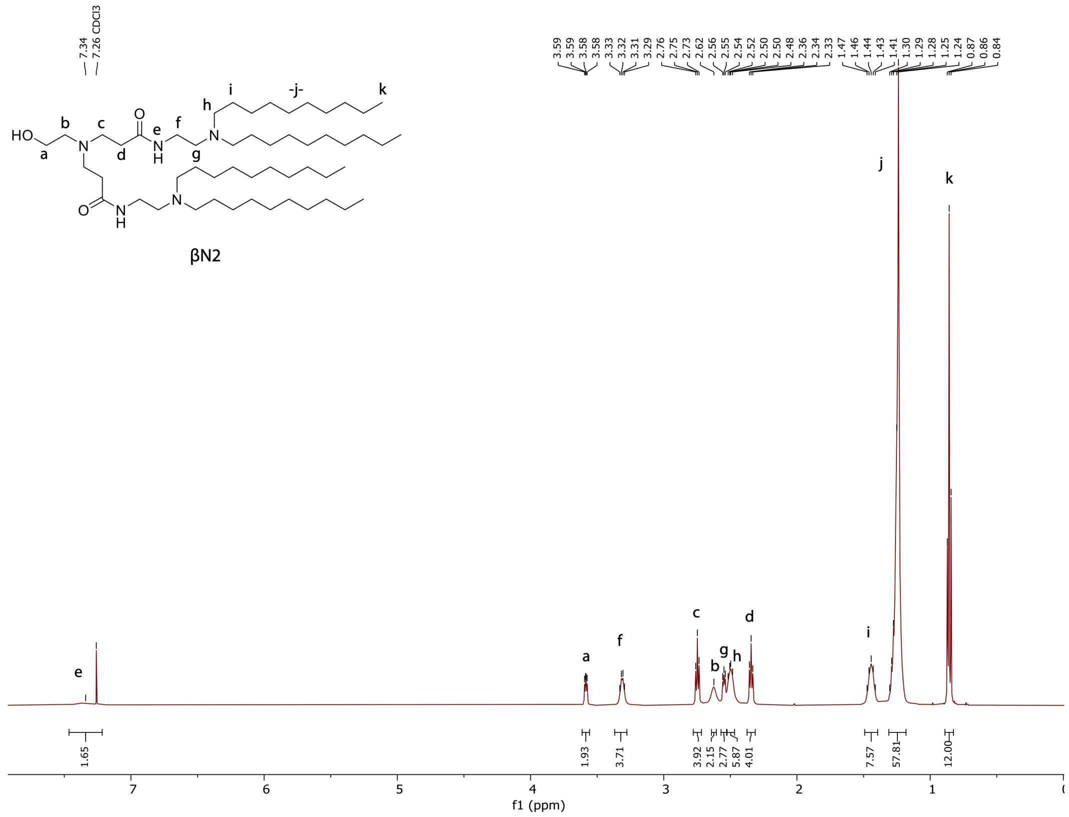


**Figure S1. $^{1}H$ NMR spectrum of βN2.**

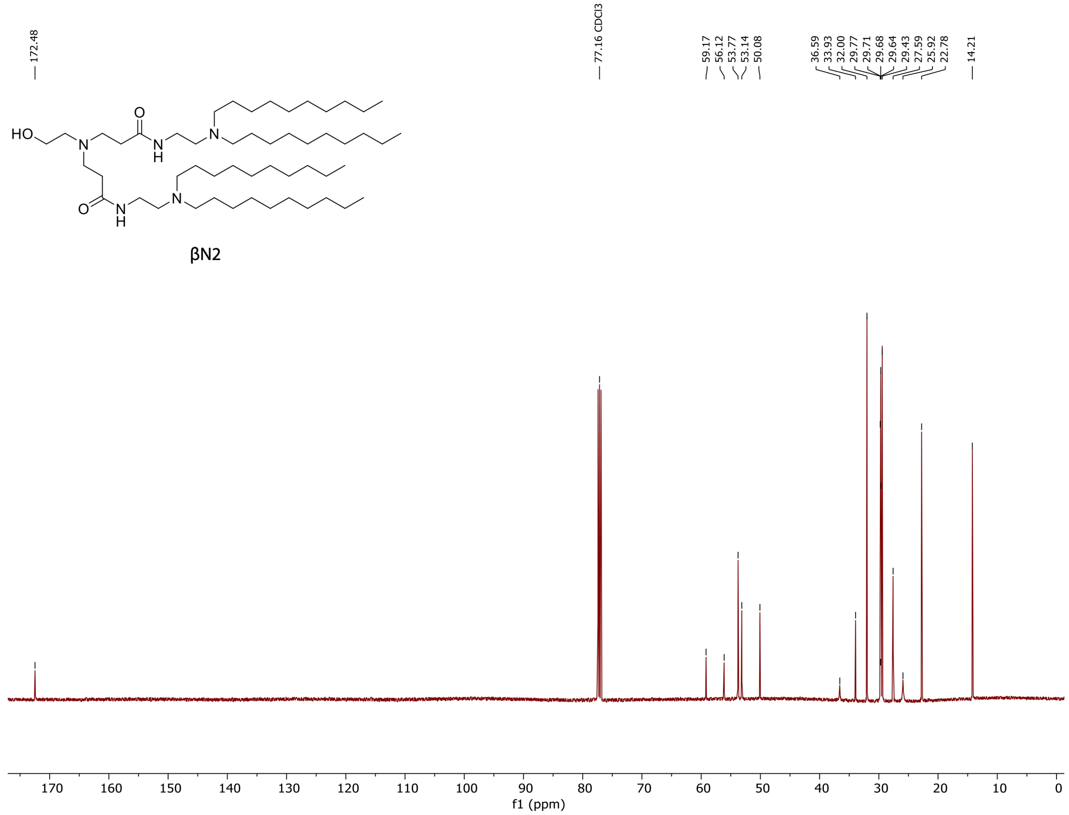


**Figure S2. $^{13}C\{^{1}H\}$ NMR spectrum of βN2.**

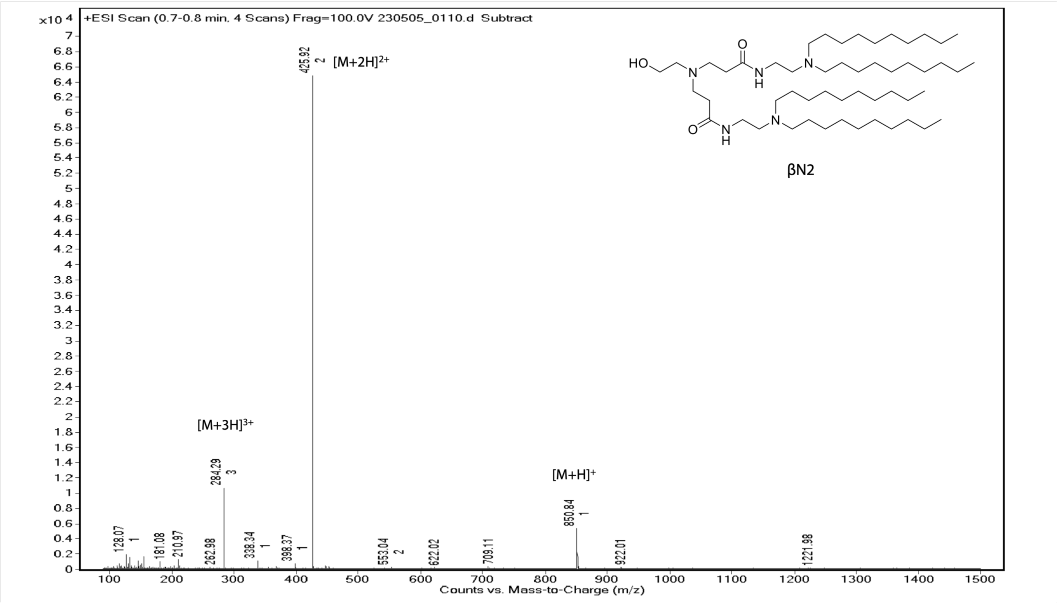


**Figure S3. ESI (positive mode) mass spectrum of βN2.**

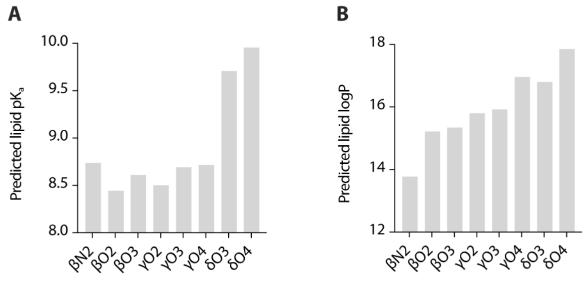


**Figure S4. Marvin predictions of ionizable lipid pK$_a$ and logP. A,** Predicted lipid pK$_a$. **B,** Predicted logP.

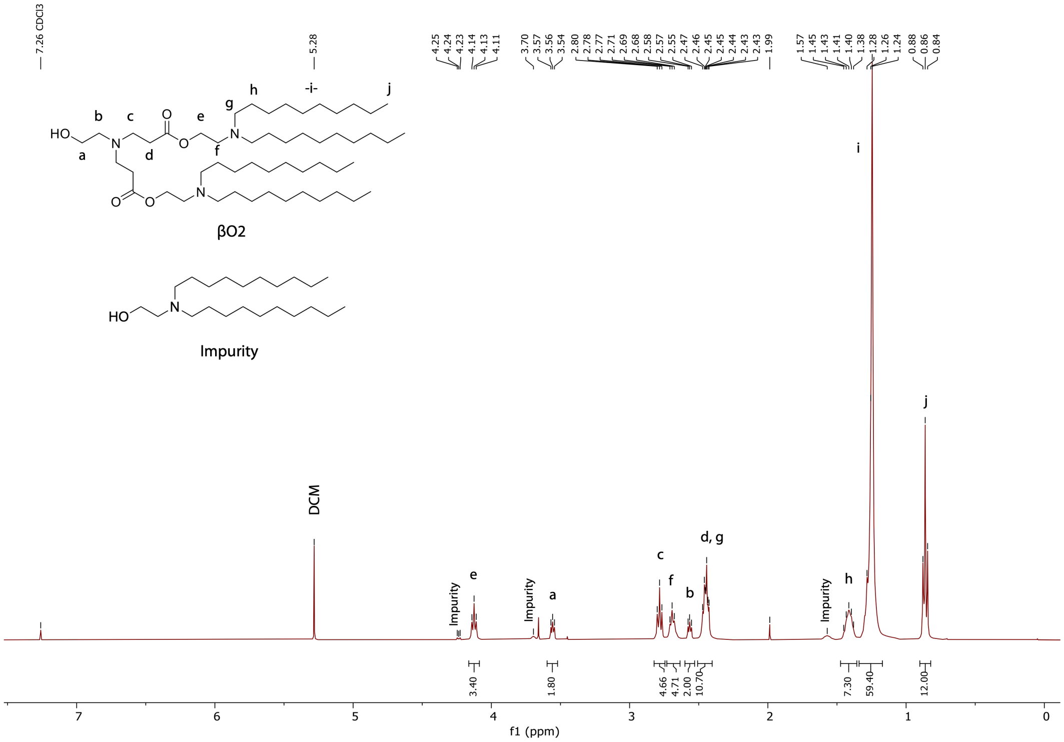


**Figure S5. $^1$H NMR spectrum of βO2.**

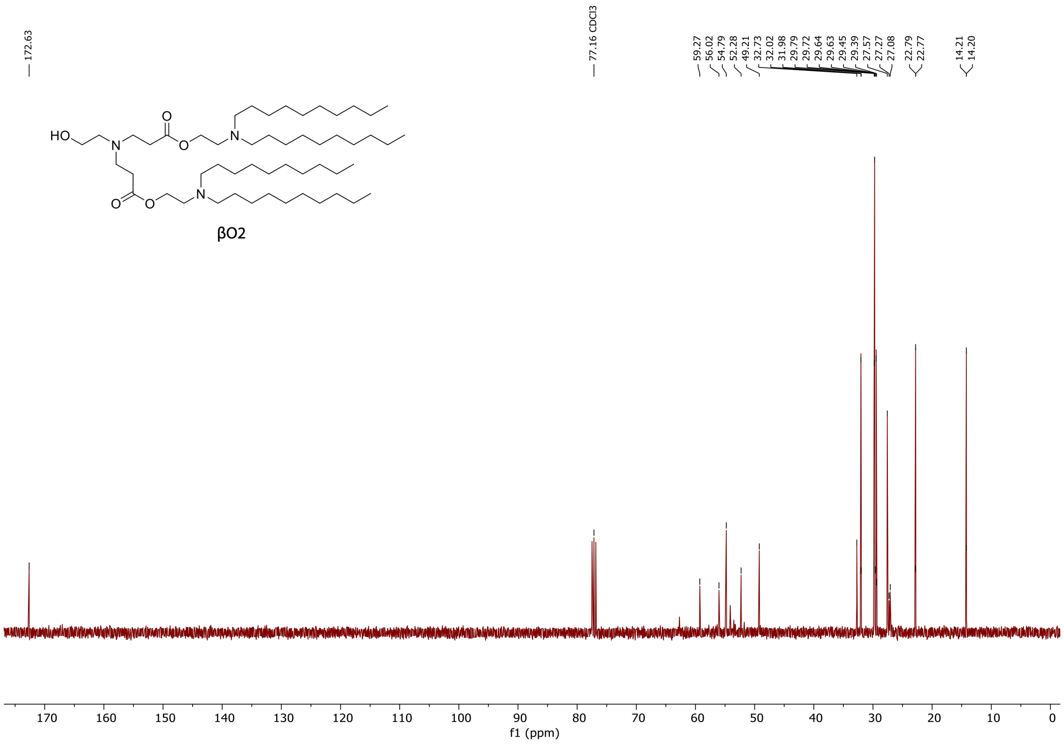


**Figure S6. $^{13}C\{^1H\}$ NMR spectrum of βO2.**

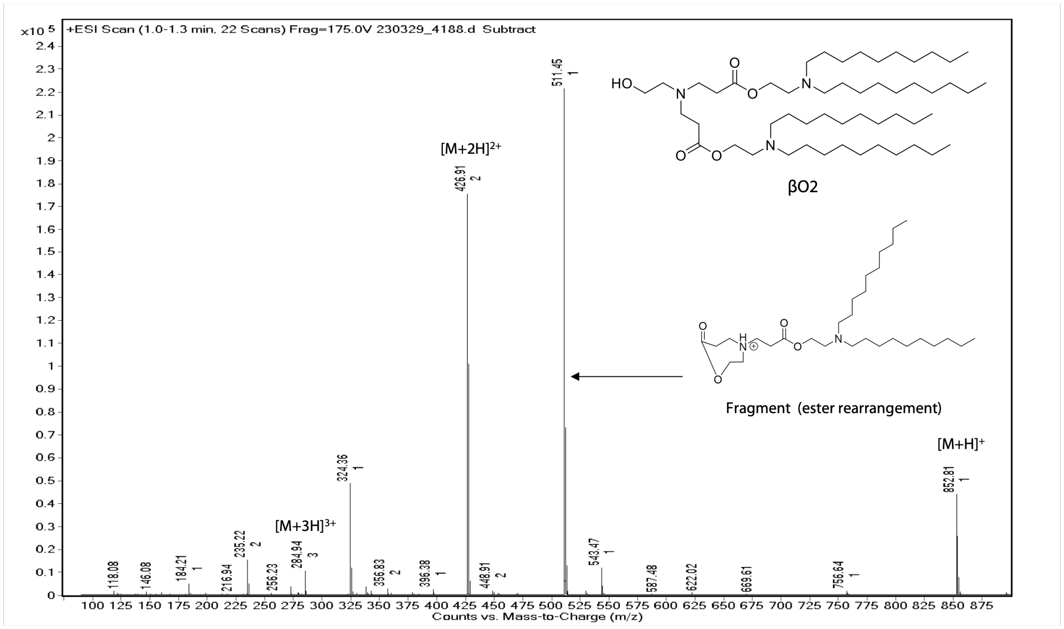


**Figure S7. ESI (positive mode) mass spectrum of βO2.**

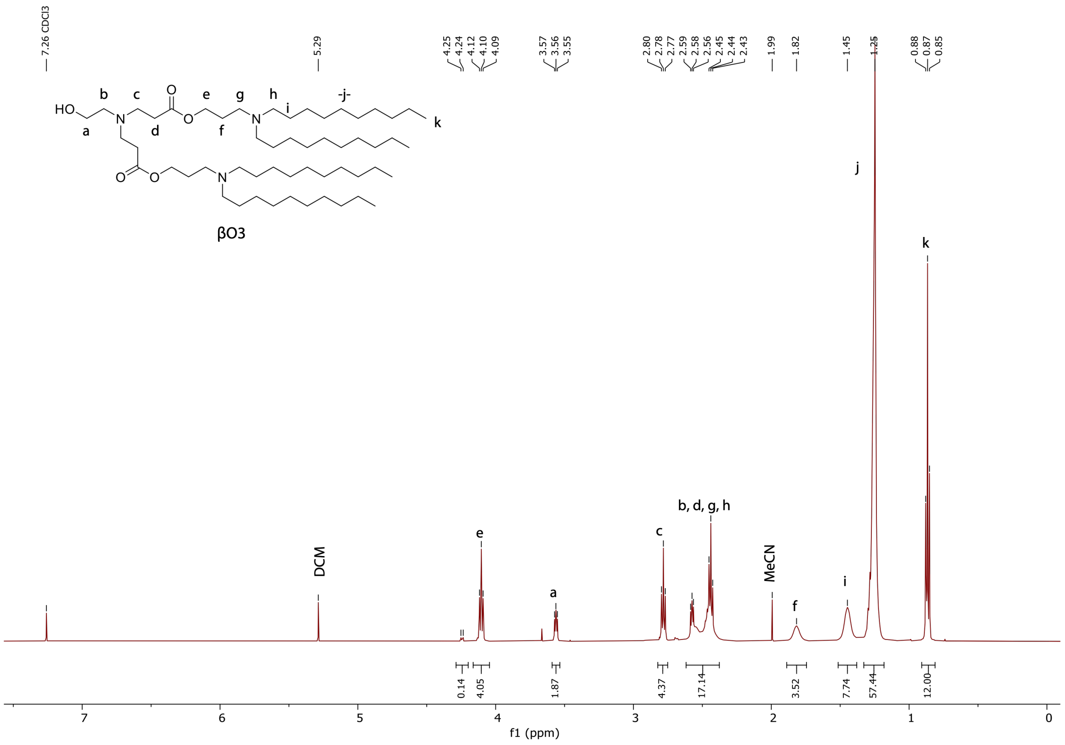

**Figure S8. $^{1}H$ NMR spectrum of βO3.**

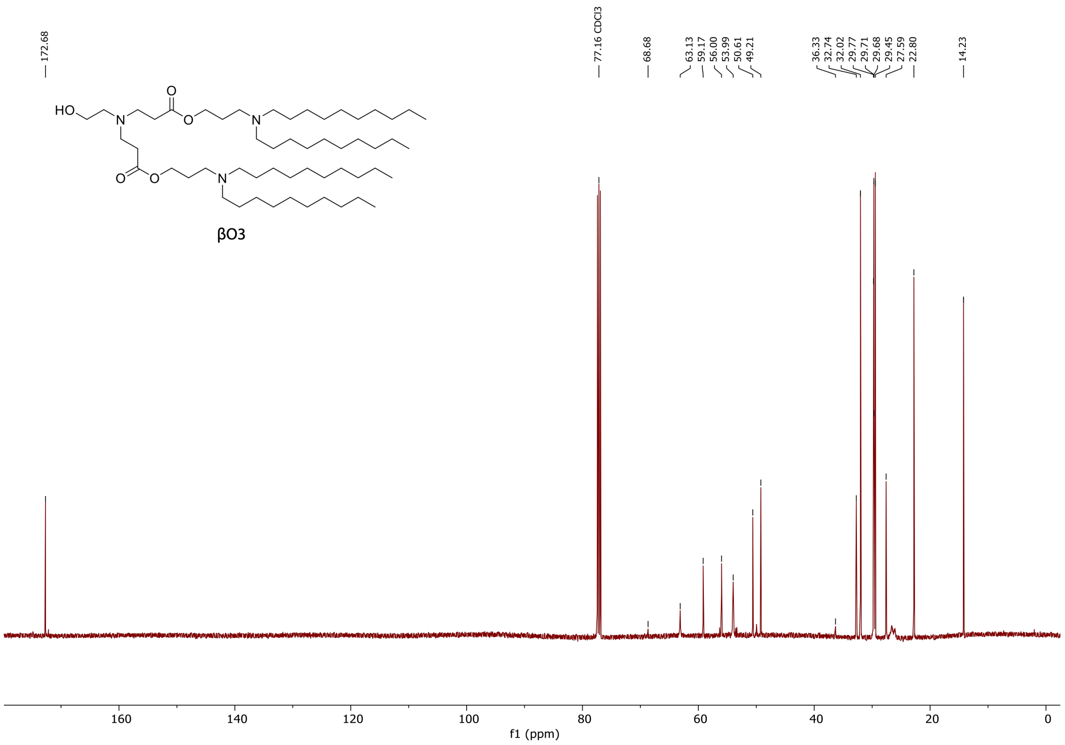

**Figure S9. $^{13}C\{^{1}H\}$ NMR spectrum of βO3.**

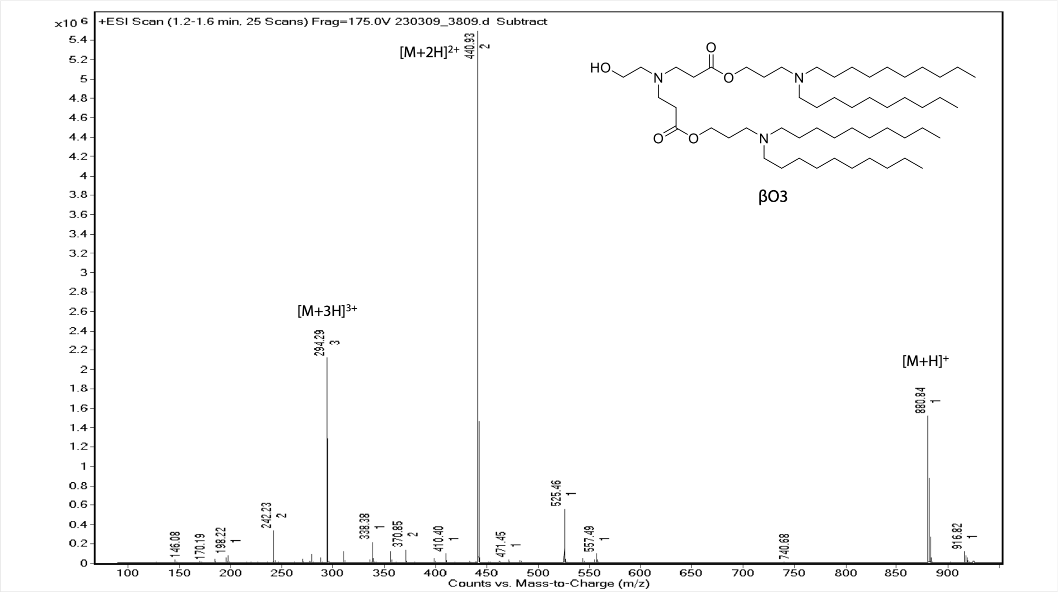


**Figure S10. ESI (positive mode) mass spectrum of βO3.**

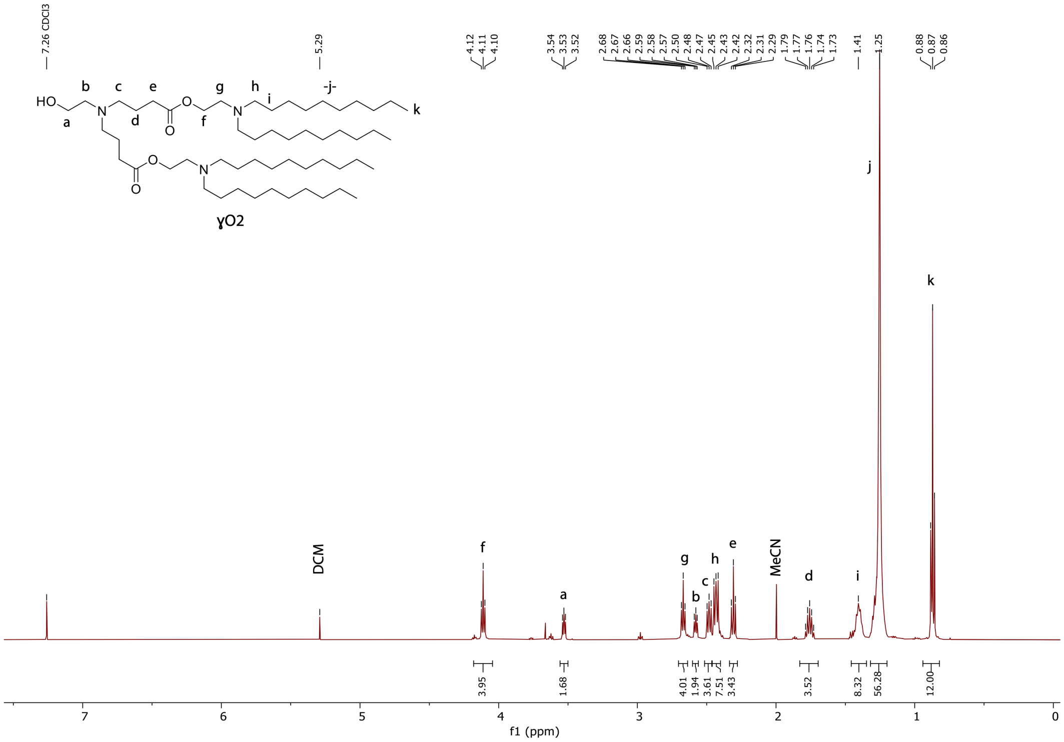


**Figure S11. $^1$H NMR spectrum of ɣO2.**

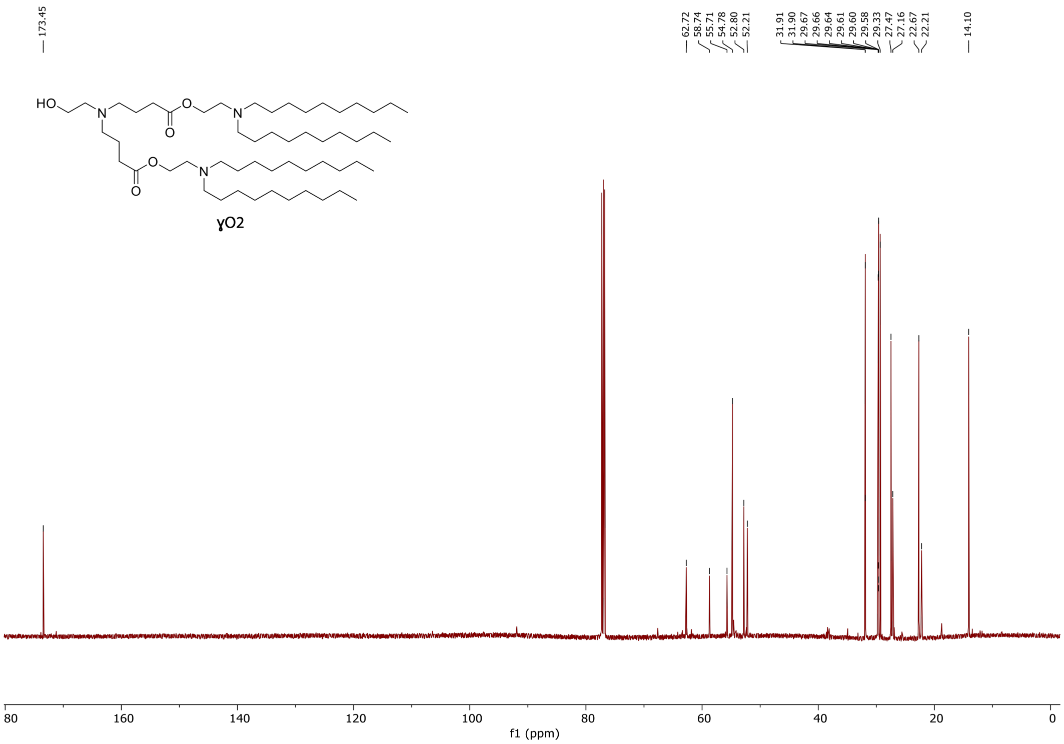


**Figure S12. $^{13}C\{^{1}H\}$ NMR spectrum of ɣO2.**

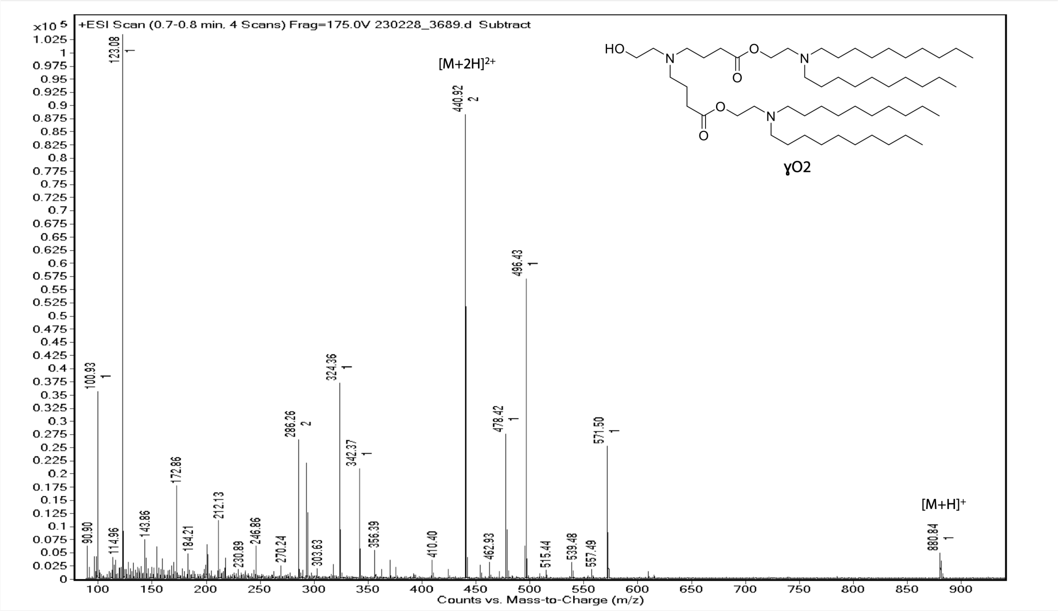


**Figure S13. ESI (positive mode) mass spectrum of ɣO2.** Peaks at 571.50, 496.43, 342.37, and 324.36 m/z were identified as potential fragments induced by ionization.

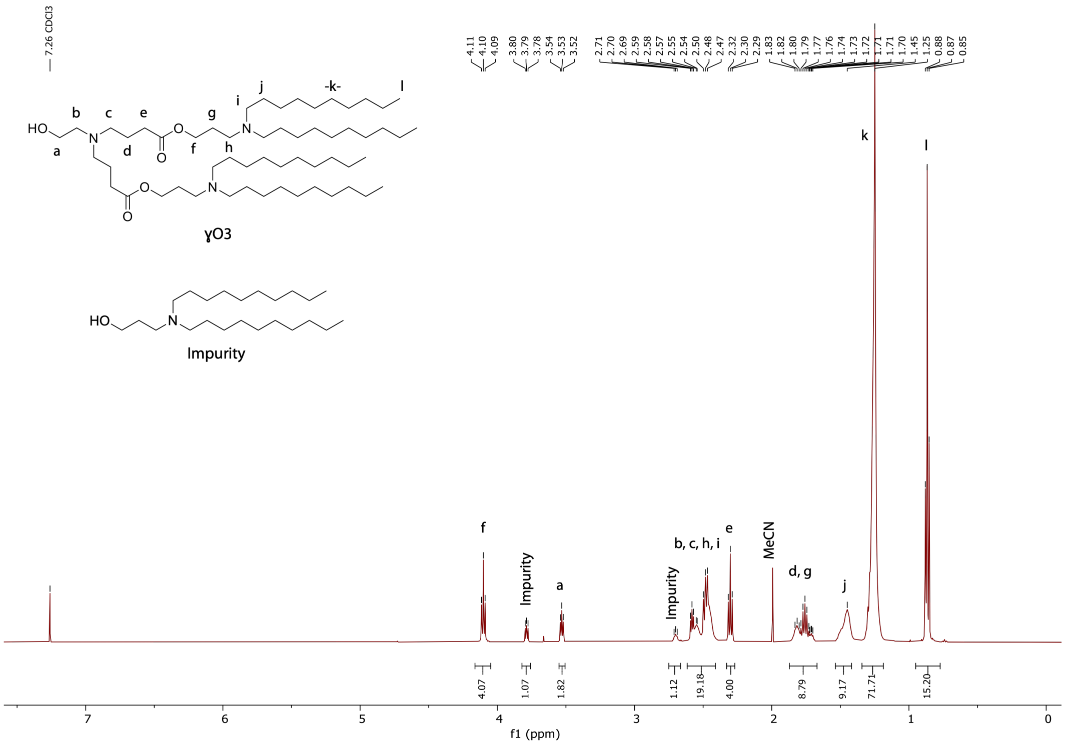


**Figure S14.** $^{1}$**H NMR spectrum of ɣO3.**

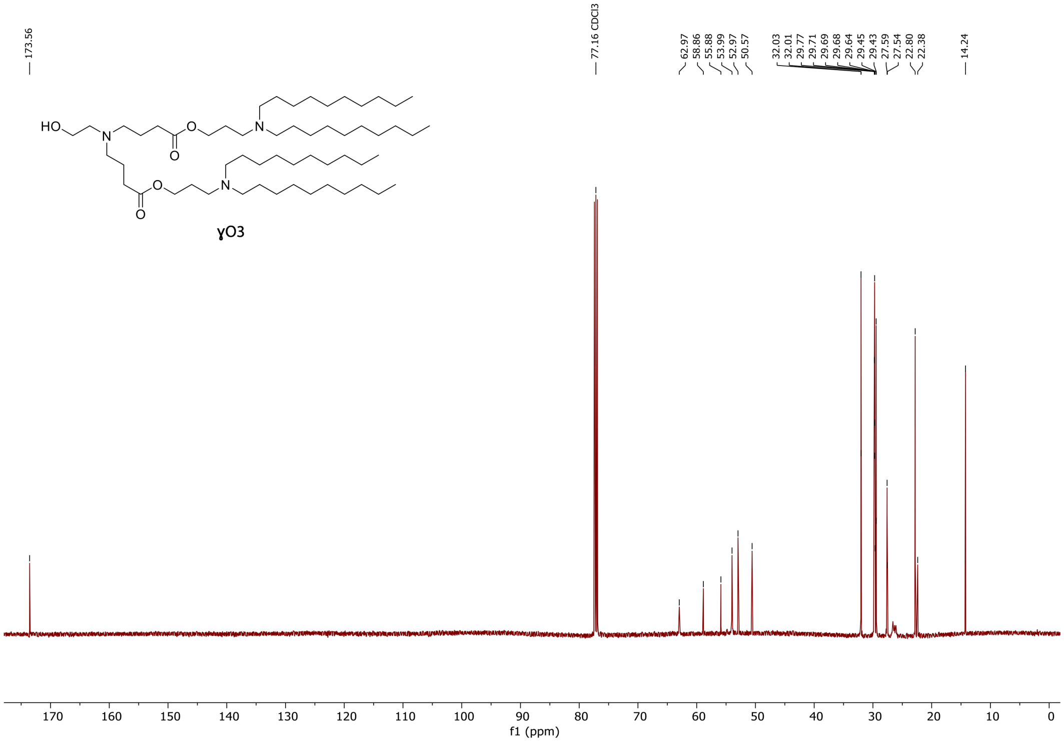


**Figure S15.** $^{13}$**C{**$^{1}$**H} NMR spectrum of ɣO3.**

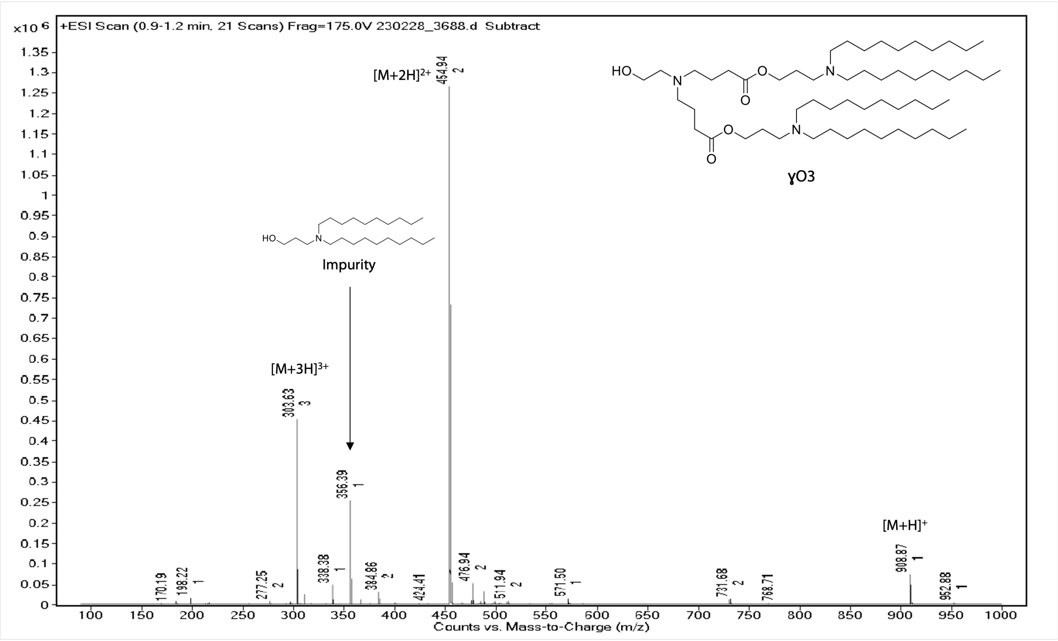


**Figure S16. ESI (positive mode) mass spectrum of ɣO3.**

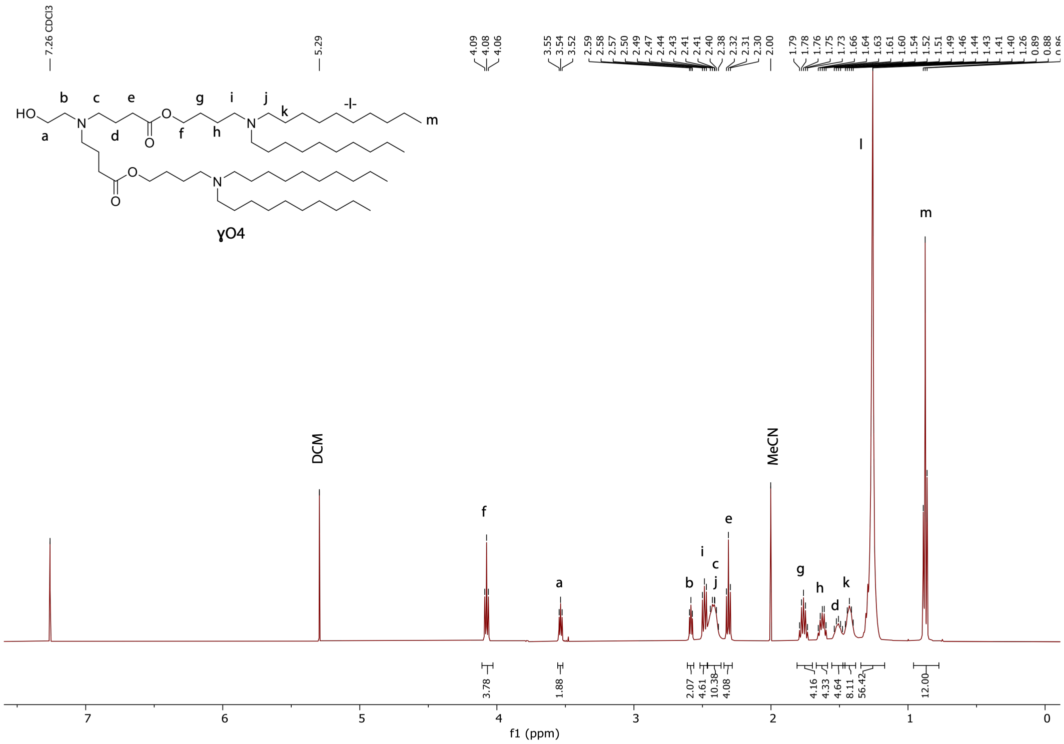


**Figure S17. $^{1}$H NMR spectrum of ɣO4.**

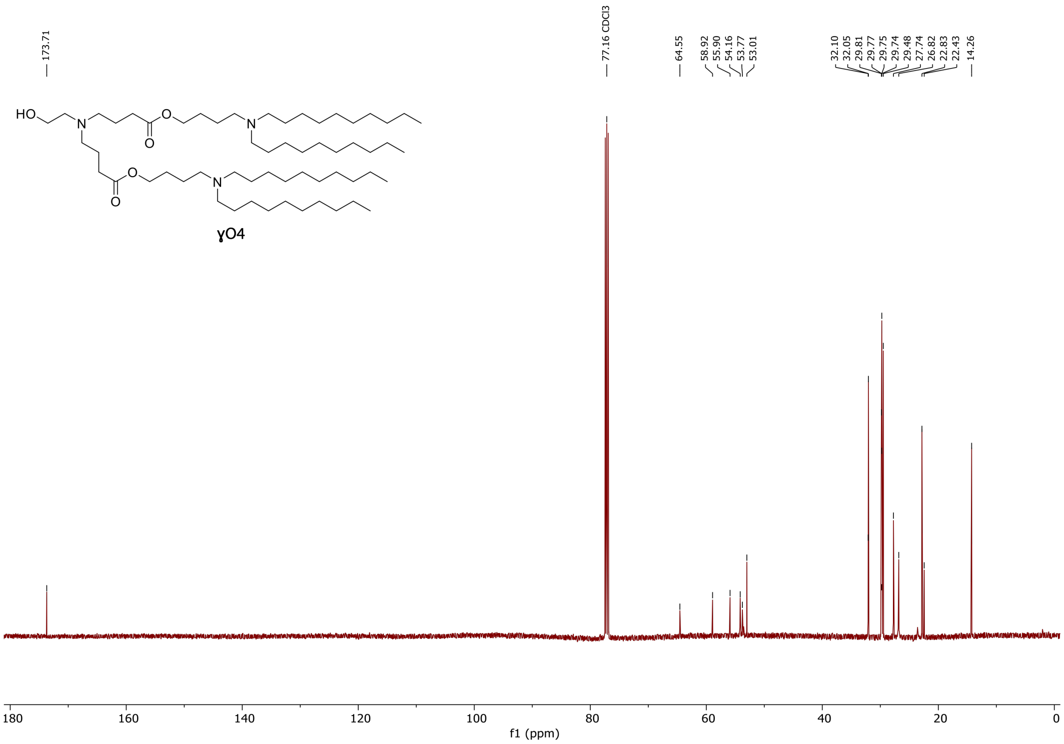


**Figure S18.** $^{13}C\{^{1}H\}$ **NMR spectrum of ɣO4.**

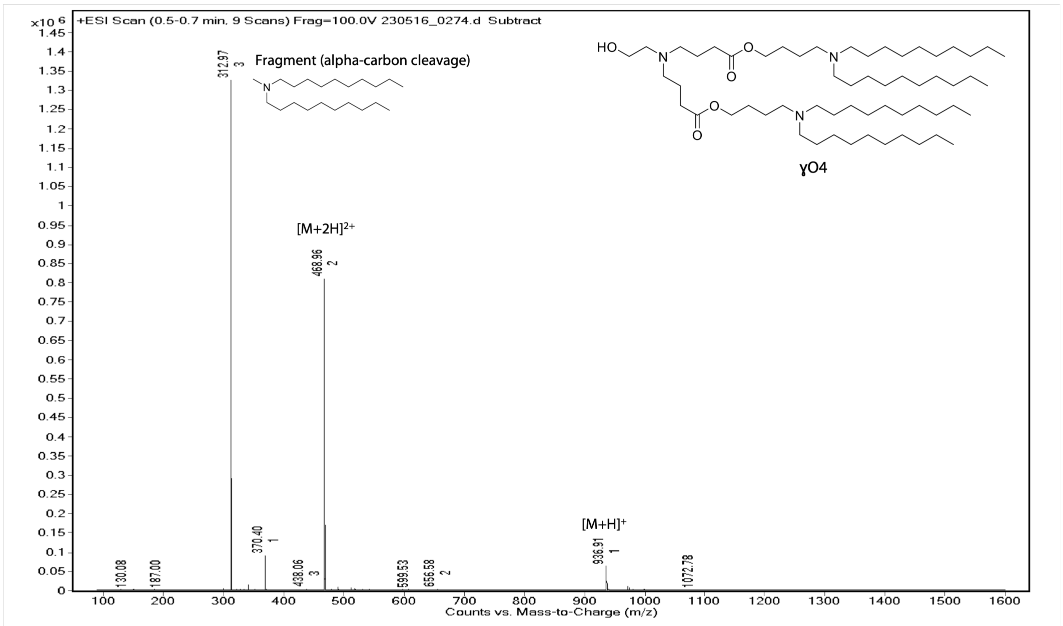


**Figure S19. ESI (positive mode) mass spectrum of ɣO4.**

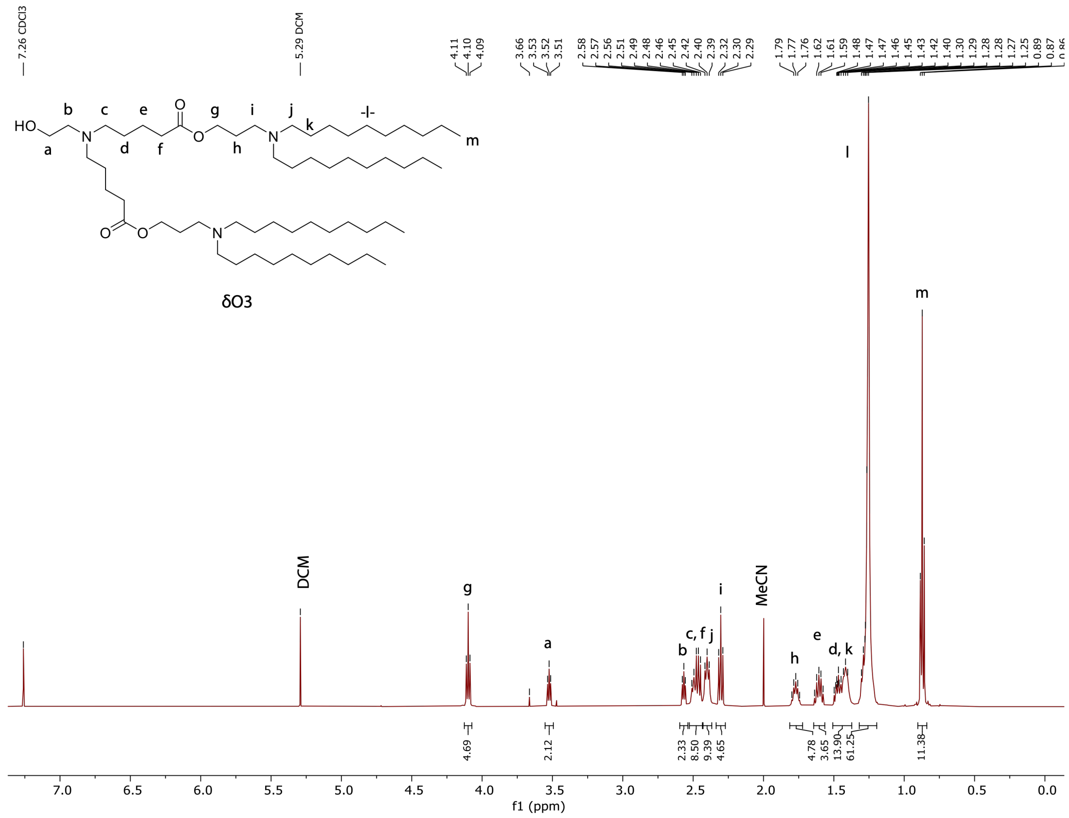

**Figure S20. $^{1}H$ NMR spectrum of δO3.**

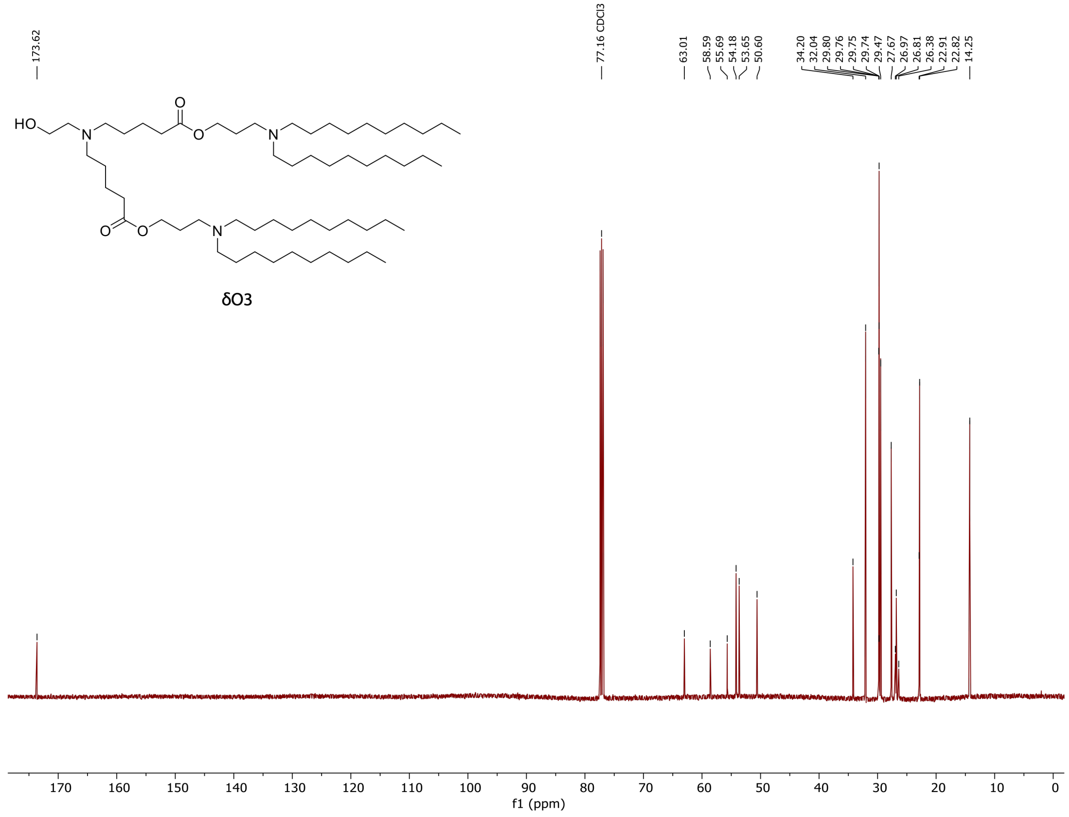

**Figure S21. $^{13}C\{^{1}H\}$ NMR spectrum of δO3.**

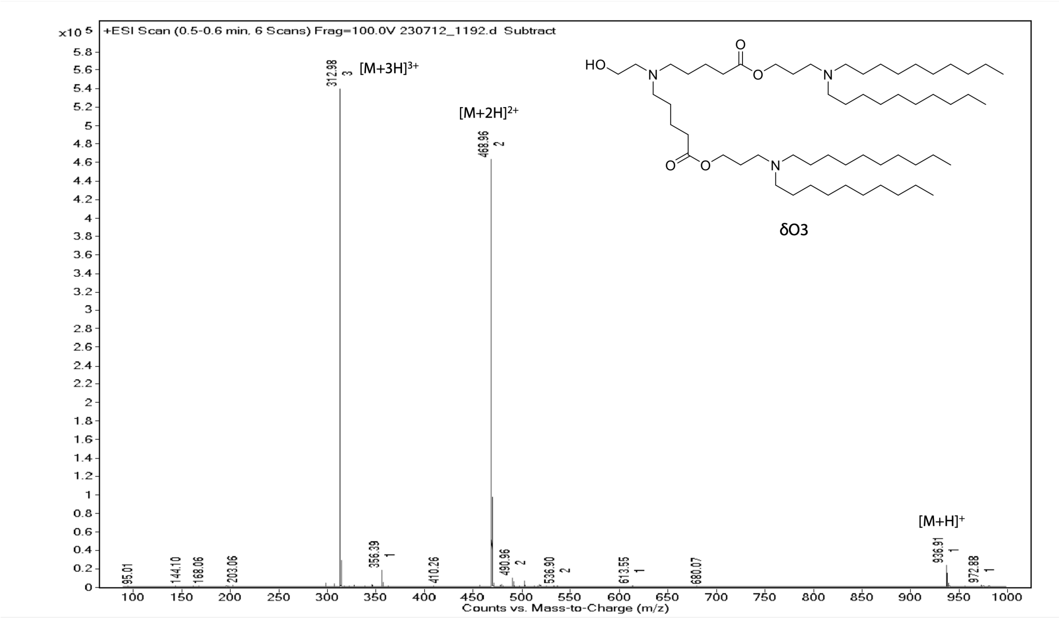


**Figure S22. ESI (positive mode) mass spectrum of δO3.**

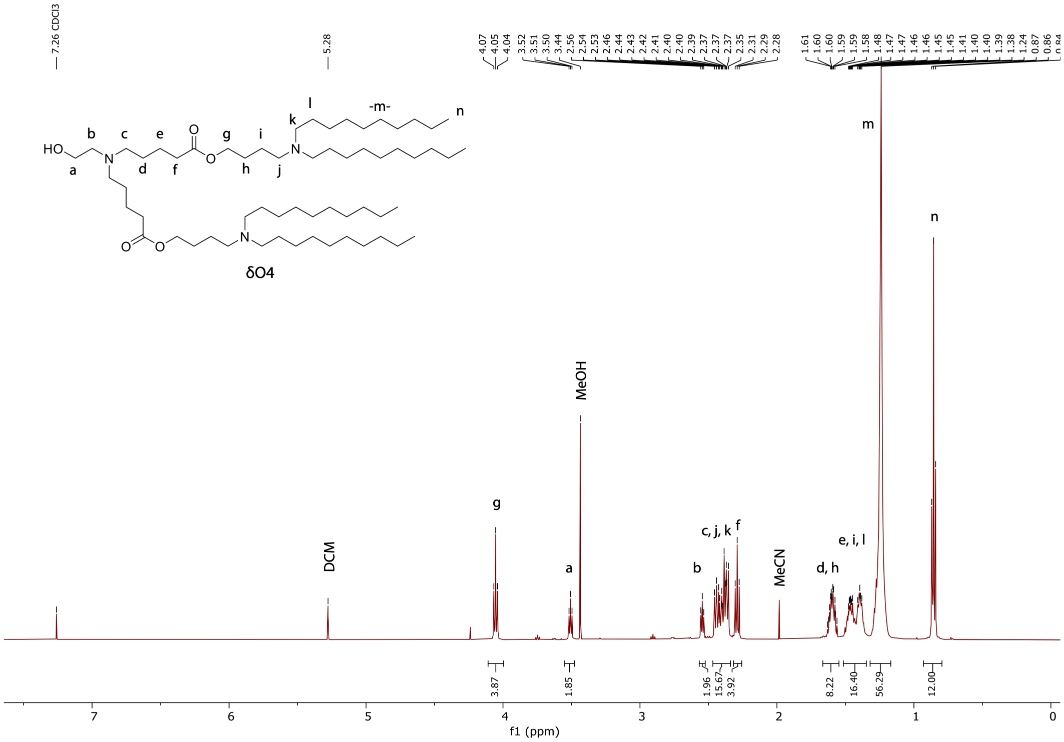


**Figure S23. $^{1}$H NMR spectrum of δO4.**

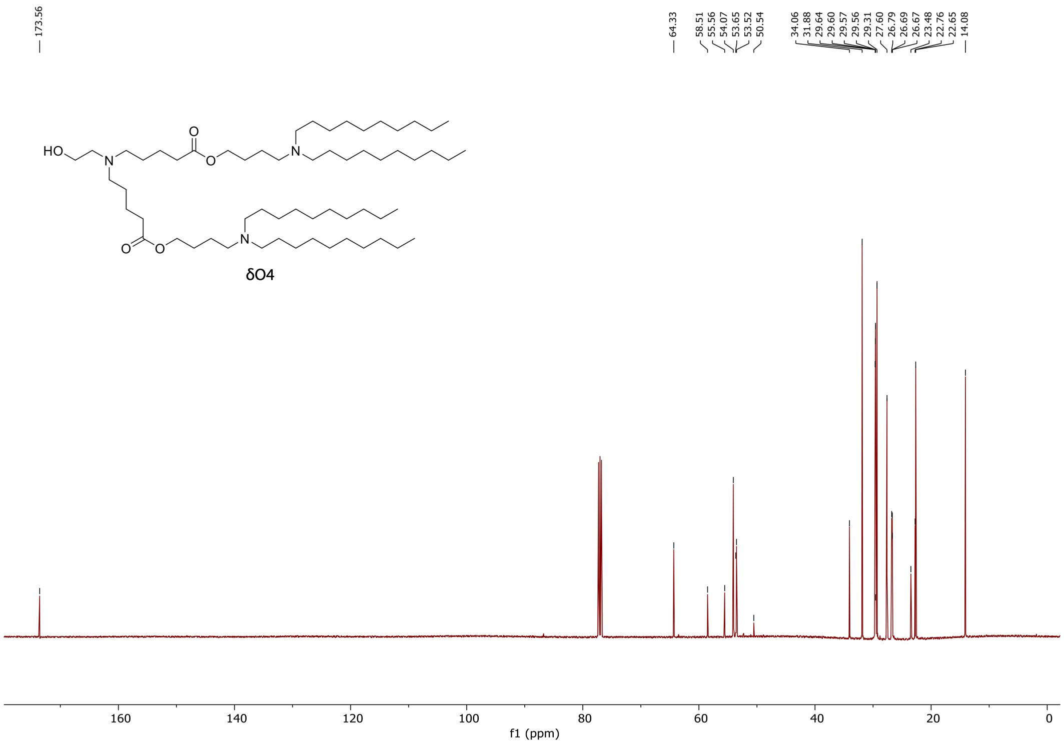


**Figure S24.** $^{13}C\{^1H\}$ **NMR spectrum of δO4.**

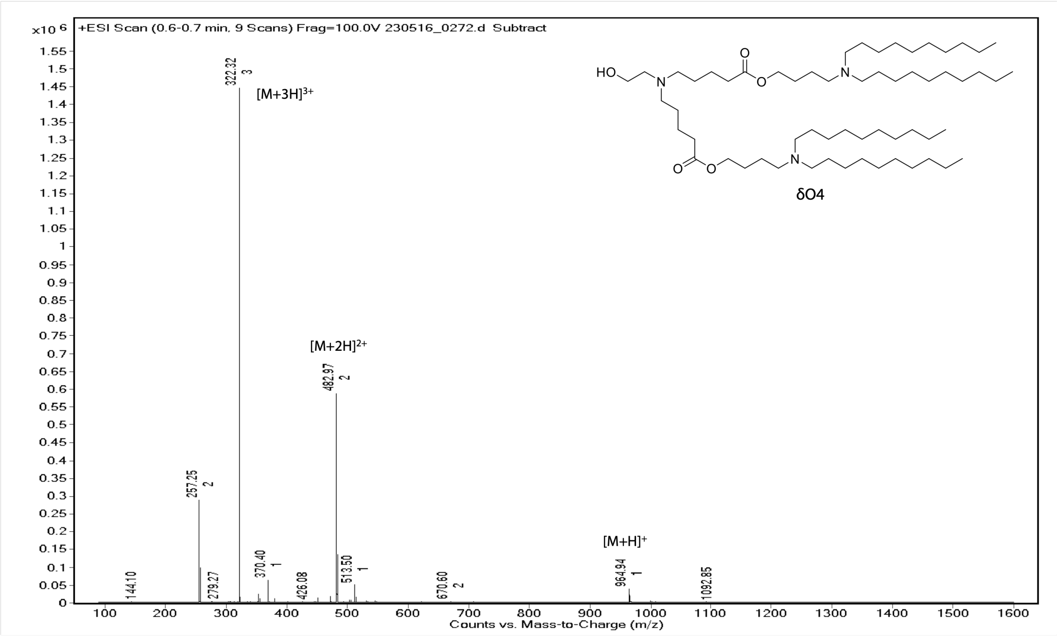


**Figure S25. ESI (positive mode) mass spectrum of δO4.**

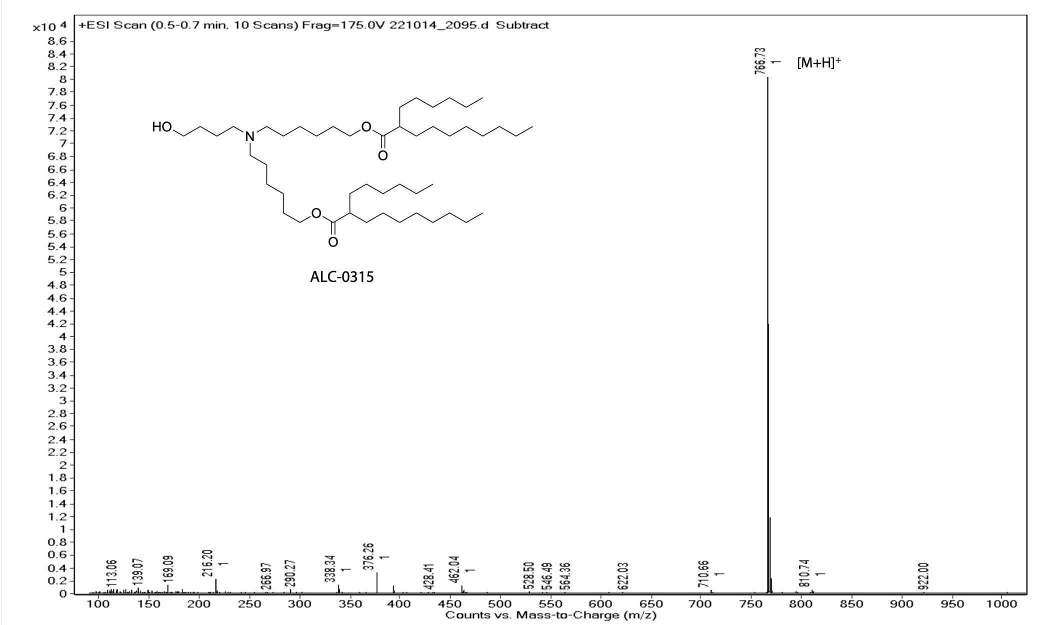


**Figure S26. ESI (positive mode) mass spectrum of ALC-0315.**

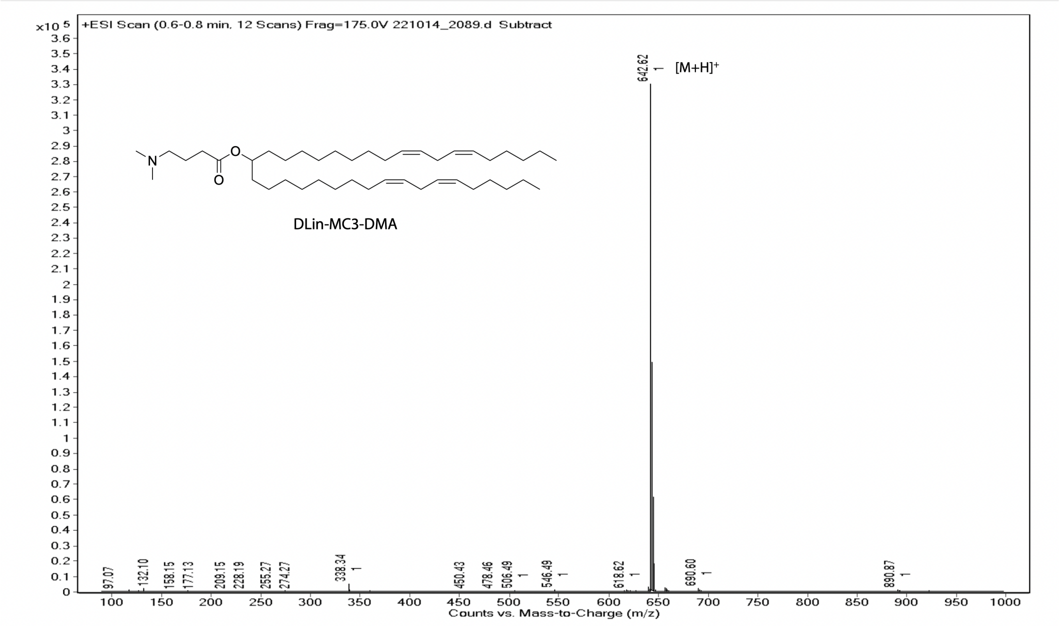


**Figure S27. ESI (positive mode) mass spectrum of DLin-MC3-DMA.**

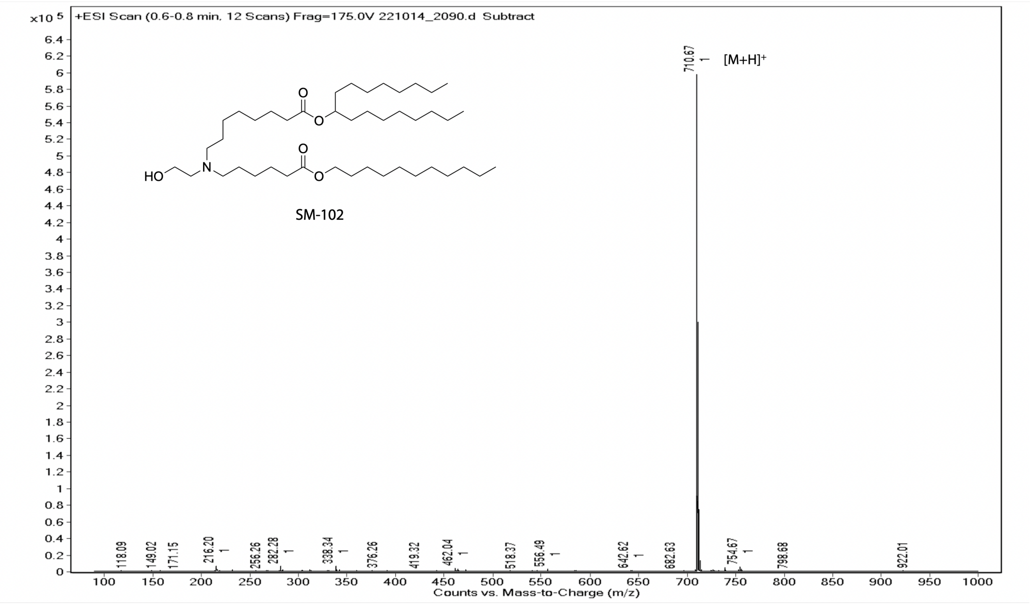


**Figure S28. ESI (positive mode) mass spectrum of SM-102.**

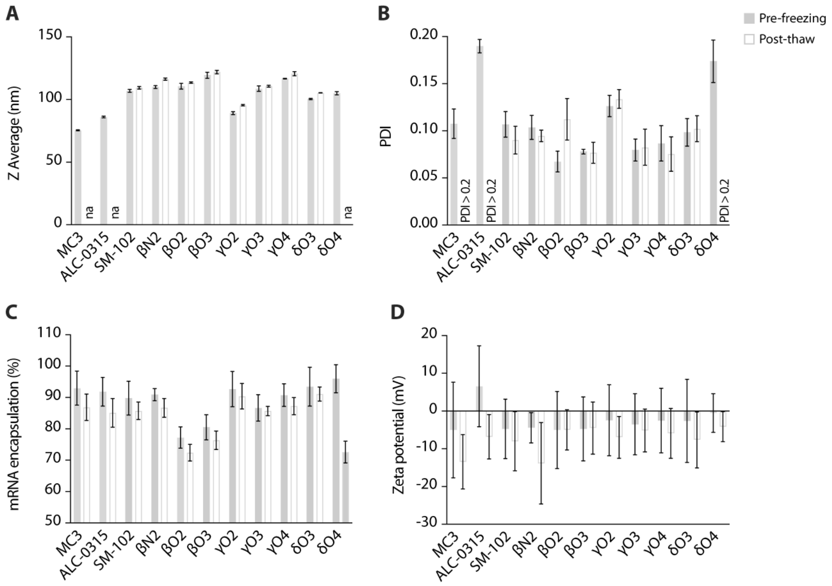


**Figure S29. LNP physicochemical parameters pre-freezing and post-thaw. A,** Z Average size. **B,** PDI. **C,** mRNA encapsulation efficiency. **D,** Zeta potential. Data and error bars indicate the mean and standard deviation (n = 3 technical replicates). na denotes not applicable.

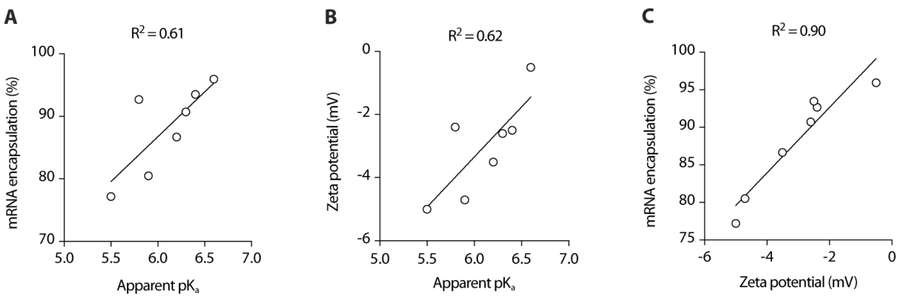


**Figure S30. Relationship between LNP physicochemical parameters of ester analogs. A,** Linear correlation between mRNA encapsulation efficiency and apparent $pK_a$. **B,** Linear correlation between zeta potential and LNP apparent $pK_a$. **C,** Linear correlation between mRNA encapsulation efficiency and zeta potential.

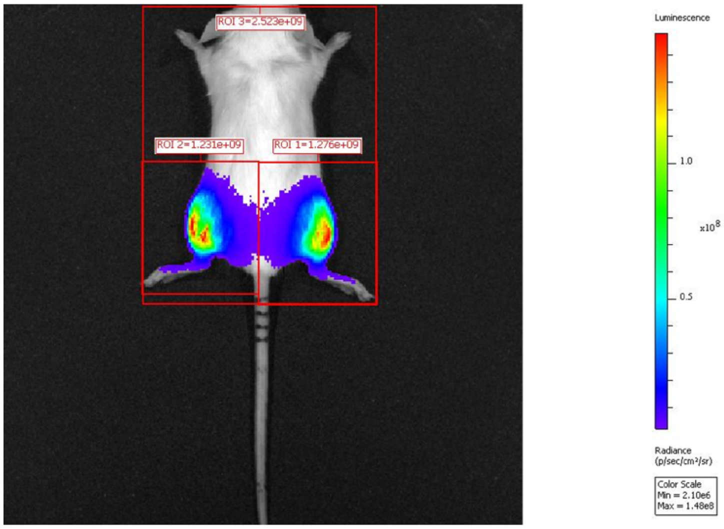


**Figure S31. Representative image of a total luminescent flux measurement in BALB/c mice.** Red boxes indicate the region of interest (ROI).

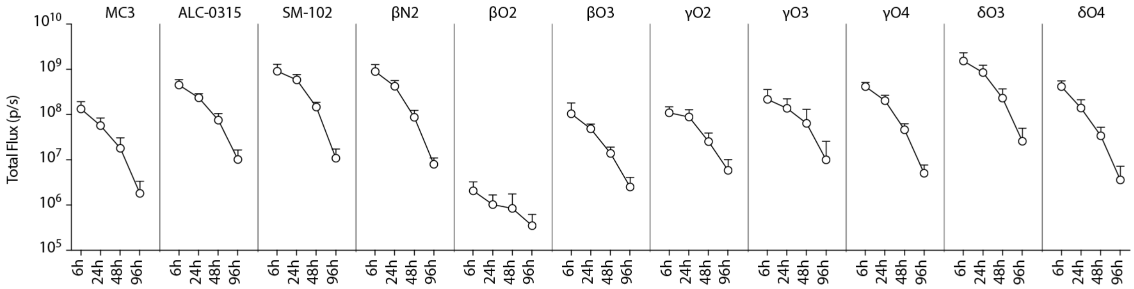


**Figure S32. Intramuscular expression profile.** Total luminescent flux in muscle after intramuscular injection of 0.5 µg FLuc mRNA-LNPs containing different ionizable lipids. Data and error bars indicate the mean and standard deviation (n = 4 mice, 8 hindlimbs).

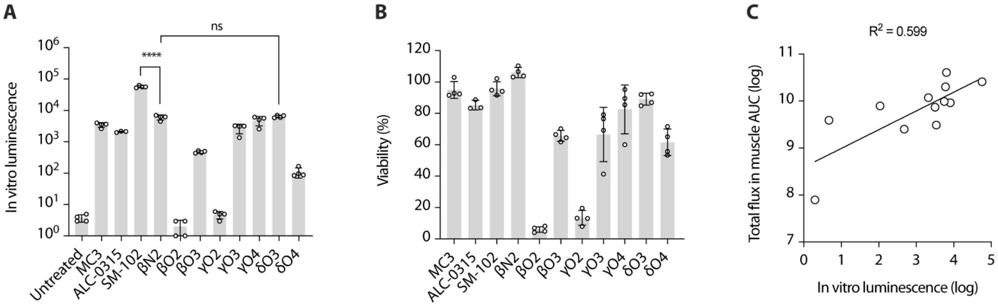


**Figure S33. In vitro screening of ionizable lipids with THP-1 monocytes. A,** In vitro luminescence. **B,** Cell viability. **C,** Linear correlation between mean total luminescent flux in BALB/c muscle and mean in vitro luminescence. Data and error bars indicate the mean and standard deviation (n = 4 wells). P-values were determined by one-way ANOVA with Bonferroni's multiple comparisons test.

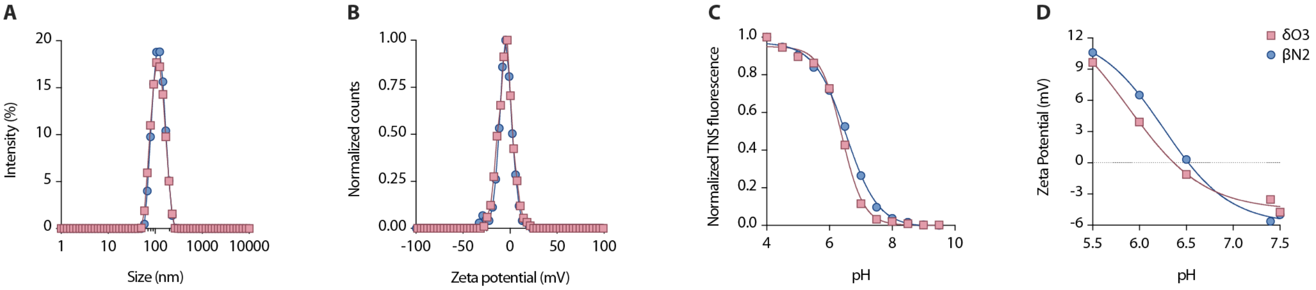


**Figure S34. Physicochemical characterization of hEPO mRNA-LNPs. A,** Size distribution. **B,** Zeta potential distribution. **C,** TNS assay with curve fit. **D,** Zeta potential at different pH (100 mM phosphate buffer). Data for βN2 are indicated by blue circles, and data for δO3 are indicated by red squares. Data indicate the mean of technical replicates.

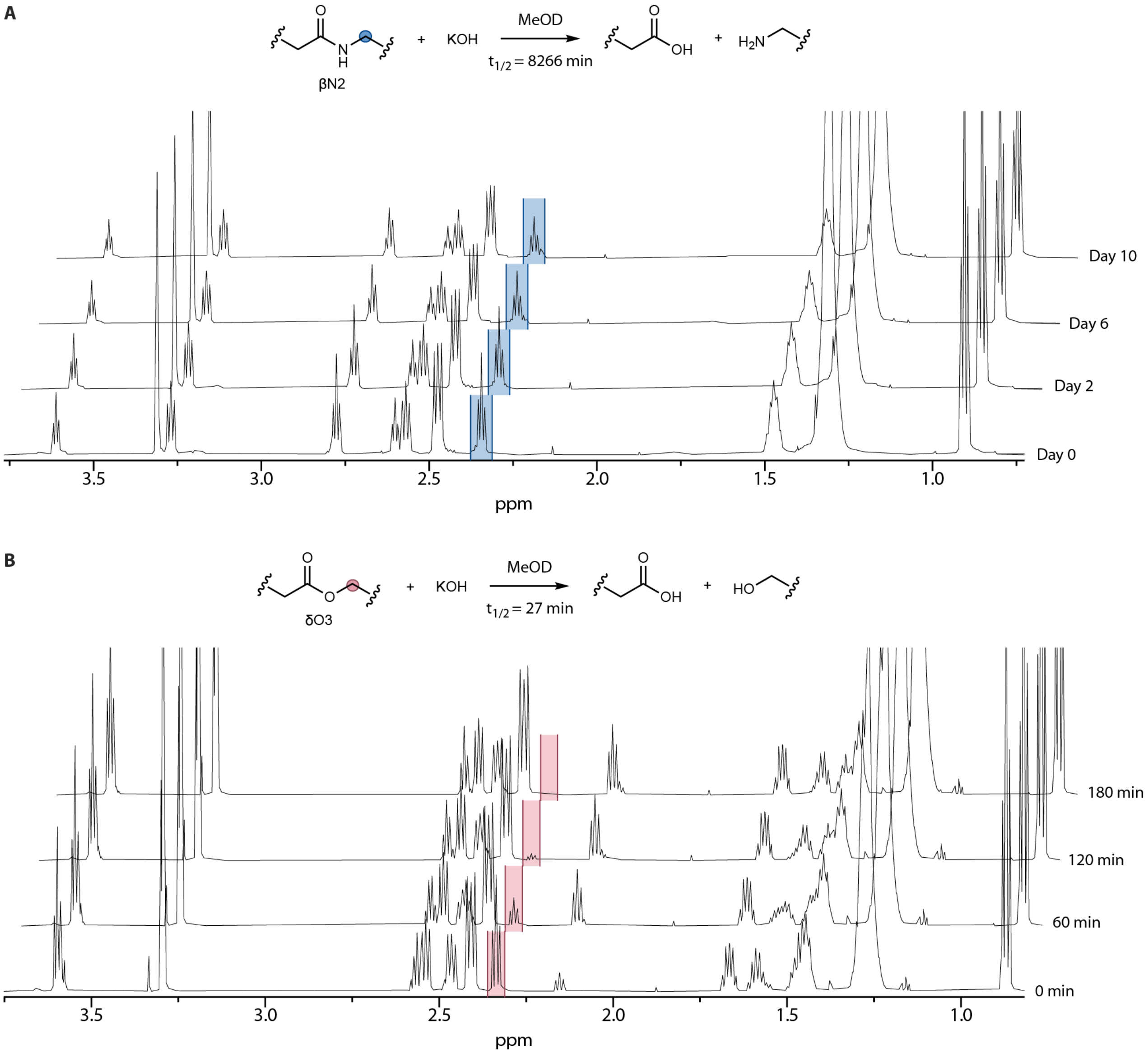


**Figure S35. Base hydrolysis of ionizable lipids.** $^{1}$H-NMR stack plot of (A) βN2 and (B) δO3 reacting with potassium hydroxide (KOH) in deuterated methanol (MeOD) over time. The characteristic signal of the parent lipid used for integration is highlighted in blue for βN2 and red for δO3.

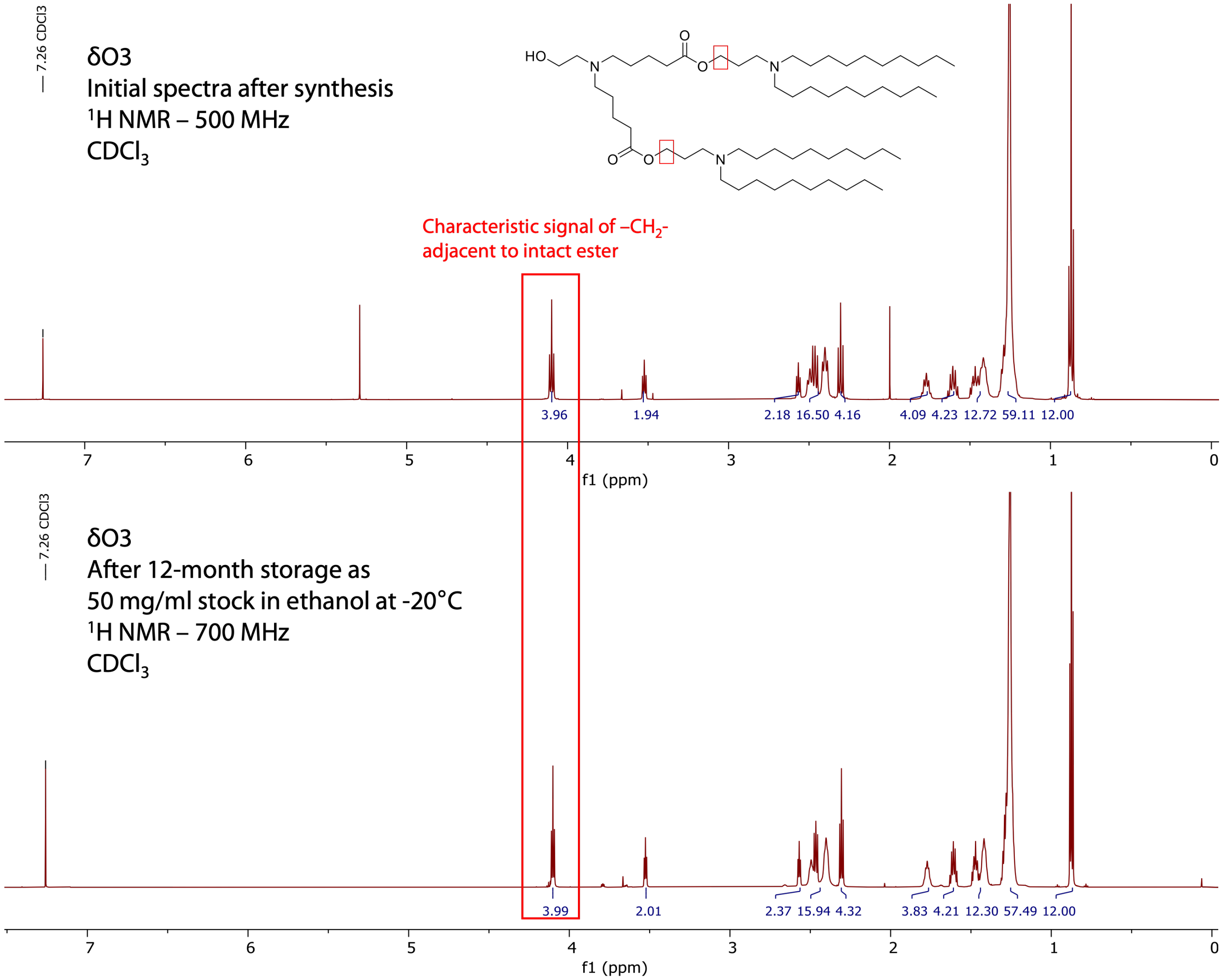


**Figure S36. Comparison of $^1$H NMR spectra of δO3 immediately after synthesis, and after -20°C storage for one year as 50 mg/mL stock in ethanol.**

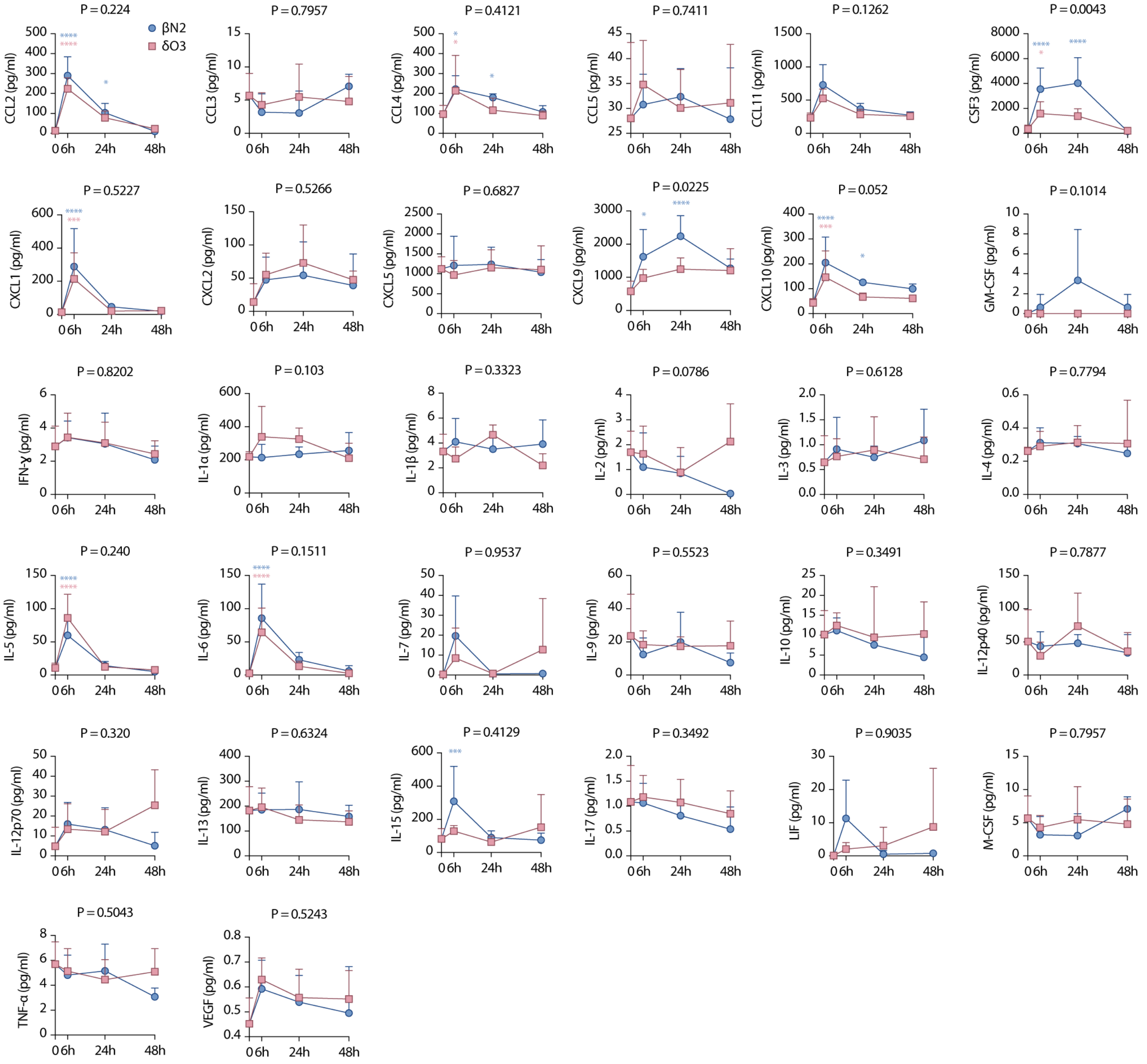


**Figure S37. Cytokine profile to mRNA-LNPs**. BALB/c mice were injected intramuscularly with mRNA-LNPs containing βN2 or δO3. Cytokines were measured in serum 6, 24, and 48 hours by Luminex assay. Data for βN2 are indicated by blue circles, and data for δO3 are indicated by red squares. Data and error bars indicate the mean and standard deviation across samples from independent mice (n = 4, terminal timepoints). P-values were determined by two-way ANOVA with Bonferroni's multiple comparisons test. P values above graphs represent the ionizable lipid factor. P values above data points represent the comparison with baseline.

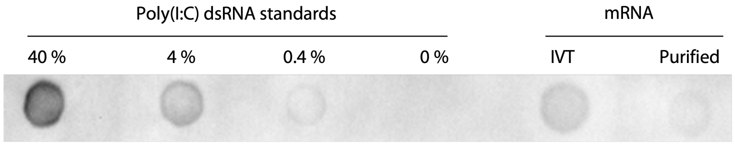


**Figure S38. Representative dot blot using J2 dsRNA-specific antibody of mRNA synthesized by in vitro transcription (IVT) after cellulose purification.**

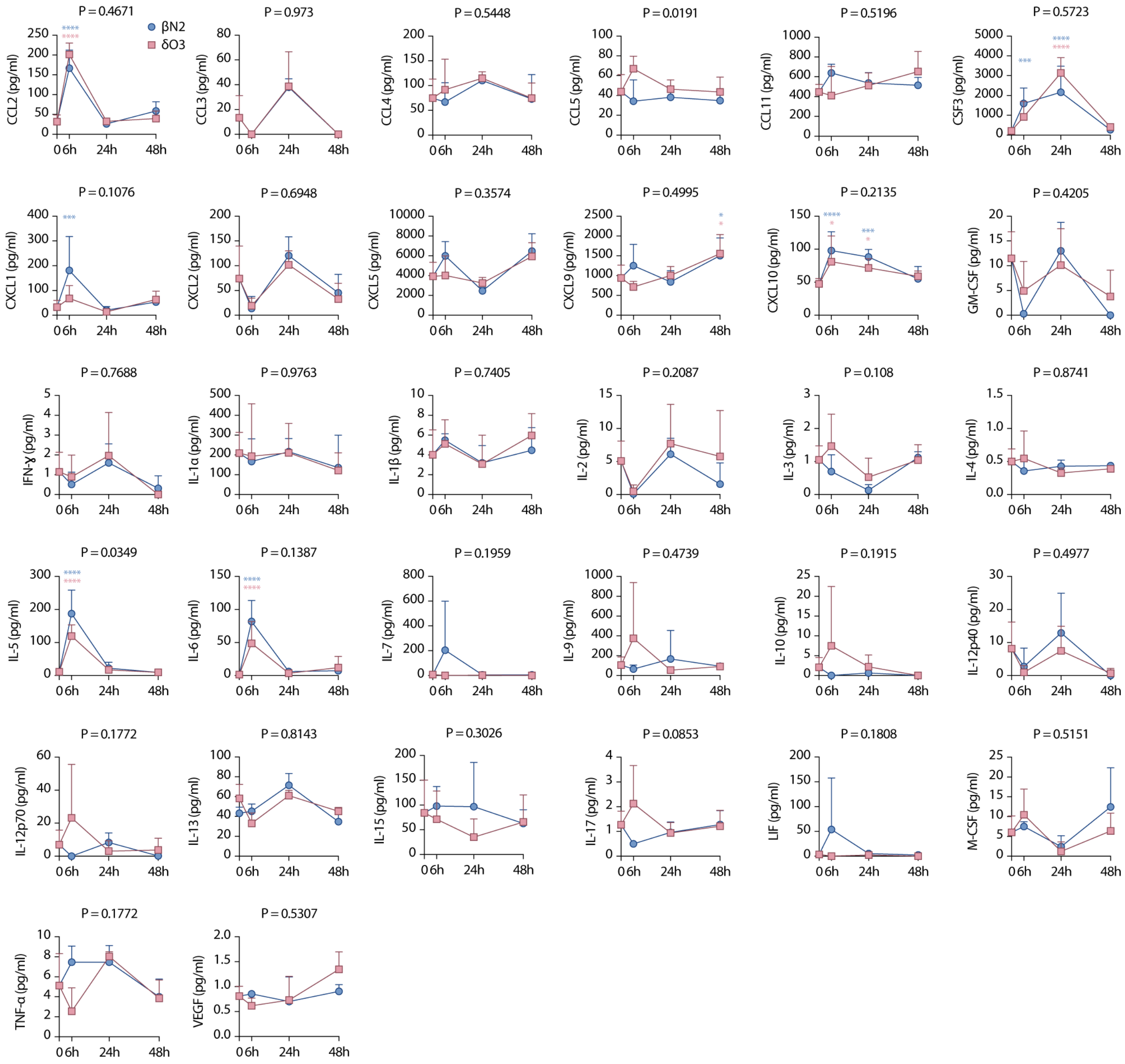


**Figure S39. Cytokine profile to empty-LNPs**. BALB/c mice were injected intramuscularly with empty-LNPs containing βN2 or δO3. Cytokines were measured in serum 6, 24, and 48 hours by Luminex assay. Data for βN2 are indicated by blue circles, and data for δO3 are indicated by red squares. Data and error bars indicate the mean and standard deviation across samples from independent mice (n = 4, terminal timepoints). P-values were determined by two-way ANOVA with Bonferroni's multiple comparisons test. P values above graphs represent the ionizable lipid factor. P values above data points represent the comparison with baseline.

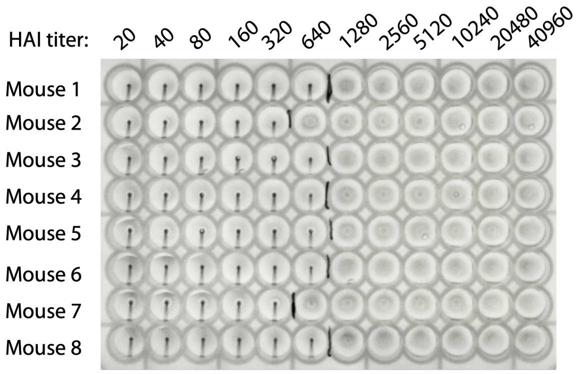


**Figure S40. Representative image of an HAI assay plate**

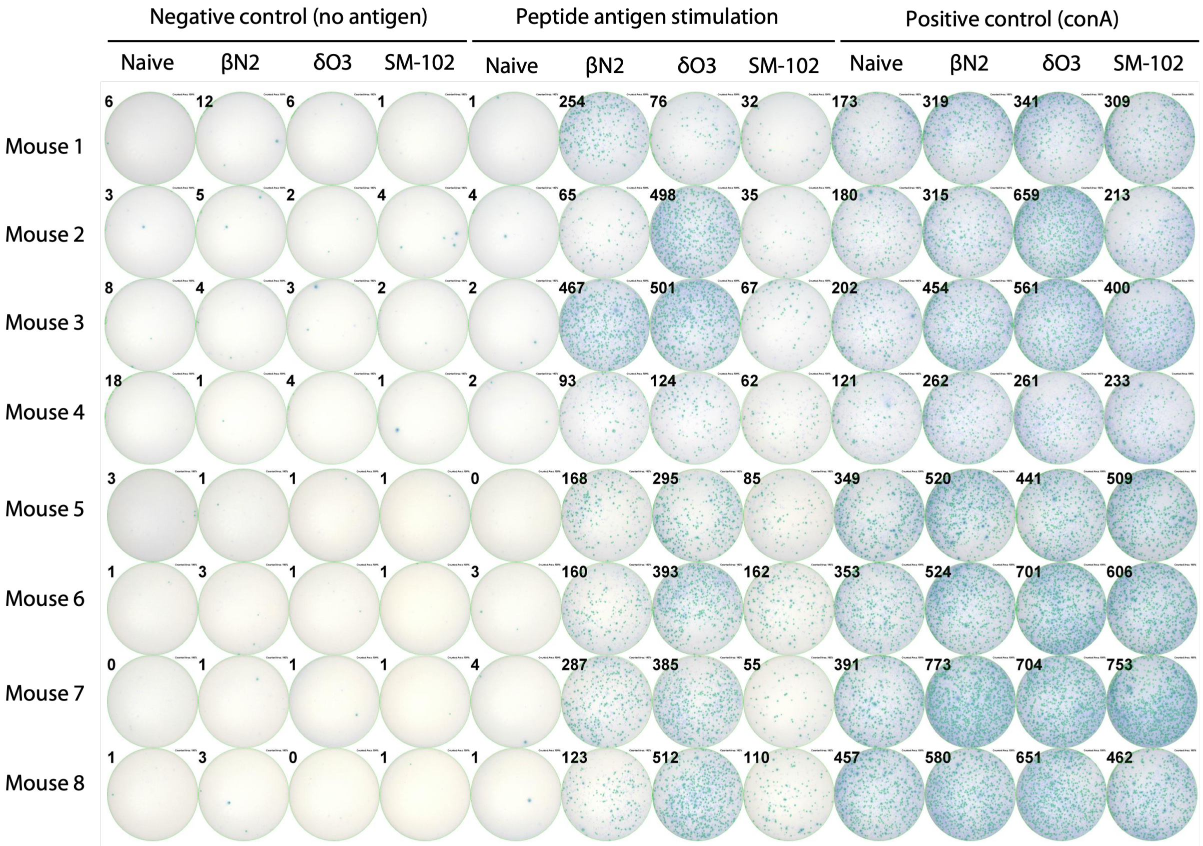


**Figure S41. IFN-ɣ T cell ELISPOT assay wells with automated counts.**

### Supplementary Tables

#### Table S1. mRNA sequences

| Encoded protein | Length | mRNA sequence (5’ to 3’) |
|---|---|---|
| Firefly luciferase (FLuc) | 1865 nt | AGGGAGACUGCCACCAUGGAGGACGCCAAGAACAUCAAGAAGGGCCCCGC CCCCUUCUACCCCCUGGAGGACGGCACCGCCGGCGAGCAGCUGCACAAGG CCAUGAAGCGGUACGCCCUGGUGCCCGGCACCAUCGCCUUCACCGACGCC CACAUCGAGGUGGACAUCACCUACGCCGAGUACUUCGAGAUGAGCGUGCG GCUGGCCGAGGCCAUGAAGCGGUACGGCCUGAACACCAACCACCGGAUCG UGGUGUGCAGCGAGAACAGCCUGCAGUUCUUCAUGCCCGUGCUGGGCGC CCUGUUCAUCGGCGUGGCCGUGGCCCCCGCCAACGACAUCUACAACGAGC GGGAGCUGCUGAACAGCAUGGGCAUCAGCCAGCCCACCGUGGUGUUCGU GAGCAAGAAGGGCCUGCAGAAGAUCCUGAACGUGCAGAAGAAGCUGCCCA UCAUCCAGAAGAUCAUCAUCAUGGACAGCAAGACCGACUACCAGGGCUUC CAGAGCAUGUACACCUUCGUGACCAGCCACCUGCCCCCCGGCUUCAACGA GUACGACUUCGUGCCCGAGAGCUUCGACCGGGACAAGACCAUCGCCCUGA UCAUGAACAGCAGCGGCAGCACCGGCCUGCCCAAGGGCGUGGCCCUGCC CCACCGGACCGCCUGCGUGCGGUUCAGCCACGCCCGGGACCCCAUCUUC GGCAACCAGAUCAUCCCCGACACCGCCAUCCUGAGCGUGGUGCCCUUCCA CCACGGCUUCGGCAUGUUCACCACCCUGGGCUACCUGAUCUGCGGCUUCC GGGUGGUGCUGAUGUACCGGUUCGAGGAGGAGCUGUUCCUGCGGAGCCU GCAGGACUACAAGAUCCAGAGCGCCCUGCUGGUGCCCACCCUGUUCAGCU UCUUCGCCAAGAGCACCCUGAUCGACAAGUACGACCUGAGCAACCUGCAC GAGAUCGCCAGCGGCGGCGCCCCCCUGAGCAAGGAGGUGGGCGAGGCCG UGGCCAAGCGGUUCCACCUGCCCGGCAUCCGGCAGGGCUACGGCCUGAC CGAGACCACCAGCGCCAUCCUGAUCACCCCCGAGGGCGACGACAAGCCCG GCGCCGUGGGCAAGGUGGUGCCCUUCUUCGAGGCCAAGGUGGUGGACCU GGACACCGGCAAGACCCUGGGCGUGAACCAGCGGGGCGAGCUGUGCGUG CGGGGCCCCAUGAUCAUGAGCGGCUACGUGAACAACCCCGAGGCCACCAA CGCCCUGAUCGACAAGGACGGCUGGCUGCACAGCGGCGACAUCGCCUACU GGGACGAGGACGAGCACUUCUUCAUCGUGGACCGGCUcAAGAGCCUGAUC AAGUACAAGGGCUACCAGGUGGCCCCCGCCGAGCUGGAGAGCAUCCUGCU GCAGCACCCCAACAUCUUCGACGCCGGCGUGGCCGGCCUGCCCGACGAC GACGCCGGCGAGCUGCCCGCCGCCGUGGUGGUGCUGGAGCACGGCAAGA CCAUGACCGAGAAGGAGAUCGUGGACUACGUGGCCAGCCAGGUGACCACC GCCAAGAAGCUGCGGGGCGGCGUGGUGUUCGUGGACGAGGUGCCCAAGG GCCUGACCGGCAAGCUGGACGCCCGGAAGAUCCGGGAGAUCCUGAUCAAG GCCAAGAAGGGCGGCAAGAUCGCCGUGUGAUAAGCUGCCUUCUGCGGGG CUUGCCUUCUGGCCAUGCCCUUCUUCUCUCCCUUGCACCUGUACCUCUUG GUCUUUGAAUAAAGCCUGAGUAGGAAGAAAAAAAAAAAAAAAAAAAAAAAAA AAAAAAAAAAAAAAAAAAAAAAAAAAAAAAAAAAAAAAAAAAAAAAAAAAAAAA AAAAAAAAAAAAAAAAAAAAAA |
| Human erythropoietin (hEPO) | 794 nt | AGGGAGACUGCCACCAUGGGCGUGCACGAGUGCCCCGCCUGGCUGUGGC UGCUGCUGAGCCUGCUGAGCCUGCCCCUGGGCCUGCCCGUGCUGGGCGC CCCCCCCCGGCUGAUCUGCGACAGCCGGGUGCUGGAGCGGUACCUGCUG GAGGCCAAGGAGGCCGAGAACAUCACCACCGGCUGCGCCGAGCACUGCAG CCUGAACGAGAACAUCACCGUGCCCGACACCAAGGUGAACUUCUACGCCU GGAAGCGGAUGGAGGUGGGCCAGCAGGCCGUGGAGGUGUGGCAGGGCCU GGCCCUGCUGAGCGAGGCCGUGCUGCGGGGCCAGGCCCUGCUGGUGAAC AGCAGCCAGCCCUGGGAGCCCCUGCAGCUGCACGUGGACAAGGCCGUGA GCGGCCUGCGGAGCCUGACCACCCUGCUGCGGGCCCUGGGCGCCCAGAA GGAGGCCAUCAGCCCCCCCGACGCCGCCAGCGCCGCCCCCCUGCGGACC AUCACCGCCGACACCUUCCGGAAGCUGUUCCGGGUGUACAGCAACUUCCU GCGGGGCAAGCUGAAGCUGUACACCGGCGAGGCCUGCCGGACCGGCGAC CGGUGAUAAGCUGCCUUCUGCGGGGCUUGCCUUCUGGCCAUGCCCUUCU UCUCUCCCUUGCACCUGUACCUCUUGGUCUUUGAAUAAAGCCUGAGUAGG AAGAAAAAAAAAAAAAAAAAAAAAAAAAAAAAAAAAAAAAAAAAAAAAAAAAA AAAAAAAAAAAAAAAAAAAAAAAAAAAAAAAAAAAAAAAAAAAAAAAAAA |

| | | |
|---|---|---|
| Hemagglutinin of influenza A H1N1 strain A/PR/8/34 (HA) | 1911 nt | AGGGAGACUGCCACCAUGAAGGCCAACCUGCUGGUGCUGCUGUGCGCCC<br>UGGCCGCCGCCGACGCCGACACCAUCUGCAUCGGCUACCACGCCAACAAC<br>AGCACCGACACCGUGGACACCGUGCUGGAGAAGAACGUGACCGUGACCCA<br>CAGCGUGAACCUGCUGGAGGACAGCCACAACGGCAAGCUGUGCCGGCUGA<br>AGGGCAUCGCCCCCCUGCAGCUGGGCAAGUGCAACAUCGCCGGCUGGCU<br>GCUGGGCAACCCCGAGUGCGACCCCCUGCUGCCCGUGCGGAGCUGGAGC<br>UACAUCGUGGAGACCCCCAACAGCGAGAACGGCAUCUGCUACCCCGGCGA<br>CUUCAUCGACUACGAGGAGCUGCGGGAGCAGCUGAGCAGCGUGAGCAGC<br>UUCGAGCGGUUCGAGAUCUUCCCCAAGGAGAGCAGCUGGCCCAACCACAA<br>CACCAACGGCGUGACCGCCGCCUGCAGCCACGAGGGCAAGAGCAGCUUCU<br>ACCGGAACCUGCUGUGGCUGACCGAGAAGGAGGGCAGCUACCCCAAGCUG<br>AAGAACAGCUACGUGAACAAGAAGGGCAAGGAGGUGCUGGUGCUGUGGGG<br>CAUCCACCACCCCAGCAACAGCAAGGAGCAGCAGAACCUGUACCAGAACGA<br>GAACGCCUACGUGAGCGUGGUGACCAGCAACUACAACCGGCGGUUCACCC<br>CCGAGAUCGCCGAGCGGCCCAAGGUGCGGGACCAGGCCGGCCGGAUGAA<br>CUACUACUGGACCCUGCUGAAGCCCGGCGACACCAUCAUCUUCGAGGCCA<br>ACGGCAACCUGAUCGCCCCCAUGUACGCCUUCGCCCUGAGCCGGGGCUUC<br>GGCAGCGGCAUCAUCACCAGCAACGCCAGCAUGCACGAGUGCAACACCAA<br>GUGCCAGACCCCCCUGGGCGCCAUCAACAGCAGCCUGCCCUACCAGAACA<br>UCCACCCCGUGACCAUCGGCGAGUGCCCCAAGUACGUGCGGAGCGCCAAG<br>CUGCGGAUGGUGACCGGCCUGCGGAACAUCCCCAGCAUCCAGAGCCGGG<br>GCCUGUUCGGCGCCAUCGCCGGCUUCAUCGAGGGCGGCUGGACCGGCAU<br>GAUCGACGGCUGGUACGGCUACCACCACCAGAACGAGCAGGGCAGCGGCU<br>ACGCCGCCGACCAGAAGAGCACCCAGAACGCCAUCAACGGCAUCACCAACA<br>AGGUGAACACCGUGAUCGAGAAGAUGAACAUCCAGUUCACCGCCGUGGGC<br>AAGGAGUUCAACAAGCUGGAGAAGCGGAUGGAGAACCUGAACAAGAAGGU<br>GGACGACGGCUUCCUGGACAUCUGGACCUACAACGCCGAGCUGCUGGUG<br>CUGCUGGAGAACGAGCGGACCCUGGACUUCCACGACAGCAACAUGAAGAA<br>CCUGUACGAGAAGGUGAAGAGCCAGCUGAAGAACAACGCCAAGGAGAUCG<br>GCAACGGCUGCUUCGAGUUCUACCACAAGUGCGACAACGAGUGCAUGGAG<br>AGCGUGCGGAACGGCACCUACGACUACCCCAAGUACAGCGAGGAGAGCAA<br>GCUGAACCGGGAGAAGGUGGACGGCGUGAAGCUGGAGAGCAUGGGCAUC<br>UACCAGAUCCUGGCCAUCUACAGCACCGUGGCCAGCAGCCUGGUGCUGCU<br>GGUGAGCCUGGGCGCCAUCAGCUUCUGGAUGUGCAGCAACGGCAGCCUG<br>CAGUGCCGGAUCUGCAUCUGAUAAGCUGCCUUCUGCGGGGCUUGCCUUC<br>UGGCCAUGCCCUUCUUCUCUCCCUUGCACCUGUACCUCUUGGUCUUUGAA<br>UAAAGCCUGAGUAGGAAGAAAAAAAAAAAAAAAAAAAAAAAAAAAAAAAAAAAAA<br>AAAAAAAAAAAAAAAAAAAAAAAAAAAAAAAAAAAAAAAAAAAAAAAAAAAAAAA<br>AAAAAAAAAAAAU |

**Table S2. LNP characterization before freezing.**

| Ionizable Lipid | mRNA | Size (nm) | PDI | Zeta Potential (mV) | Zeta Potential Deviation (mV) | mRNA Encapsulation (%) | Apparent $pK_a$ |
|---|---|---|---|---|---|---|---|
| βO2 | FLuc | 110.8 | 0.067 | -5.0 | 10.2 | 77.2 | 5.5 |
| βO3 | FLuc | 119.5 | 0.078 | -4.7 | 8.5 | 80.5 | 5.9 |
| γO2 | FLuc | 89.3 | 0.126 | -2.4 | 9.4 | 92.7 | 5.8 |
| γO3 | FLuc | 108.7 | 0.080 | -3.5 | 8.1 | 86.7 | 6.2 |
| γO4 | FLuc | 116.8 | 0.087 | -2.5 | 8.6 | 90.7 | 6.3 |
| δO3 | FLuc | 100.5 | 0.098 | -2.6 | 11.0 | 93.5 | 6.4 |
| δO4 | FLuc | 105.1 | 0.174 | -0.5 | 5.1 | 96.0 | 6.6 |
| ALC-0315 | FLuc | 86.1 | 0.190 | -6.6 | 10.7 | 91.8 | 6.1 |
| MC3 | FLuc | 75.5 | 0.108 | -5.0 | 12.7 | 93.0 | 6.4 |
| βN2 | FLuc | 110.1 | 0.104 | -4.4 | 4.0 | 90.9 | 6.5 |
| SM-102 | FLuc | 107.0 | 0.107 | -4.7 | 7.9 | 89.8 | 6.5 |
| δO3 | hEPO | 100.3 | 0.106 | -4.7 | 8.4 | 93.4 | nd[a] |
| βN2 | hEPO | 105.3 | 0.075 | -4.0 | 7.1 | 93.8 | nd |
| δO3 | no mRNA | 67.6 | 0.081 | -2.4 | 12.3 | na[b] | na |
| βN2 | no mRNA | 60.9 | 0.131 | 2.5 | 13.4 | na | na |
| βN2 | HA | 113.0 | 0.089 | -11.1 | 9.1 | 90.4 | nd |
| δO3 | HA | 112.7 | 0.039 | -4.7 | 8.6 | 92.6 | nd |
| SM-102 | HA | 78.9 | 0.054 | -7.0 | 9.4 | 95.0 | nd |

[a] nd denotes not determined
[b] na denotes not applicable

**Table S3. LNP characterization after one freeze-thaw cycle.**

| Ionizable Lipid | mRNA | Size (nm) | PDI | Zeta Potential (mV) | Zeta Potential Deviation (mV) | mRNA Encapsulation (%) |
|---|---|---|---|---|---|---|
| βO2 | FLuc | 113.6 | 0.112 | -5.0 | 5.3 | 72.5 |
| βO3 | FLuc | 122.1 | 0.077 | -4.5 | 6.9 | 76.4 |
| γO2 | FLuc | 95.5 | 0.134 | -6.9 | 5.5 | 90.4 |
| γO3 | FLuc | 110.7 | 0.083 | -5.1 | 5.7 | 85.7 |
| γO4 | FLuc | 120.8 | 0.075 | -5.9 | 6.6 | 87.3 |
| δO3 | FLuc | 105.5 | 0.102 | -7.6 | 7.4 | 91.1 |
| δO4 | FLuc | 232.1 | 0.392 | -4.1 | 4.0 | 72.7 |
| βN2 | FLuc | 116.4 | 0.095 | -13.8 | 10.8 | 86.71 |
| ALC-0315 | FLuc | 107.7 | 0.224 | -6.8 | 5.9 | 85.2 |
| MC3 | FLuc | 109.2 | 0.329 | -13.5 | 7.2 | 86.9 |
| SM-102 | FLuc | 109.5 | 0.090 | -8.0 | 7.8 | 85.79 |

**Table S4. Summary of one-way ANOVA for total luminescent flux in injected hindlimbs at 6 hours**

| LNP comparison | Mean difference | Percent difference | Adjusted P value |
|---|---|---|---|
| βN2 vs. Saline | 8.87E+08 | 100% | <0.0001 |
| βN2 vs. MC3 | 7.53E+08 | 85% | <0.0001 |
| βN2 vs. ALC-0315 | 4.39E+08 | 49% | 0.0364 |
| βN2 vs. SM-102 | -2.23E+07 | -3% | >0.9999 |
| βN2 vs. βO2 | 8.85E+08 | 100% | <0.0001 |
| βN2 vs. βO3 | 7.83E+08 | 88% | <0.0001 |
| βN2 vs. γO2 | 7.77E+08 | 88% | <0.0001 |
| βN2 vs. γO3 | 6.70E+08 | 76% | 0.0002 |
| βN2 vs. γO4 | 4.73E+08 | 53% | 0.0174 |
| βN2 vs. δO3 | -6.48E+08 | -73% | 0.0003 |
| βN2 vs. δO4 | 4.73E+08 | 53% | 0.0176 |
| δO3 vs. Saline | 1.54E+09 | 100% | <0.0001 |
| δO3 vs. MC3 | 1.40E+09 | 91% | <0.0001 |
| δO3 vs. ALC-0315 | 1.09E+09 | 71% | <0.0001 |
| δO3 vs. SM-102 | 6.26E+08 | 41% | 0.0005 |
| δO3 vs. βO2 | 1.53E+09 | 100% | <0.0001 |
| δO3 vs. βO3 | 1.43E+09 | 93% | <0.0001 |
| δO3 vs. γO2 | 1.43E+09 | 93% | <0.0001 |
| δO3 vs. γO3 | 1.32E+09 | 86% | <0.0001 |
| δO3 vs. γO4 | 1.12E+09 | 73% | <0.0001 |
| δO3 vs. δO4 | 1.12E+09 | 73% | <0.0001 |

**Table S5. Summary of mean pharmacokinetic parameters for βN2.**

| Parameter | Parameter estimate | | | |
|---|---|---|---|---|
| | Plasma | Muscle | Liver | Spleen |
| $t_{max}$ (h) | 6.00 | 3.00 | 48 &168[a] | 504 |
| $C_{max}$ (ng/mL or ng/g) | 348 | 12023 | 1890 &1900[a] | 909 |
| Apparent $t_{1/2}$ (h) | 295[b] | 362[b] | nc[c] | nc |
| $AUC_{0\text{-tlast}}$ (h*ng/mL or h*ng/g) | 13772 | 3243535 | 693954 | 391429 |
| Tissue/plasma AUC ratio | na[d] | 236 | 50.4 | 28.4 |

[a] Two peaks were observed at 48 and 168 h post-dose with similar $C_{max}$ values.
[b] Apparent $t_{1/2}$ estimate is not accurate as the sampling interval during the terminal phase is <2 x $t_{1/2}$.
[c] nc denotes not calculable as the terminal phase is not well defined ($r^2$<0.8).
[d] na denotes not applicable

**Table S6. Summary of mean pharmacokinetic parameters for δO3.**

| Parameter | Parameter estimate | | | |
|---|---|---|---|---|
| | Plasma | Muscle | Liver | Spleen |
| $t_{max}$ (h) | 3.00 | 3.00 | 12.0 | 3.00 |
| $C_{max}$ (ng/mL or ng/g) | 108 | 13500 | 625 | 1173 |
| Apparent $t_{1/2}$ (h) | 15.3 | nc[a] | nc | nc |
| $AUC_{0\text{-tlast}}$ (h*ng/mL or h*ng/g) | 1033 | 994206 | 46954 | 61096 |
| Tissue/plasma AUC ratio | na[b] | 962 | 45.5 | 59.1 |

[a] nc denotes not calculable as the terminal phase is not well defined ($r^2$<0.8).
[b] na denotes not applicable

**Table S7. Lipid dose in pooled feces and urine**

| Experimental time (h) | Parent lipid in feces (% injected dose) | | Parent lipid in urine (ng/mL) | |
|---|---|---|---|---|
| | βN2 | δO3 | βN2 | δO3 |
| 0-3 | 0.0596 | 0.118 | BLQ | No Peak |
| 0-6 | 1.58 | 0.311 | BLQ | No Peak |
| 0-12 | 1.32 | 1.51 | BLQ | No Peak |
| 0-24 | 5.02 | 0.811 | BLQ | No Peak |
| 0-48 | 1.04 | 1.88 | BLQ | No Peak |
| 0-96 | 4.25 | 3.35 | BLQ | No Peak |
| 0-168 | 2.46 | 0.284 | nm[b] | nm |
| 0-504 (21 days) | 17.8 | nm | nm | nm |

[a] BLQ denotes below the lower limit of quantification (0.5 ng/mL).
[b] nm denotes not measured

**Table S8. Cytokine profile of mRNA-LNPs ($AUC_{0\text{-}48h}$)**

| Cytokines | βN2 | | | δO3 | | |
|---|---|---|---|---|---|---|
| | Total Area | Std. Error | 95% CI | Total Area | Std. Error | 95% CI |
| IL-6 | 1598 | 523.9 | 570.9 to 2625 | 1092 | 366.2 | 374.5 to 1810 |
| CSF3 | 130762 | 34567 | 63012 to 198512 | 51981 | 12744 | 27003 to 76958 |
| CXCL1 | 4731 | 2171 | 475.5 to 8987 | 3329 | 1515 | 358.9 to 6299 |
| CCL2 | 5827 | 1144 | 3584 to 8070 | 4654 | 746.7 | 3190 to 6117 |
| IL-5 | 1119 | 249.9 | 629.0 to 1609 | 1427 | 339.7 | 760.7 to 2092 |
| CXCL10 | 6418 | 1001 | 4457 to 8380 | 4036 | 1032 | 2013 to 6059 |

**Table S9. Cytokine profile of empty-LNPs ($AUC_{0\text{-}48h}$)**

| Cytokines | βN2 | | | δO3 | | |
|---|---|---|---|---|---|---|
| | Total Area | Std. Error | 95% CI | Total Area | Std. Error | 95% CI |
| IL-6 | 1205 | 321.6 | 574.7 to 1835 | 814.5 | 373.8 | 81.75 to 1547 |
| CSF3 | 68723 | 21038 | 27490 to 109956 | 82693 | 13804 | 55638 to 109747 |
| CXCL1 | 1981 | 670.3 | 667.4 to 3295 | 3353 | 1337 | 733.0 to 5974 |
| CCL2 | 3683 | 371.3 | 2955 to 4411 | 3357 | 527.3 | 2323 to 4390 |
| IL-5 | 2874 | 725.4 | 1453 to 4296 | 1958 | 335.7 | 1300 to 2616 |
| CXCL10 | 3839 | 392.7 | 3070 to 4609 | 3332 | 438.1 | 2473 to 4191 |
